# Predicting tumour evolution and drug resistance from heterogenous longitudinal cancer data

**DOI:** 10.1101/2024.12.30.630757

**Authors:** Giovanni Santacatterina, Riccardo Bergamin, Antonella Zucchetto, Roberta Laureana, Maria Ilaria del Principe, Valter Gattei, Leonardo Egidi, Giulio Caravagna

**Author notes:** These authors contributed equally to this work (Leonardo Egidi, Giulio Caravagna). **Corresponding author:** Leonardo Egidi and Giulio Caravagna *E-mail address:* and.

## Abstract

The kinetic parameters of cancer population dynamics are critical for developing reliable predictors of tumour growth patterns, extracting metrics for patient stratification and creating algorithms that can forecast clinically significant events. Here, we introduce a model-based Bayesian framework that leverages longitudinal phenotypic (e.g., tumour volume, cell counts) or genotypic (e.g., mutation frequency) data to infer critical parameters of tumour progression within a single patient. Our models uses population genetics to estimate probability distributions for tumour growth rates, initiation and extinction times, pinpointing abrupt shifts in tumour dynamics due to treatment response and revealing associations between drug resistance and pre-existing cancer cell populations. We apply our framework to address pivotal clinical questions across three major cancer types. In colorectal cancer, we use tumour markers data to identify extensive pre-existing RAS-linked resistance to cetuximab. In lung cancer, we use somatic mutation frequencies in citculating tumour DNA to determine prognostic growth rates and develop a test for monitoring minimal residual disease. In chronic lymphocytic leukaemia, we use white blood cell counts to stratify patients by growth patterns and predict time to treatment, advancing adaptive monitoring strategies.

## 1. Introduction

Cancer is a disease based on the dysregulation of standard cellular growth, where malignant cells proliferate at excessive rates, out-competing normal cells and eventually disrupting tissue homeostasis^1^. The drivers of this process are genetic and epigenetic alterations that accrue in distinct cancer clones^2^, the fundamental elements that rule tumour evolution^3,4^. The cancer genomics community has invested much in the development of sequencing and analysis technologies to determine, from single tumour snapshots (i.e., observations at a single time point) the patterns of such growth^5–12^. High-resolution sequencing data is the best readout to determine cancer clonal dynamics^13^. Still, costs limit the number of data points available to, usually, one pre-treatment and, only in some cases, one post-treatment or metastatic sample. However, modern sampling and sequencing approaches are making longitudinal data increasingly available. For instance, in the wet lab, rich time series are available through low-resolution measurements such as cell counts or tumour mass assays, often generated from experimental models like patient-derived organoids or xenografts^14^. These assays are particularly useful for conducting large-scale drug screening in experimentally controlled environments^15^. In the clinical setting, similar measurements are obtained using various non-invasive biopsy technologies. These include liquid biopsies^16^, which can assess molecular markers (e.g., antigen-based markers or others), circulating tumour DNA and minimal residual disease for monitoring^17^. Additionally, tissue-level assays based on imaging techniques (e.g., computed tomography or others^18^) measure phenotypes that correlate with tumour growth.

Despite the experimental development of these technologies, an open machine learning problem is how to use these measurements to parameterise cancer dynamics that are latent, noisy, and can follow distinct mathematical patterns^19^, a fundamental bottleneck to transform these data into useful clinical predictors. The ground to frame this problem is population genetics, the branch of mathematics studying alleles diffusing in expanding populations^20,21^. In cancer, however, growth rates are not constant because patients undergo treatments, which are interventions that change growth rates and affect latent tumour dynamics. Moreover, tumour often relapses to treatment, leading to waves of shrink and re-growth which can be associated with distinct growth parameters in the clinical history of a patient^22^. While treatment initiation events are typically well-documented in clinical records, the exact timing of tumor regrowth is often less clear. This uncertainty highlights the need to infer these rate-changing events from available data when they are not explicitly recorded. Finally, multiple competing populations also challenge decoupling the contribution of the individual components to the tumour growth, even because measurements report the total tumour burden and not the components^23,24^. These features make the inferential problem challenging and, to the best of our understanding, a general framework to approach these problems is missing.

In this work, we develop a general framework that uses Bayesian inference^25,26^ to resist noise in the data and infer tumour evolutionary parameters from various longitudinal measurements, including tumour volume data, tumour-associated cell counts or markers, and somatic mutation frequencies in circulating tumour DNA (ctDNA). We use our approach to solve three clinical questions at the cornerstone of cancer precision medicine. First, we understand treatment resistance patterns in colon cancer patients, using tumour markers to reveal extensive pre-existing resistance to treatment with a monoclonal antibody. Second, we predict prognosis from tumour evolution of lung cancer patients, using ctDNA to derive growth rates and a novel test for minimal residual disease monitoring. Third, we predict outcome from tumour evolution of chronic leukaemia patients, using white-blood count data to predict which and when patients will go to treatment. Our work resolves a critical missing gap in an evolving field, offering a solid ground for researchers and clinicians who wish to turn longitudinal tumour evolution patterns into translational predictors.

## 2. Results

### 2.1. Bayesian inference of population dynamics (biPOD)

We developed a framework to perform *Bayesian Inference of Population Dynamics* (biPOD) for multiple types of regression of tumour abundance against time, using the principles of population genetics^27^ (Fig.1). All our models are supervised, and use *N* measurements ***X*** = {(*x*_*i*_, *y*_*i*_) ∣ *i* = 1, …, *N*} of tumour size *y*_*i*_ at time *x*_*i*_ to infer some tumour parameters **Θ**. The mathematical details and derivations are given in Supplementary Material C.

**Figure 1.**
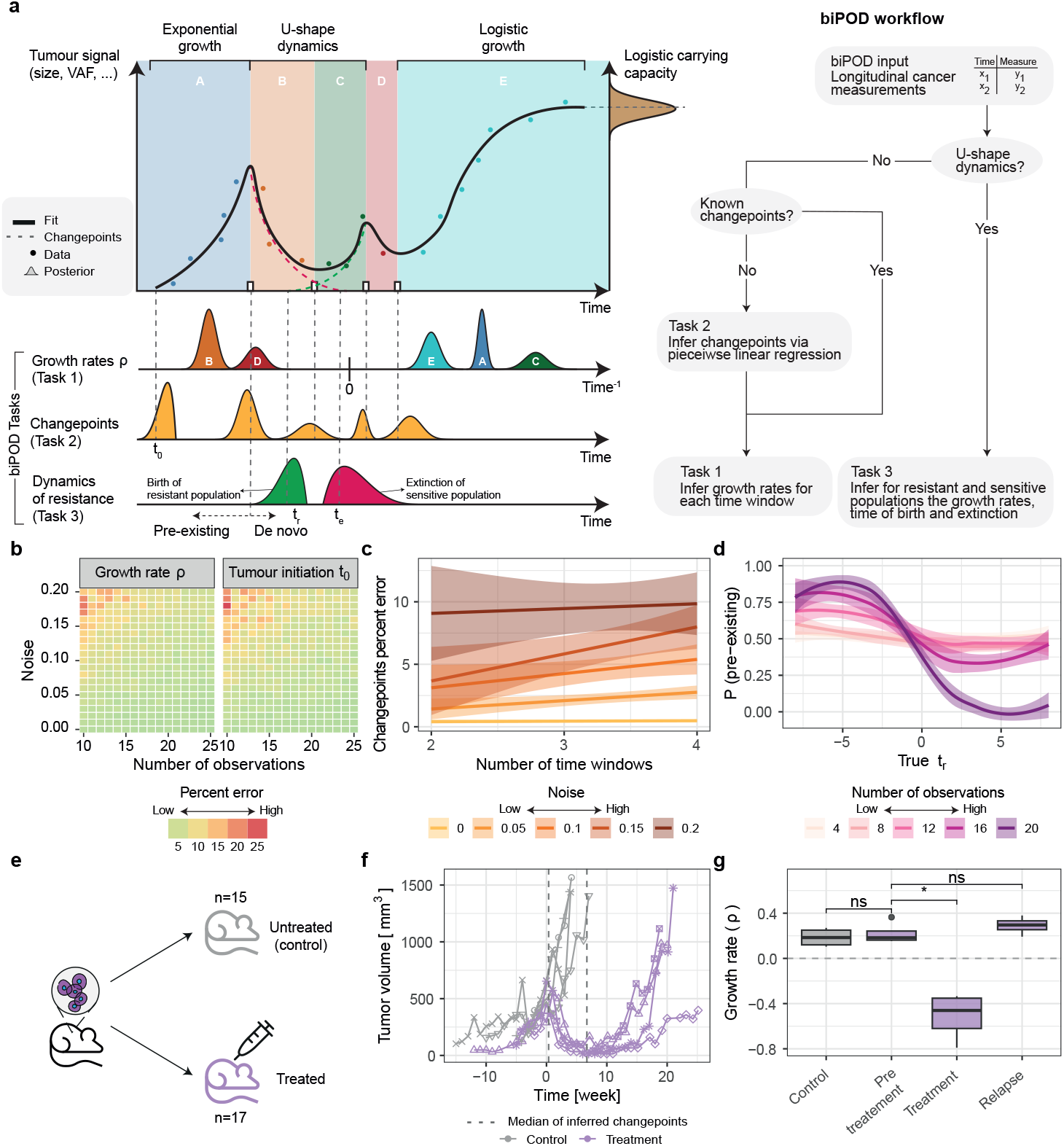
**a**. biPOD is a Bayesian inference framework to infer kinetic tumour parameters from longitudinal (*x*_*i*_, *y*_*i*_) phenotypic (e.g., tumour mass, cell counts) and genotypic (e.g., Variant Allele Frequency, VAF) data. (Task1) From known time windows (e.g., between treatments), biPOD considers exponential and logistic tumour growth mode to infer probabilistic tumour growth rate parameters *ρ* and a tumour initiation time *t*_0_. (Task 2) biPOD can automatically detect instant of times *t*_∗_ when tumour parameters change abruptly. (Task 3) biPOD can model dynamics of resistance assuming the presence of two populations as well as the likelihood of pre-existing resistance when tumours relapse after shrinking (*t*_*s*_ < *t*_1_). **b, c, d** biPOD performance with synthetic tests considering Gaussian noise and variable number of observations. Error (percentages) in growth rates and tumour initiation time estimates (panel b). Trend errors in automatically detecting when tumour growth parameters change (panel c). Probability of correctly classifying a resistant population as pre-existing (panel d). **e**. Validation of biPOD through patient-derived xenograft dynamics in a drug screening assay^14^ (*n* = 15 control group, *n* = 17 group treated with carboplatin). **f**. Weekly tumour volumes used by biPOD to detect two changes in tumour dynamics in the treatment group. biPOD detects a first changepoint perfectly matching the recorded data of treatment initiation (*t* = 0), and a second changepoint that explain the beginning of tumour re-growth. **g**. Tumour growth rates reflect distinct values for each time window identified by biPOD. Pre-treatment (baseline growth) dynamics are statistically equivalent among groups (*p* = 0.69). Treatment dynamics differ significantly from pre-treatment (*p* = 0.029), while relapse dynamics are statistically equivalent to pre-treatment (*p* = 0.34). P-values are computed using Wilcoxon test at *α* = 0.05 significance level.

In a first task, tumour dynamics span *J* ≥ 1 time windows determined by therapeutic interventions at known times *t*_*i*_ (Extended Fig.1a). biPOD can infer (Extended Fig.1b) the effective growth rate **Θ** = {*ρ*_*j*_ = *α*_*j*_ −*β*_*j*_ ∣ 1 ≤ *j* ≤ *J*} ∪{*t*_0_}, which depends on death (*β*_*j*_ > 0) and birth (*α*_*j*_ > 0) rates, together with a tumour-initiation time (*t*_0_ < *x*_1_), considering both exponential and logistic tumour growth models. For tumour size *y*(*x*) at time *x* > *t*_0_, under exponential growth the expected value for *y*(*x*) is a piece-wise exponential growth over *J* time windows

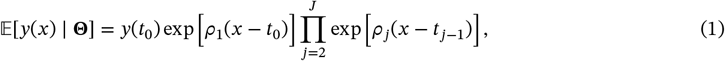

whereas under logistic growth the parameters **Θ** also include a carrying capacity *L* ≫ 0, and a recursive equation defines the expected size

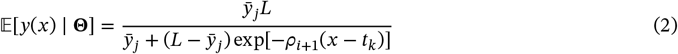

when 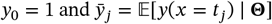. In terms of growth model selection, the best growth pattern given the data is chosen by using either Bayes factors^28^, or a mixture of models^29^ with competing components.

In a second task, biPOD can learn the change points *t*_*i*_ from ***X***, if these are unknown from data (Extended Fig.1c). Assuming *K* change-points and the logarithm of *y* as the outcome variable, a linear piece-wise regression for eq.(1) is

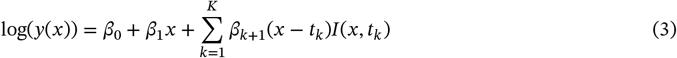

where *I*(*x, t*_*k*_) is an indicator function for *x* ≥ *t*_*k*_. This model can be solved as a system of linear equations to obtain the ***β*** coefficients that minimise the sum of squared residuals, yielding the optimal set of breakpoints. biPOD infers (Extended Fig.1d) the model parameters by combining differential evolution^30^ and leave-one-out cross-validation^31^.

In a third task, biPOD can also capture tumour shrinkage and re-growth dynamics, where a resistant population born at time *t*_*r*_ outgrows a sensitive population that goes extinct at time *t*_*s*_ (Extended Fig.1e). To do so, biPOD also models the time *t*_∗_ in which the resistant population is larger than the sensitive one, as well as the growth rates *ρ*_*s*_ and *ρ*_*r*_ for the populations. This inference (Extended Fig.1f) leverages the simplifying assumption that only two populations exist and the growth rates are constant. Nonetheless, in this scenario the model considers both pre-existing (*t*_*r*_ < *x*_1_) versus de novo resistance (*t*_*r*_ ≥ *x*_1_). Let *y*_*s*_(*x*) (resp *y*_*r*_(*x*)) be the number of sensitive (resp. resistant) cells at *x*, the equation that captures the total number of cells at any time *x* is

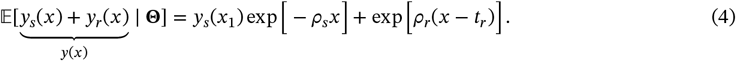

In this model, biPOD can estimate the size of the resistant population at the first timepoint *x*_1_ by computing *y*_*r*_(*x*_1_).

In all the settings, the model likelihood is Poisson with a rate equal to the expected population size, assuming the observations to be independent and identically distributed

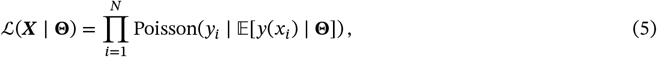

and the parameters **Θ** are inferred via Markov Chain Monte Carlo (MCMC)^32,33^ and/or variational inference^34–36^ using the Stan probabilistic programming language^37,38^. We remark that by developing biPOD as a Bayesian framework, uncertainty estimates of estimated parameters represent an intrinsic feature of the methodology.

### 2.2. Validation of biPOD

We validated biPOD with about 18000 synthetic simulations across the three tasks and with a controlled experiment with tumour volume data derived from a xenograft (Supplementary Material). In the simulations for the first task, even with various combinations of sample size and noise applied to the data, biPOD’s parameter estimates showed low percentile error even with noisy or few observations (MCMC error mean 5.32%, s.d. 3.79%, max 26.45%; Fig.1b), and with misspecified input change points (Extended Fig.2a). In these tests, the most challengins inferences were for small populations (error ≥ 30%; population order of magnitude ~ 10^1^). Even in those cases, however, the model correctly identified logistic versus exponential growth models (true positive rate, TPR, 0.809; false positive rate, FPR, 0; Extended Fig.2b,c). The variational implementation of biPOD achieved the same precision but was 100x faster than MCMC (mean 42.8x, s.d. 27.58x; Extended Fig.2d–f). biPOD performed very well also in the second and third inference tasks. Change-points were detected with mean errors below 5% even for high noise and multiple time windows (mean 4.58%, s.d. 4.92%; Fig. 1c), and pre-existing resistance was confidently identified as long as the process is observed for enough time, i.e., for ≥ 10 observations (54% accuracy with 4 observations, 91.3% with 20 observations). In the resistance setting a good accuracy was achieved even with just 12 observations (71%; Fig. 2d). As expected, datasets with many observations reduced errors (≥ 40%) for the parameters of the resistant population, while the sensitive population required fewer observations (mean 14.84%, s.d. 21.82%; Extended Fig.2g).

**Figure 2.**
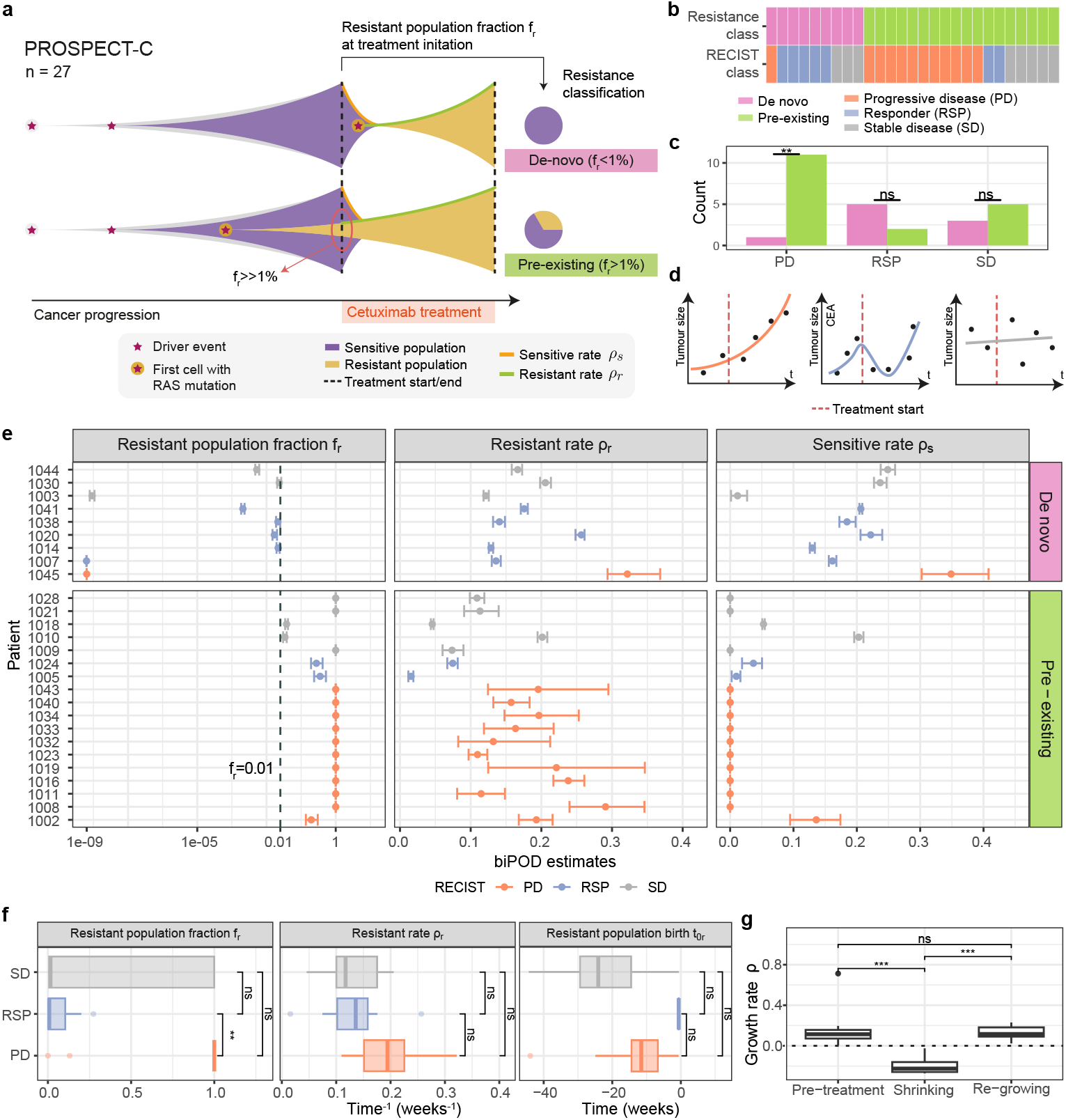
**a**. We used biPOD to determine pre-existing and de novo RAS-associated resistance patterns to cetuximab, in 27 patients from the PROSPECT-C phase II colorectal cancer clinical trial^44^ (whole cohort in Supplementary Fig.S6). Using growth parameters extracted from longitudinal carcinoembryonic antigen (CEA) data, we determined the fraction of pre-existing resistant cells at the time of treatment. **b**. Number of cases with more than 1% of pre-existing resistant cells at the time of treatment in the whole cohort and divided by RECIST classification^45^. **c**. Bar plots showing the number of pre-existing and de-novo cases in each RECIST classification category. In the progressive disease category, the majority of patients presents a pre-existing resistant clone. **d**. Visual representation of the RECIST classes: in the progressive disease case, the tumour grows, in the responder case, the tumour significantly shrinks, and in the stable disease case, the tumour exhibits minimal to no change. **e**. Bayesian credible intervals of response rates (*ρ*_*s*_), progression rates (*ρ*_*r*_), and fraction of the resistant population for each patient stratified by *de novo* vs *pre-existing*. **f**. Box plots showing fraction of the resistant population(*f*_*r*_), progression rates (*ρ*_*r*_), and instant of birth of the resistant clone (*t*_0,*r*_), stratified by RECIST classification categories. For the *t*_0,*r*_ parameter, only patients deemed as *pre-existing* are shown. P-values are computed using Wilcoxon test at *α* = 0.05 significance level. **g**. Boxplot showing net growth rate after, during, and before treatment in *n* = 7 patients that exhibited a clear diminuition of CEA right after the treatment. P-values are computed using Wilcoxon test at *α* = 0.05 significance level. P-values are computed using Wilcoxon test at *α* = 0.05 significance level.

We also validated biPOD against drug-response xenograft data (Fig. 1e), where tumour tissues were implanted into immunodeficient mice that received carboplatin, platinum-based chemotherapy at known times^14^. We used weekly tumour volume data to identify such times and infer tumour growth rate (*ρ*). In a typical case (Fig. 1f), biPOD was able to identify two change points with high precision: the first was clinically measured at *t* = 0, perfectly matching the treatment initiation (mean offset 4.22 days from treatment), while the second, though not clinically available, could explain the tumour relapse (mean offset 49.19 days from treatment). Importantly, the model did not detect any change point in four control samples, consistent with steady tumour dynamics (untreated mice; Supplementary Fig.S1). Overall, considering xenografts from three additional patients, biPOD identified change points correctly in 30 out of 32 mice (94% of total; Supplementary Figs.S1-S4). The estimated mean growth rate *ρ* was 0.187 weeks^−1^ (sd 0.081) in controls, and −0.522 weeks^−1^ (sd 0.230) in treated samples. Interestingly, relapse growth rates of treated mice were not significantly different (*p* = 0.34, Wilcoxon test) before or after treatment (mean *ρ* = 0.225, sd 0.808 before; mean *ρ* = 0.292, sd 0.823 after; Fig. 1g). This pattern is rather clear in some samples (Supplementary Fig.S1), where the posteriors over *ρ* are close. biPOD’s analysis therefore suggests no strong evidence for mutations conferring additional growth advantages (i.e., positive selection) to resist carboplatin. Instead, resistance may involve mechanisms enabling tumor cells to avoid negative selection, such as drug resistance, or neutral evolution of pre-existing resistant clones. In the case of carboplatin, resistance mechanisms might include enhanced DNA repair, such as increased ERCC1 expression^39,40^, altered drug uptake or efflux via transporters like p-glycoprotein^41^, detoxification through glutathione^42^, or defects in apoptotic pathways, such as TP53 mutations^43^. These mechanisms allow tumor survival under treatment pressure without altering growth rates, as observed with biPOD. Molecular analyses targeting these pathways could provide further insights into the basis of resistance and inform therapeutic strategies.

### 2.3. Characterising cetuximab resistance in a phase II colorectal cancer clinical trial

We used biPOD to study the dynamics of RAS-associated resistance to cetuximab, a monoclonal antibody used to treat metastatic cancers, in 27 patients from the PROSPECT-C phase II colorectal cancer clinical trial^44^. The response of each patient to treatment has been classified following the response evaluation criteria in solid tumors (RECIST^45^) criteria of progressing disease (PD), stable disease (SD), and responder (RSP). In the original paper, Khan *et al*.^44^ used differential equations modelling to estimate a pervasive pattern of pre-existing resistance in all non-responders. In this work, we use biPOD to (*i*) detect change points of tumour dynamics, (*ii*) confirm the pattern identified by Khan *et al*.^44^ and (*iii*) time the origin of the RAS-associated resistant clone (Fig.2a).

In some patients, serial measurements (23 median number of points, s.d. 10) of the carcinoembryonic antigen (CEA), a biomarker of tumour burden, showed characteristic U-shaped dynamics (example patient in Extended Fig. 3c,e,g). In this type of patient, biPOD successfully detected both the start and end of treatment (days 301 and 481; average offset 4.3 days for the patient in Extended Fig. 3c,e,g), as well as an additional change-point in between (day 355). Across the whole cohort, biPOD identified precisely (percentage errors below 15%) treatment change points (median offset days 17.60, s.d., 51.11; Extended Fig. 3a-b). Non-responders yielded slightly larger errors (percentage errors over 20% with a max of 63%) because their CEA dynamics were steady (Supplementary Fig.S5).

We classified states of pre-existing resistance if the fraction of resistant cells was exceeding 1% of the population size before treatment and found n=18 patients (66.6%) that had a pre-existing cetuximab-resistant population (Fig.2b and Supplementary Fig.S6). Interestingly, we found that 11 out of 12 patients with progressive disease had a pre-existing resistant clone, which suggests that the probability of a pre-existing clone is much higher than the one of a de novo clone in progressive disease (*p* = 0.0039, Chi-square test; Fig.2c-d). Across the cohort, we observed heterogeneous growth rates likely reflecting distinct clonal compositions (Fig.2e-f) – progression 0.02 ≤ *ρ* ≤ 0.30 weeks^−1^ (median *ρ* = 0.15); response 0 ≤ *ρ* ≤ 0.35 (median *ρ* = 0.07). Our results were qualitatively equivalent to^44^, but our response rates were generally lower, and the fraction of pre-existing resistant population was also smaller. This variability might be explained by the difference between the deterministic and stochastic model formulations, with our approach capturing noise at the expense of additional uncertainty in the predictions. Compared to^44^, however, our model could also determine the temporal origin (*t*_0,*r*_) of the RAS-mutant clone (Fig.2f). Across the RECIST groups, we found that the resistant population was born on average 13.6 weeks before treatment (95% confidence intervals: 1.86 to 39.2 weeks) for progressive disease, or 22.6 (95% confidence intervals: 1.98 to 42.8) for stable disease. These values represent an upper bound for estimating when the resistant population, responsible for at least one measurable unit of carcinoembryonic antigen (CEA), could be detected, highlighting its clinical relevance. This temporal offset offers a crucial window to scrutinise patients for RAS mutations before treatment. For instance, one could use biPOD’s distribution of the estimated clone sizes to design deep sequencing tests (with sufficient specificity) that search for RAS mutations in the pre-treatment sample and avoid delivering cetuximab to patients with adequate evidence for pre-existing RAS-mutant clones.

With our model, we determined growth rates (*ρ*) for all patients (Supplementary Fig.S5). In *n* = 7 patients that exhibited a clear diminuition of CEA right after the treatment, we found that the net growth rate before treatment and the the one when the CEA started to grow again were not significantly different (*p* = 0.71, Wilcoxon test) while the growth rate right after treatment was significantly lower (*p* = 5.8 × 10^−4^). One of the patients in this group is shown in Extended Fig. 3c. In this case the model predicts a posterior estimate of *ρ* = 0.073 weeks^−1^ (s.d. 0.004) before treatment, *ρ* = −0.348 weeks^−1^ (s.d. 0.012) in response to cetuximab, and *ρ* = 0.231 weeks^−1^ (s.d.0.004) at relapse. In this specific example, the difference among the growth rates before and after relapse (absolute difference 0.158 weeks^−1^) was striking. Given the leading role of RAS among colorectal cancer driver genes^46–48^, this might suggest that RAS mutations confer strong positive selection (percentage increase of about 200%) relative to the background genotype of the sensitive clone at least in some patients, but not in the whole cohort.

Nonetheless, biPOD’s parameters allowed us to predict the outcome of clonal dynamics among the RAS-mutant clone and the wildtype ancestor, even without the effect of treatment. For instance, assuming that the mutant clone is 1% of the sensitive one and using posterior means, the two clones would have the same size in about 8 weeks (almost 2 months), and in about one extra week, the mutant would have doubled the size of the sensitive clone. Moreover, by accessing a full posterior distribution over the parameters, these predictions can be extended to account for the standard deviations of the estimates. For example, with Gaussian posteriors, one can estimate 7.95 weeks (95% confidence intervals: 7.62 to 8.30) to match clone size and 9.15 weeks (95% confidence intervals: 8.76 to 9.55) to double the size of the mutant, using MCMC sampling. While these estimates ultimately depend on the initial size of the mutant clone, which is unknown, this analysis that quantitative biological hypothesis regarding resistance to treatment can be efficiently formulated using biPOD.

### 2.4. Patient stratification and minimal residual disease testing in TRACERx lung cancers

Tumour growth rate (biPOD’s parameter *ρ*) can become a robust patient stratification metric if subgroups of patients have distinct growth dynamics^51,52^. We could assess this from a large cohort of *n* = 101 early-stage non-small-cell lung cancer (NSCLC) which underwent surgical treatment and were available from the TRACERx (Tracking NSCLC Cancer Evolution through therapy) project^49,50^. This dataset included longitudinal variant allele frequencies (VAFs) for 200 patient-specific mutations derived from primary tumours (before surgery) and measured from circulating tumour DNA (ctDNA) collected after surgery. This VAF correlates with tumour size as dead tumour cells shed into the bloodstream and can be used with biPOD to detect post-surgery NSCLC relapse.

We modelled VAFs to obtain patient-specific growth rates. Estimates were heterogeneous, −0.03 ≤ *ρ* < 0.35 weeks^−1^, revealing dynamic patterns with different rates of exponential growth (Fig.3a), and Gaussian clustering of the rates (posterior means) revealed three groups of patients (Fig.3b). The first group (*n* = 52, clustered *ρ* = 0.001 weeks^−1^, s.d 0.004) showed a shallow growth rate, compatible with almost complete remission dynamics (absence of growth). The second group showed an intermediate rate (*n* = 33, clustered *ρ* = 0.044 weeks^−1^, s.d 0.036) and, finally, a third group had the highest growth rate (*n* = 14, clustered *ρ* = 0.207 weeks^−1^, s.d 0.079). With these parameters, the high-rate group had a doubling time that was 4.81 times faster than the intermediate group. Strikingly, these growth rate-based clusters were not correlated with the presence of key driver mutations, including *TP53, KRAS*, or *PIK3CA* (median Chi-square *p* = 0.859 for 11 tested mutations; Supplementary Table 1, Extended Data Fig. 4a). For instance, *TP53* mutations were distributed similarly across all groups (high: *n* = 7, intermediate: *n* = 18, low: *n* = 28), as were *KRAS* (high: *n* = 5, intermediate: *n* = 11, low: *n* = 14) and *PIK3CA* (high: *n* = 2, intermediate: *n* = 2, low: *n* = 3). This highlights a critical limitation of relying solely on genetic mutations in canonical NSCLC genes for patient stratification. In contrast, stratification based on growth rates derived from ctDNA dynamics provided a prognostic indicator for relapse or death. Survival analysis confirmed the predictive value of growth rate stratification (Fig. 3c), with statistical significance maintained after adjusting for sex and tumor stage in a Cox proportional hazard model (*p* ≤ 2 × 10^−16^, log-rank test).

**Figure 3.**
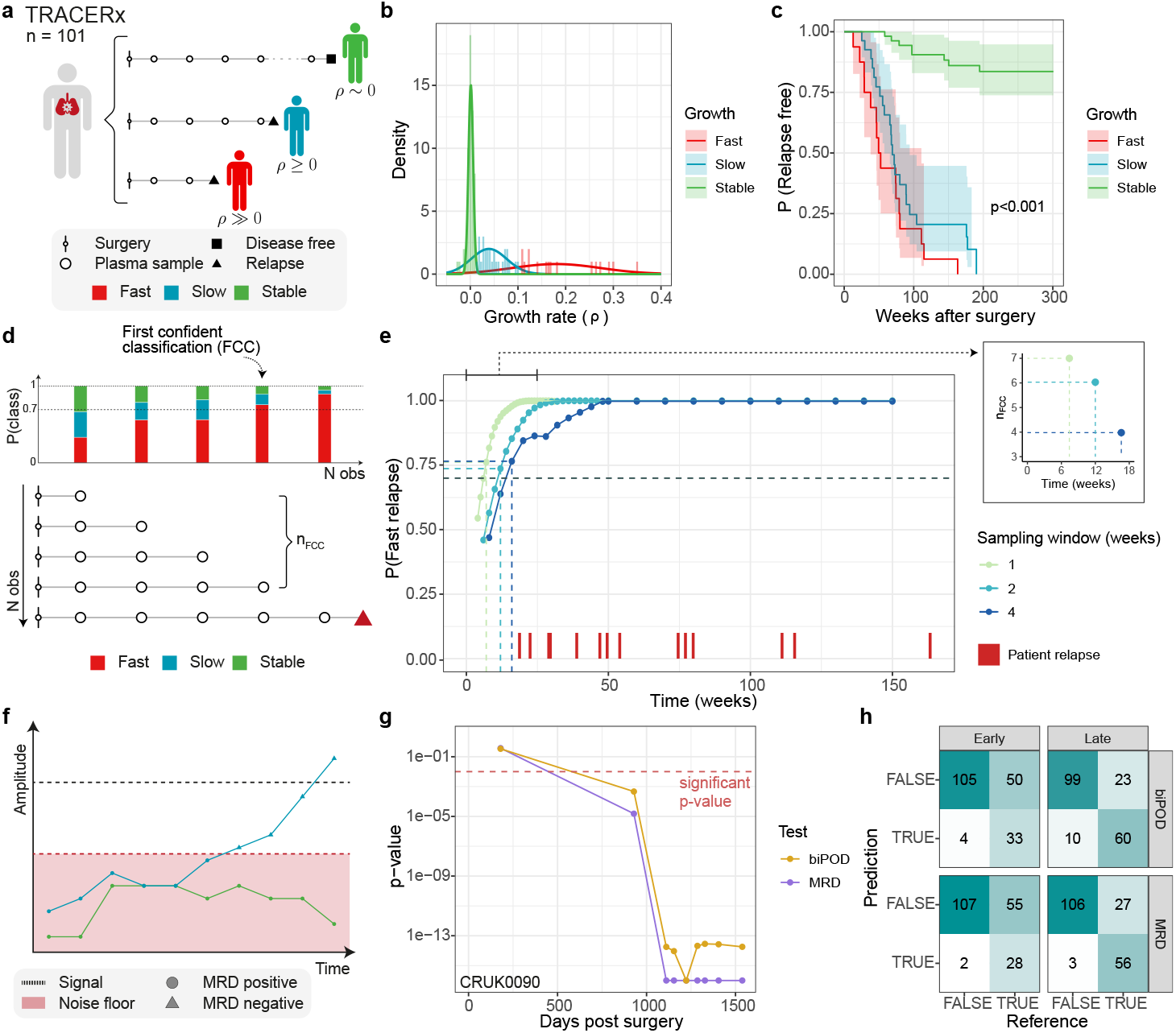
**a**. We used biPOD to stratify post-surgery disease progression patterns in 101 early-stage non-small-cell lung cancer from the TRACERx project^49,50^. Using growth parameters extracted from circulating tumour DNA (ctDNA), we identified patients who relapse fast to treatment, as opposed to those who relapse slowly or do not relapse at all (whole-cohort in Supplementary Fig.S8). **b**,**c**. Clusters of growth rates estimated in TRACERx using a Gaussian mixture applied to biPOD’s *ρ* estimates. According to the Bayesian information criterion, the best model has *k* = 3 components that corresponds to groups with different probability of relapse – Kaplan-Meier curves in panel (c). The inferred growth rates are statistically significant in a survival analysis (*p* ≤ 2 × 10^−16^, log-rank test). **d**,**e**. We simulated a real-time classification of these patients with biPOD and determined, as far as data becomes available, which patients relapse fast. We defined the first confident classification (FCC) time as the first ctDNA observation in which a patient is classifcied as fast with probability above 70%. Assuming a starting VAF of 3 × 10^−6^ and assuming that each patient is profiled every *n* weeks (*n* ∈ [1, 2, 4]), biPOD is able to confidently (*p* ≥ 70%) detect fast relapsing patients at most 16 weeks after surgery. **f**. We used biPOD to derive an alternative MRD testing, with the aim of modeling the techincal noise underlying the VAF measurements, since tumour-specific alternative reads might be observed in even in the absence of tumour cells and might be due to false positives. **g**. Comparison of MRD and biPOD test for patient CRUK0090, showing threshold of p-value that corresponds to the case in which the null hypothesis (i.e. alternative reads do not come from noise) cannot be rejected. **h**. Confusion matrices showing performance comparison between MRD and biPOD test, divided between early detection (before 120 days after surgery) and late detection (after 120 days after surgery).

Given these groups, a critical translational opportunity is the creation of statistical procedures that can flag patients in the high-risk group as far as their data becomes available in the clinic. This procedure is called online learning, and we simulated it by sampling data from the TRACERx cohort, using as control a standard approach that uses the whole data available for a patient (Fig 3d). We simulated VAF dynamics for both high and low-rate patients, assuming an initial VAF of 3 × 10^−6^ that was lower than the minimum VAF observed (2.66 × 10^−5^). To mimic realistic clinical sampling, we assumed patient data to become available every *n* weeks, ranging from weekly (fast, *n* = 1) to monthly (slow, *n* = 4) sampling. As far as data arrived, we measured how biPOD was confident in detecting fast relapsing patients. Strikingly, we discovered that even with monthly samples (*n* = 4) the high-risk group could be determined with high confidence (posterior probability of group assignment *p* ≥ 70%), as soon as 16 weeks (4 months) after surgery (Fig 3e). Therefore, biPOD detected fast relapsing patients on average 49 weeks before the clinical detection, corresponding to a mean anticipation of nearly a year, which suggests that a monthly sampling could greatly help anticipate intervention in post/surgery NSCL monitoring. This is especially important since these results imply that sampling more frequently could reduce the time needed for clinical detection by around 75%. This performance is quite remarkable, as this model could anticipate even the earliest clinical detection (131st day) by nearly two weeks, highlighting how this sampling scheme might be helpful even in the worst observed scenario.

One crucial point raised in the TRACERx ctDNA study^50^ was establishing an accurate test for minimal residual disease (MRD) from these VAFs. This task was challenging because the low-frequency regime of ctDNA VAFs could be confounded with sequencing noise (i.e., the probability of a false positive nucleotide readout), potentially leading to misleading relapse assessments (positive MRD) caused by technical noise (Fig.3f). Using biPOD (Extended Data Fig.4b), we derived an alternative MRD test that leverages the dynamics of observed patterns to estimate tumor-specific signals in the low-frequency regime. This approach involved regressing the mean VAFs at the time of surgery, 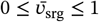, corresponding to the intercept with the y-axis (Extended Data Fig.5c), across three patient groups. Patients with negative MRD showed 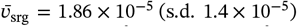, those with early positive MRD tests (within 120 days post-surgery) showed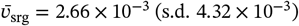, and those with late positive MRD tests (after 120 days post-surgery) showed 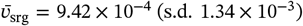. The estimates of 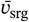 for negative MRD patients were significantly different from both early (*p* ≤ 2 × 10^−5^, t-test) and late positive MRD patients (*p* = 0.04, t-test) (Extended Data Fig.5d). We used the 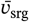 estimates from patients with negative MRD to model technical noise and developed a Beta-Binomial test to determine whether observed variant read counts could be explained by noise (null hypothesis *H*_0_; Extended Data Fig.4b). Failing to reject *H*_0_ would indicate negative MRD (example depicted in Fig 3g). When compared with the TRACERx MRD test at significance *α* = 0.01, our method yielded comparable results for identifying patients likely to relapse. Specifically, it demonstrated a true positive rate (TPR) of 0.397 before 120 days and 0.723 after 120 days, compared to the TRACERx MRD TPR of 0.337 and 0.675 in the early and late scenarios, respectively. The false positive rates (FPR) were 0.036 (TRACERx MRD FPR 0.018) in the early case and 0.091 (TRACERx MRD FPR 0.027) in the late case These findings highlight that studying ctDNA dynamics can offer powerful statistical tools for MRD detection, with performance metrics comparable to those achieved using complex genetic or bioinformatics pipelines, such as the established method discussed in the original paper, which estimates intralibrary, trinucleotide-specific sequencing error rates^50^. By focusing on the dynamics of ctDNA patterns, biPOD provides an interpretable test rooted in population dynamics principles, bridging statistical performance with biological insight.

### 2.5. Prediction of time-to-treatment in chronic lymphocyte leukemia

We sought to forecast time-to-treatment initiation in chronic lymphocytic leukemia (CLL), a malignancy characterized by a ‘watch-and-wait’ management approach. In this active surveillance scenario patients are monitored over time, and treatment is delayed until disease progression^53^. One strong indicator of progression is the time it takes for white blood counts (WBCs) to double^54^. Our aim was to predict time-to-treatment earlier than current clinical practice (Fig.4a).

**Figure 4.**
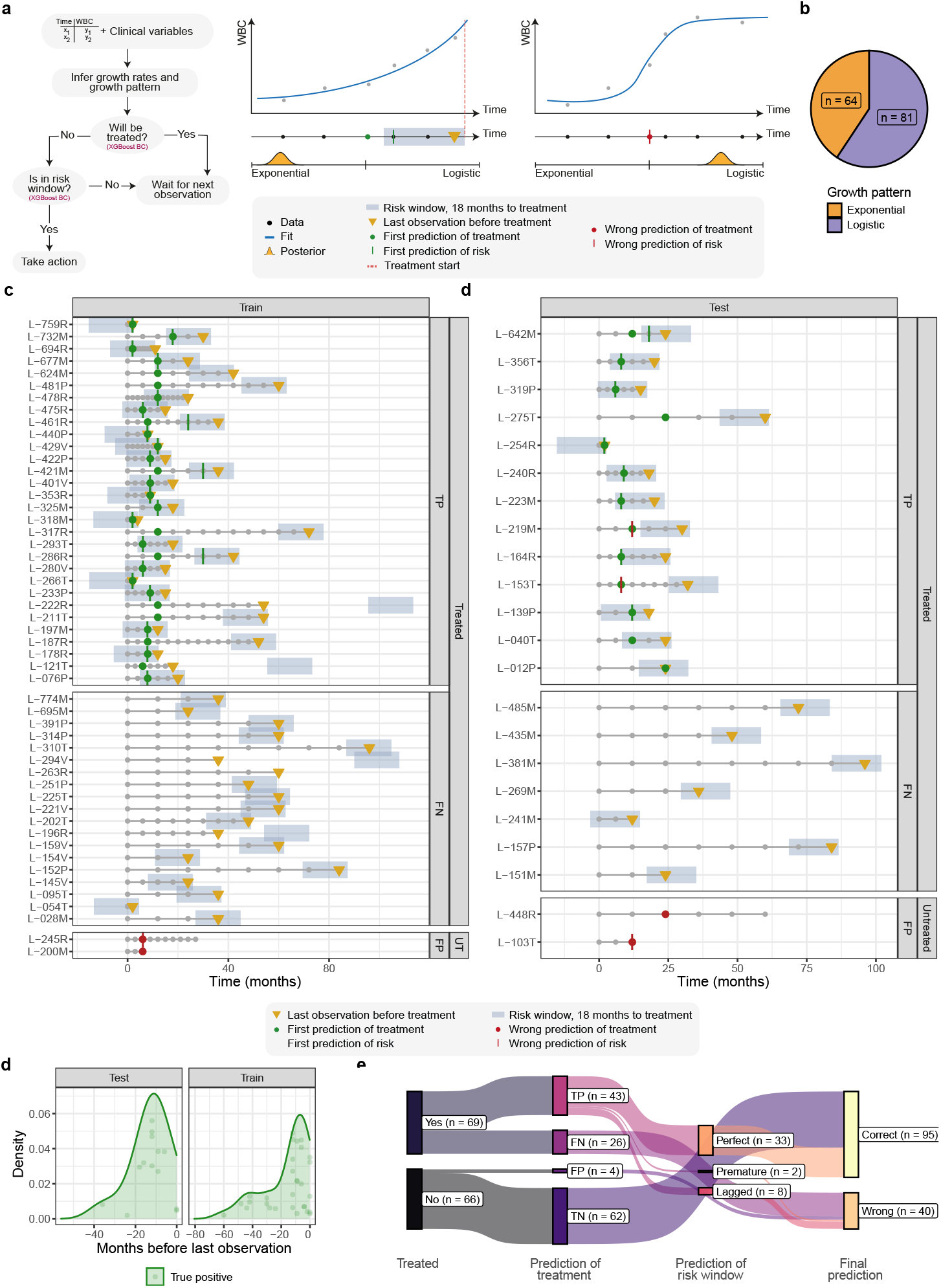
**a**. We used biPOD to predict the need for treatment and its proximity in a cohort of 145 patients with chronic lymphocytic leukemia (CLL) using longitudinal white blood cell (WBC) measurements. At each new observation, biPOD-inferred parameters from the WBC time series, combined with clinical variables, were used to predict whether a patient would require treatment and, if so, whether it would be needed within the next 18 months. Additionally, biPOD was used to classify patients based on their growth patterns, assuming either exponential or logistic growth. **b**. Number of cases with either logistic or exponential growth. **c**,**d**. Clinical history and performance of the biPOD-based model are shown for each patient. The golden triangle marks the last observation before treatment, while the light-blue area indicates the 18-month window preceding treatment. A green dot represents a correct prediction of treatment (true positive), while a red dot indicates an incorrect prediction (false positive). Similarly, the prediction of a patient being close to treatment is depicted by a vertical line—green if the patient is within the risk window and red if not. The full cohort, including true negative patients, is presented in Extended Data Fig.8. **e**. Distribution of prediction lead times for treatment predictions relative to the last available observation. Patients are grouped into training and test sets, with predictions color-coded as true positives or false positives. In both sets, the majority of predictions are made within 20 months to treatment. **f**. Sankey plot illustrating patient classification by the biPOD-based model. Patients are initially categorized as treated or untreated, leading to classifications as true positive (TP), false negative (FN), false positive (FP), or true negative (TN). For patients predicted as positive, the model further assesses whether they are within 18 months of treatment. These predictions are classified as perfect (within the time window), premature (before the time window), or lagged (after the time window). A patient is considered correctly predicted if both steps are accurate; otherwise, the prediction is deemed wrong.

We collected WBC data measured at irregular intervals from *n* = 145 CLL patients, of whom 79 (54%) underwent treatment (median of 7 observations per patient). For every patient (Extended Data Fig.7a), binary clinical variables commonly used for CLL stratification were recorded, including trisomy of chromosome 12 (Tri12), deletions in chromosomes 11, 13, and 17 (Del11, Del13, Del17), TP53 mutation status, immunoglobulin heavy chain variable region (IGHV) mutation status, and CD49d expression^55–58^. We standardised times, used only observations before treatment, and selected among exponential or logistic growth for patients with at least three observations (Supplementary Material). Overall, 81 patients were fit with a logistic growth (56%), and 64 with an exponential growth (44%; Fig.4b). Previous studies^19^ identified logistic and exponential growth patterns in two distinct cohorts (*n* = 21 and *n* = 85). However, to the best of our knowledge, this is the first study to evaluate these growth patterns in a much larger cohort. The earlier study differed from ours in its classification method, which was based solely on a logistic growth model. In their approach, growth was classified as logistic if the carrying capacity was below a specific threshold, exponential if the carrying capacity exceeded a much higher threshold, and undefined if it fell between these thresholds. Despite the different classification and not considering the undefined cases, the difference between the proportion of exponential cases found by Gruber et al.^19^ (*f*_*exp*_ = 0.314) and the one observed in our cohort (*f*_*exp*_ = 0.44) was not statistically significant (*p* = 0.07, Z test). Another similarity was that CLLs with logistic growth tended to have longer time-to-treatment, with a median of 36 months, compared with 23.3 months for exponential patients; however, this difference was not statistically significant (*p* = 0.21, Wilcoxon test). Interestingly, the previous study reported that logistic patients tended to have mutated IGHV status more frequently than exponential patients, and we observed the same difference between the exponential (34 out of 64, 53%) and logistic (57 out of 81, 70%) groups (*p* = 0.046, *χ*^2^ test).

Similarly to NSCLC patients from TRACERx, we used an online learning strategy to predict treatment time using biPOD’s parameters. In this cohort, we (*i*) first predicted treatment in a qualitative fashion, identifying if a patient will be eventually treated or not, and then, for patients predicted to eventually undergo treatment, (*ii*) we predicted if the patient was currently within 18 months close to its putative time-of-treatment. To implement both analyses, we used an advanced classifier (Methods) and a 70/30 split of the cohort after retaining only *n* = 135 patients with at least three observations. In both models, the predictors included current and initial WBC values, estimated growth rates, type of growth pattern and significant clinical variables. The first model performed well in both the training and test sets, achieving a sensitivity (true positive rate, TPR) of 0.612 and 0.650, respectively, with an exceptionally low false positve rate of 0.044 in the training set and 0.095 in the test set. Precision was also high, at 0.938 in the training set and 0.867 in the test set (Fig. 4c,d, full cohort in Extended Data Fig.8). Therefore, the first model is highly effective at early detection of patients requiring treatment. It correctly predicts the need for treatment more than 12 months before the clinical endpoint annotated in 53% of cases, achieving an average anticipation time of 15 months (standard deviation: 14 months; Fig. 4e). The second model, which builds on predictions from the first, exhibited a TPR of 0.821 in the training set and 0.727 in the test set. The precision was 0.958 in the training set and 0.727 in the test set (Fig. 4f). All the results are reported in in Supplementary Tables 5-6. Despite these signals were statistically clear, they could not have been predicted by genomics data. In fact, while IGHV status (*p* = 1.06 × 10^−6^, *χ*^2^ test) was significantly associated with treatment, none of the other tested clinical variables demonstrated a statistically significant association. These included the prognostic marker of CD49d expression, and four copy number alterations (Tri12, Del13, Del11, and Del17) that are frequently observed in CLLs. As for NSCLC, this finding underscores the limitations of static genomic data in accurately predicting treatment requirements and optimizing patient monitoring, at least compared to dynamic parameters derived from WBC with biPOD.

## 3. Discussion

Accurately modeling tumor dynamics is essential to capture patient-specific treatment responses patterns^59,60^, and transform tumor evolution parameters into translational biomarkers with predictive and prognostic value^61,62^. For example, tumor growth rates are key to monitoring patients and identifying those likely to reach clinical endpoints (e.g., disease relapse) more quickly, and provide insights into pre-existing and de novo drug resistance. Although deriving tumor evolution parameters from cross-sectional measurements is challenging, the growing availability of longitudinal data enables mathematical models to address these critical biological questions.

This work presented the first Bayesian framework that uses population genetics to infer key patient-specific tumour evolution parameters from various data sources. We achieved accurate predictions even with noisy and limited observations from non-invasive liquid biopsies, or tumour-size estimates from in vivo experiments, and used our tool in three detailed case studies. Within a clinical trial for metastatic colon cancers, we confidently captured pre-existing resistance by RAS mutations. In the popular TRACERx cohort of lung cancer patients, instead, we established prognostic tumour evolution patterns determined by distinct growth rates and derived an innovative evolution-based test for minimal residual disease in ctDNA. Moreover, in a novel cohort of chronic leukemia patients, we established the first automatic classifier to predict time-to-treatment from routinely blood testing. In this case, first, we classified eventually-to-be-treated patients, and then we precisely assessed if a patient is within 18 months from requireing treatment time. In this cohort, we also reported the largest evidence of non-exponential growing tumours, augmenting further evidence from earlier works^19^. To the best of our understanding, our findings for both the lung and leukemia cohorts are innovative and push forward our understanding of these diseases.

Our results support earlier findings that tumor evolution is predictive^3,63,64^, and demonstrate that predictors can be integrated into online learning algorithms that process data sequentially, unlike batch learning techniques that use entire training datasets. This is particularly well-suited for clinical monitoring of cancer patients where data is incrementally acquired through routine exams. Our model-based framework can extrapolate future disease dynamics following Bayesian principles. For this reason (*i*) every prediction (with its full probability distribution) has an intrinsic notion of uncertainty, and (*ii*) can encode piori biological or clinical evidence.

While this new class of frameworks is certainly innovative, it remains open to significant improvements. A current statistical simplification is the reliance on a single longitudinal measurement, such as a median variant allele frequency value (as in lung) or a single marker (as colon and leukemia). Tumors, however, are complex ecosystems of subclones with distinct evolutionary parameters^6,7,65^ and a more advanced framework could model heterogeneous evolutionary parameters within a patient^3,24^. While remains challenging with current bulk ctDNA data, the convergence of single-cell technologies and liquid biopsy methods offers a promising avenue for this ambitious goal^66^. Another limitation is the lack of reconciliation of intra-patient heterogeneity, despite substantial evidence indicating the contingency of the evolutionary process^67,68^. In line with algorithms that identify repeated tumour evolution^10,69^, there is likely untapped statistical information that could be better harnessed by pooling multiple patients. Popular frameworks, such as temporal factor analysis^70^, provide inspiration for approaches that integrate shared evolutionary signals while preserving patient-specific insights.

In conclusion, our approach represents a significant advancement in analyzing tumor dynamics from longitudinal observations, addressing critical gaps in our understanding of cancer progression. Its successful application in various clinical scenarios suggests that it can be a powerful tool for researchers and clinicians alike, paving the way for more personalised and effective cancer treatment strategies. The integration of temporal dynamics through biPOD tries to fill some of the gaps left by traditional static genomic analyses, which often fail to capture the complexities of tumor evolution and treatment response. By focusing on dynamic patterns, biPOD enables deeper insights into cancer biology, offering new opportunities to anticipate clinical events, optimize patient monitoring, and tailor interventions in ways previously unattainable. While continued development and validation of this framework will be essential as we strive to improve our understanding of cancer evolution and response to therapy, biPOD highlights the critical role of studying dynamics to address unresolved challenges in oncology and to enhance patient outcomes.

## Data availability

biPOD is available as an open-source R package hosted at https://caravagnalab.github.io/biPOD and the code to replicate the analysis of this paper, along with data, is available at Zenodo https://zenodo.org/records/14329228

## Authors contribution

GS, RB, LE, and GC developed the framework, which GS implemented. GS implemented synthetic data and tested the model. Data were provided by GL, MIP, AZ, and VG. GS gathered and analysed the data in collaboration with AZ, VG, LE, and GC. GS, LE and GC drafted the manuscript, which all authors approved. LE and GC conceptualised and supervised this work.

## Funding

The research leading to these results has received funding from AIRC under MFAG 2020 - ID. 24913 project – P.I. Caravagna Giulio. We acknowledge financial support under the National Recovery and Resilience Plan (NRRP), Mission 4, Component 2, Investment 1.1, Call for tender No. 1409 published on 14.9.2022 by the Italian Ministry of University and Research (MUR), funded by the European Union – NextGenerationEU– CUP J53D23015060001, as well as under Decreto Direttoriale No. 104 published on 02-02-2022 by MUR (NextGenerationEU– CUP J53D23003860006), under the PNRR-MAD-2022-12375673 (Next Generation EU, M6/C2 CALL 2022) and under the PNRR-MCNT2-2023-12378037 (Next Generation EU, M6/C2 CALL 2023), both published by the Italian Ministry of Health, Rome, Italy.

## Acknowledgments

We wish to thank Benjamin Werner, Andrea Sottoriva and Nicola Valeri for sharing data from the PROSPECT-C clinical trial and linked with publication^44^ Additionally, we kindly thank Nicholas McGranahan and Alexander M. Frankell for their assistance in helping us interpret the public TRACERx data^50^.

## Competing interests

The authors declare no competing interests.

## Ethical approvals

Data provided by GL, MIP, AZ and VG was collected within studies carried out in accordance with the Declaration of Helsinki, and approved by the Institutional Review Boards of the National Cancer Centre in Aviano (Approvals Nos. IRB-05-2010, IRB-05-2015).

---

**Extended Data Figure 1.**
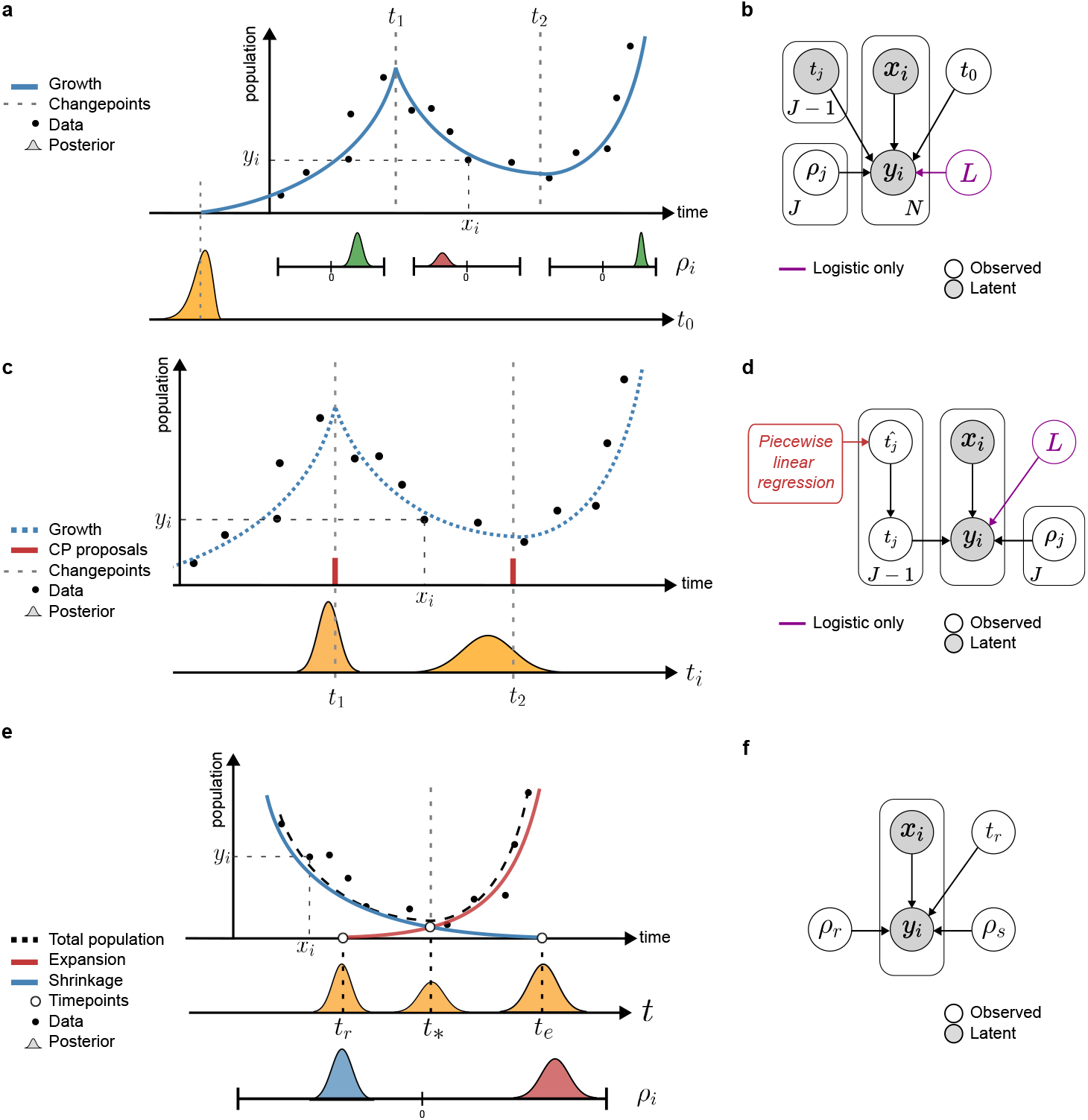
Cartoon of each Bayesian inference task possible in biPOD, with the probabilistic graphical model reporting observed variables in grey and latent ones in white. Priors are reported in Supplementary Material. **a, b**. (Task 1) Our framework uses longitudinal (*x*_*i*_, *y*_*i*_) data of tumour size (or any correlated covariate) and time points *t*_*i*_ where tumour dynamics change to infer a posterior distribution over the effective tumour growth rates *ρ*_*i*_ for each time window, as well as an upper bound on time *t*_0_ < *t*_1_ when the first tumour cell was born. The model considers both exponential and logistic tumour growth models, and in the cartoon, it learns a positive exponential growth rate for *t* < *t*_1_ and *t* > *t*_2_ and a negative growth rate for *t*_1_ ≤ *t* ≤ *t*_2_. **c, d**. (Task 2) Our framework can detect time points where tumour dynamics change, even if these are unknown. This is achieved by combining piecewise linear regression and a Bayesian model to compute a posterior distribution over each *t*_*i*_. **e, f**. (Task 3) In regions with U-shaped dynamics, one population overcoming another can describe tumour shrink and re-growth. Our framework can infer a posterior distribution for the growth rates *ρ*_*i*_ of both populations, as well as times in which the populations are born (*t*_*s*_) or extinct (*t*_*e*_), and the time *t*_∗_ when the surviving population out-competes the one that goes extinct.

**Extended Data Figure 2.**
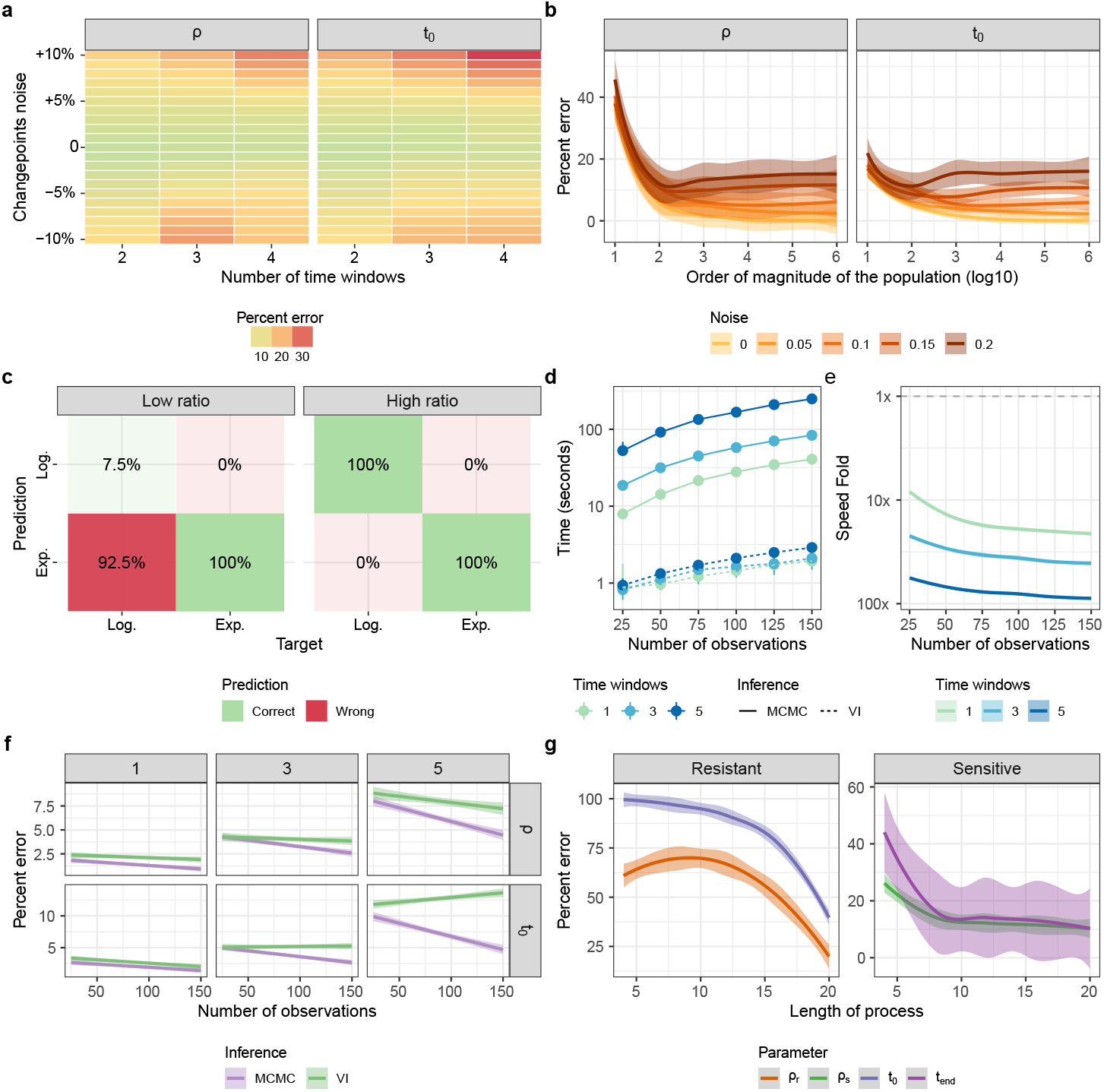
Simulated data tests validate biPOD performance in silico. **a, b, c**., **d**., **e**., **f**. For the task in Extended Data Fig. 1a, in panel (a), we report percent errors of growth rates and instant of birth depending on noisy input change-points (from −10% to +10%). In panel (b), we report the error relative to the magnitude of the simulated population (the higher the latter, the lower the stochasticity of the process and the inference error). In panel (c), we report the performance of biPOD model selection. The performance is divided based on the carrying capacity ratio, i.e. the ratio between the maximum observed population size and the true carrying capacity. The ratio is defined low if ≤ .35, and high otherwise. In panel (d) and (e), we report the computational time in logscale for biPOD implementations that use variational inference (VI) and Markov Chain Monte Carlo (MCMC) sampling, with an increasing number of observations and time windows. In panel (f) we report the trend of the error on growth rates and instant of birth for VI and MCMC sampling, depending on number of observations and time windows. **g**. For the task in Extended Data Fig. 1e, in panel (g) we report percent errors for the growth rate of resistant and sensitive populations and for their characteristic timepoints, i.e. time of birth *t*_*r*_ and time of extinction *t*_*s*_.

**Extended Data Figure 3.**
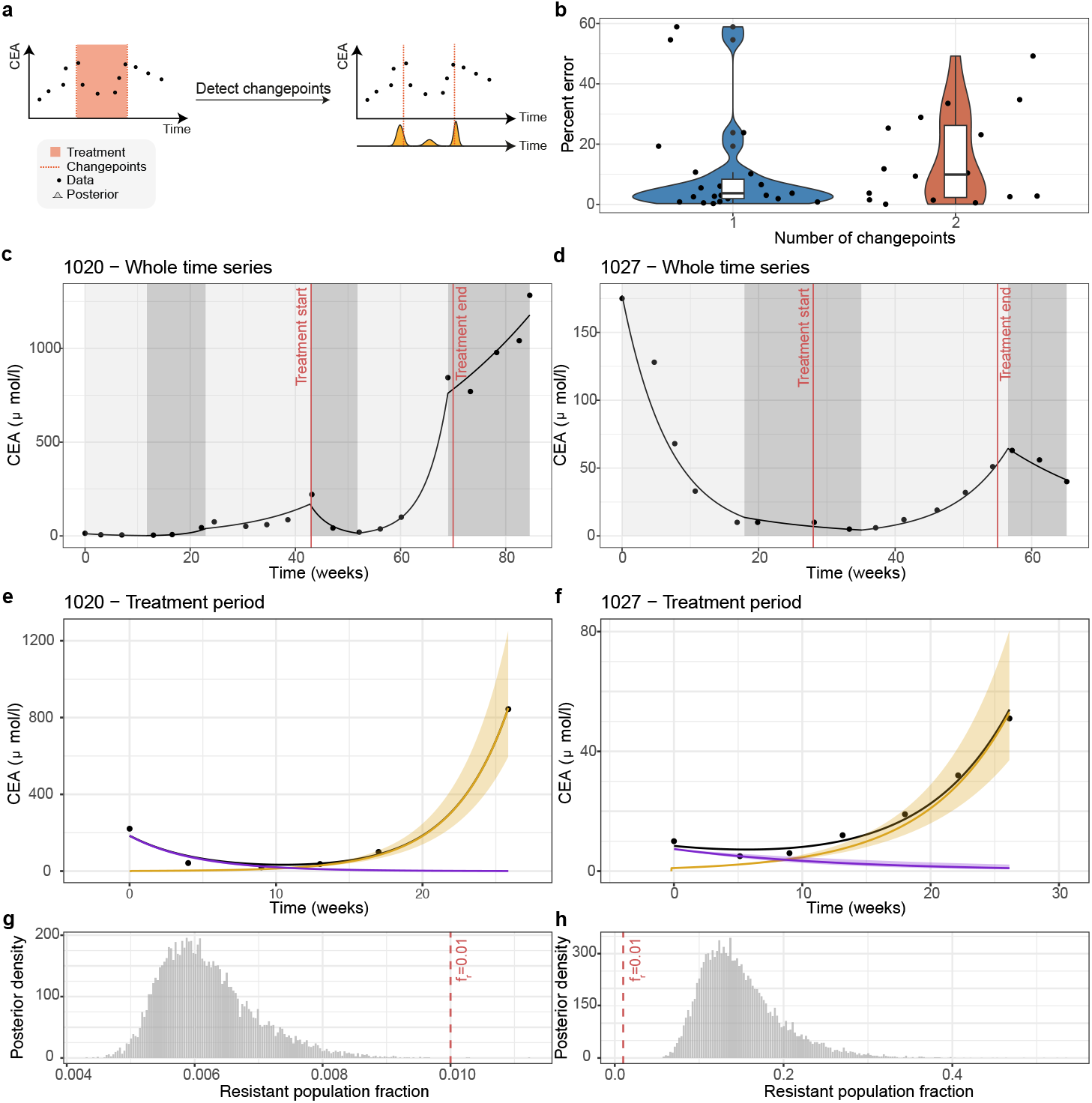
**a-b**. We used biPOD to identify changepoints in longitudinal carcinoembryonic antigen (CEA) data from 27 patients in the PROSPECT-C phase II colorectal cancer clinical trial^44^. The detected changepoints were compared to the known treatment start and end dates. Panel (b) reports the percentage errors for two scenarios: (i) patients with two changepoints, where treatment occurs in the middle of the time series, and (ii) patients with one changepoint, where the end of treatment aligns with the last available observation. **c-d**. Fit with inferred changepoints (dark and light gray areas) for two patients (i.e. 1020 and 1027). The inferred changepoints in the first case perfectly aligns with the treatment dates, while for the second case the errors are larger. The larger errors can be explained by the fact that the treatment has smaller effect on the poulation, making the change in dynamics less noticeavle. **e-f**. U-shaped dynamics for patients 1020 and 1027 are shown. The black line represents the total population dynamics, while the blue and red lines represent the dynamics of the resistant and sensitive populations, respectively. **g-h**. Posterior distributions of the resistant population fraction (*f*_*r*_) are shown for patients 1020 and 1027. Patient 1020 has a median *f*_*r*_ below 1%, suggesting that the resistant population is not pre-existing. In contrast, patient 1027 shows a median *f*_*r*_ of approximately 15%, indicating a significant pre-existing resistant population.

**Extended Data Figure 4.**
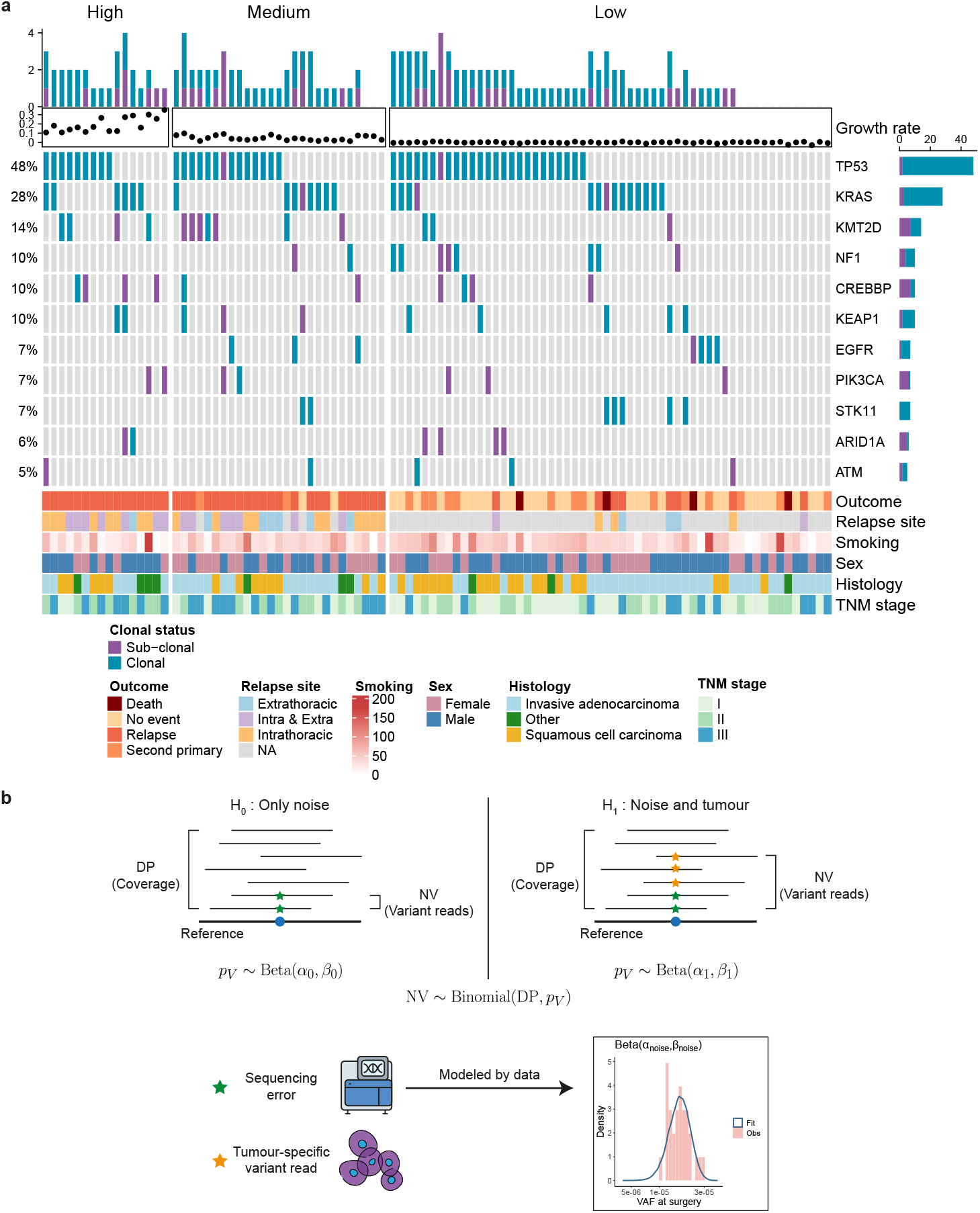
**a**. Oncoprint showing the distribution of genomic alterations in the tracerX cohort divided by growth rate class (i.e. High, Medium, and Low) along with the clonal status of each mutation. Additional covariates such as gender and TNM stage are also reported for each patient. **b**. Graphical representation of the biPOD based alternative MRD testing. The null hypothesis *H*_0_ assumes that the observed mutated reads are only due to technical noise, which is modeled using the regressed VAF values from patients with negative MRD tests. Under the null hypothesis, the number of variant reads (NV) given a total number of reads (DP) would follow a BetaBinomial(DP,*α*_*noise*_,*β*_*noise*_) distribution. If the observed reads deviate from this assumption, the alternative hypothesis (i.e. variant reads are due to noise and tumour cells) is accepted.

**Extended Data Figure 5.**
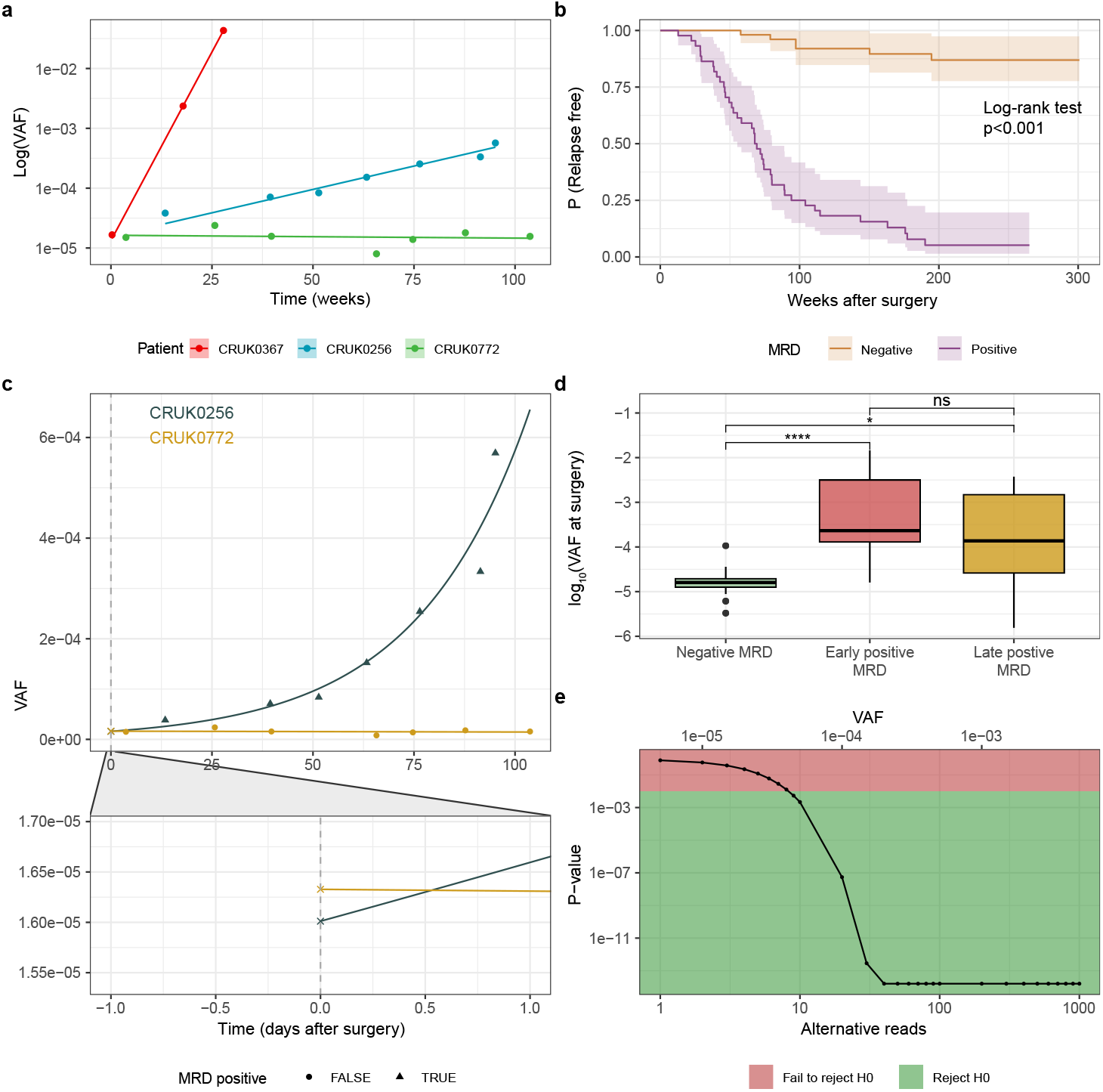
**a**. biPOD fits using VAF measurements for three patients, exhibithing fast (red), medium (blue) and slow (green) growth. **b**. Kaplan-Meier curves reporting the probability of relapse for patients in the three different clusters. The inferred growth rates are statistically significant in a survival analysis (*p* ≤ 2 × 10^−16^, log-rank test). **c**. biPOD fits using VAF measurements for patients with similar inferred VAF at surgery but very different dynamics: while CRUK0256 has a high growth rates (0.035 ≤ *ρ* ≤ 0.0366) and all observations resultest postive to an MRD test, CRUK0772 has low growth rate (−0.001 ≤ *ρ* ≤ 0.0001) and all obsevrations are MRD negative. **d**. Box-plot showing the inferred VAF at surgery stratifying patients by MRD test outcome (Negative, Early positive (within 120 days post-surgery), and Late positive (after 120 days post-surgery)). The distirbution for the negative patients is statistically different from both the Late positives (*p* = 0.04, Wilcoxon test) and the Early positives (*p* = 0.00002, Wilcoxon test). **e**. Example of biPOD test p-values given a fixed coverage of 2e5 reads and varying observed alternative reads or VAF values. In this examples, around 10 reads are required in order to reject the null hypothesis.

**Extended Data Figure 7.**
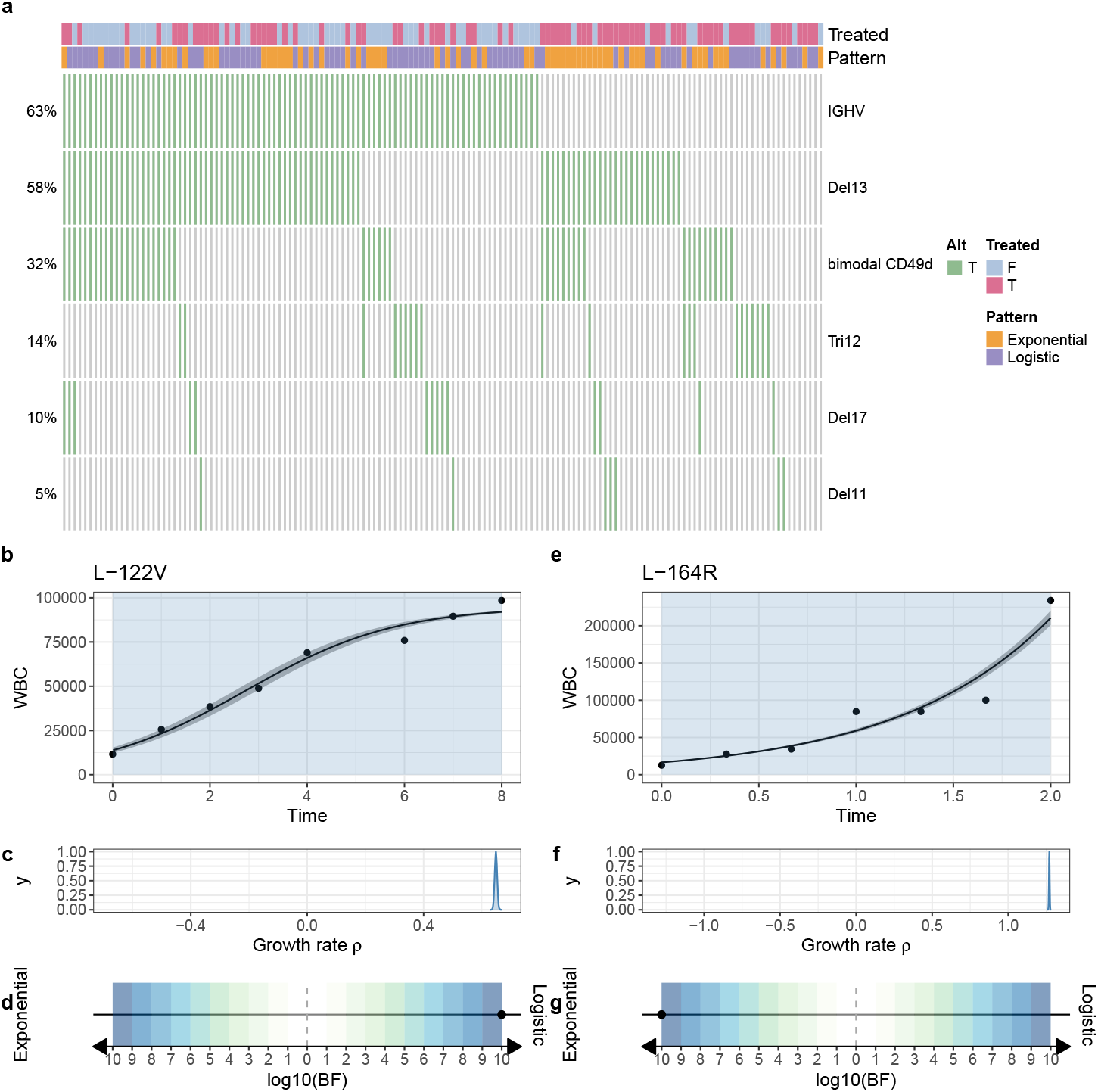
**a**. Oncoprint showing the distribution of genomic alterations in the CLL cohort. Additionally, the treatment status is reported as well as the growth pattern inferred using biPOD. **b-c**. Example of patients exhibithing a logistic (b) and exponential (c) growth pattern. Additionally, for each patient are reported the posterior distribution of the growth rate and the bayes factor used to decide between logistic and exponential.

**Extended Data Figure 8.**
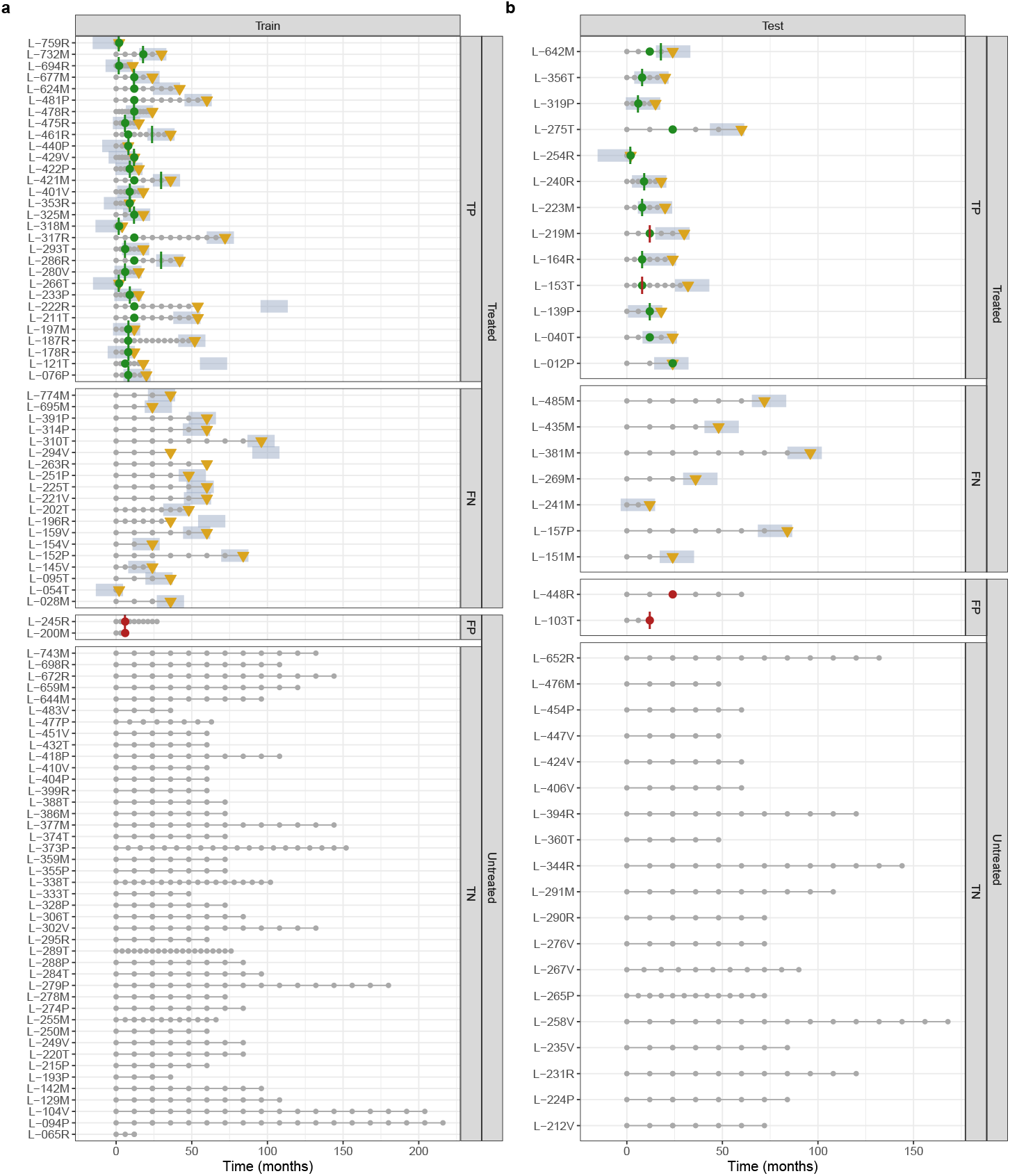
**a**. Clinical history and performance of the biPOD-based model for the full cohort. The golden triangle marks the last observation before treatment, while the light-blue area indicates the 18-month window preceding treatment. A green dot represents a correct prediction of treatment (true positive), while a red dot indicates an incorrect prediction (false positive). Similarly, the prediction of a patient being close to treatment is depicted by a vertical line—green if the patient is within the risk window and red if not.

## Notes

### Competing Interest Statement

The authors have declared no competing interest.

### Summary of Updates

Title change and minor changes to match new submission

