## Supplementary Material for "Predicting tumour evolution and drug resistance from heterogenous longitudinal cancer data"

[1]*Department of Mathematics, Informatics and Geosciences, University of Trieste, Italy*
[2]*Department of Business, Economics, Mathematics and Statistics, University of Trieste, Italy*
[3]*Department of Biomedicine and Prevention, University Tor Vergata, Rome, Italy.*
[4]*Clinical and Experimental Onco-Hematology, Centro di Riferimento Oncologico, Aviano*
[5]*Area Science Park, Italy*
[†]*These authors contributed equally to this work*

Creative Commons CC BY 4.0

#### A. Supplementary Figures

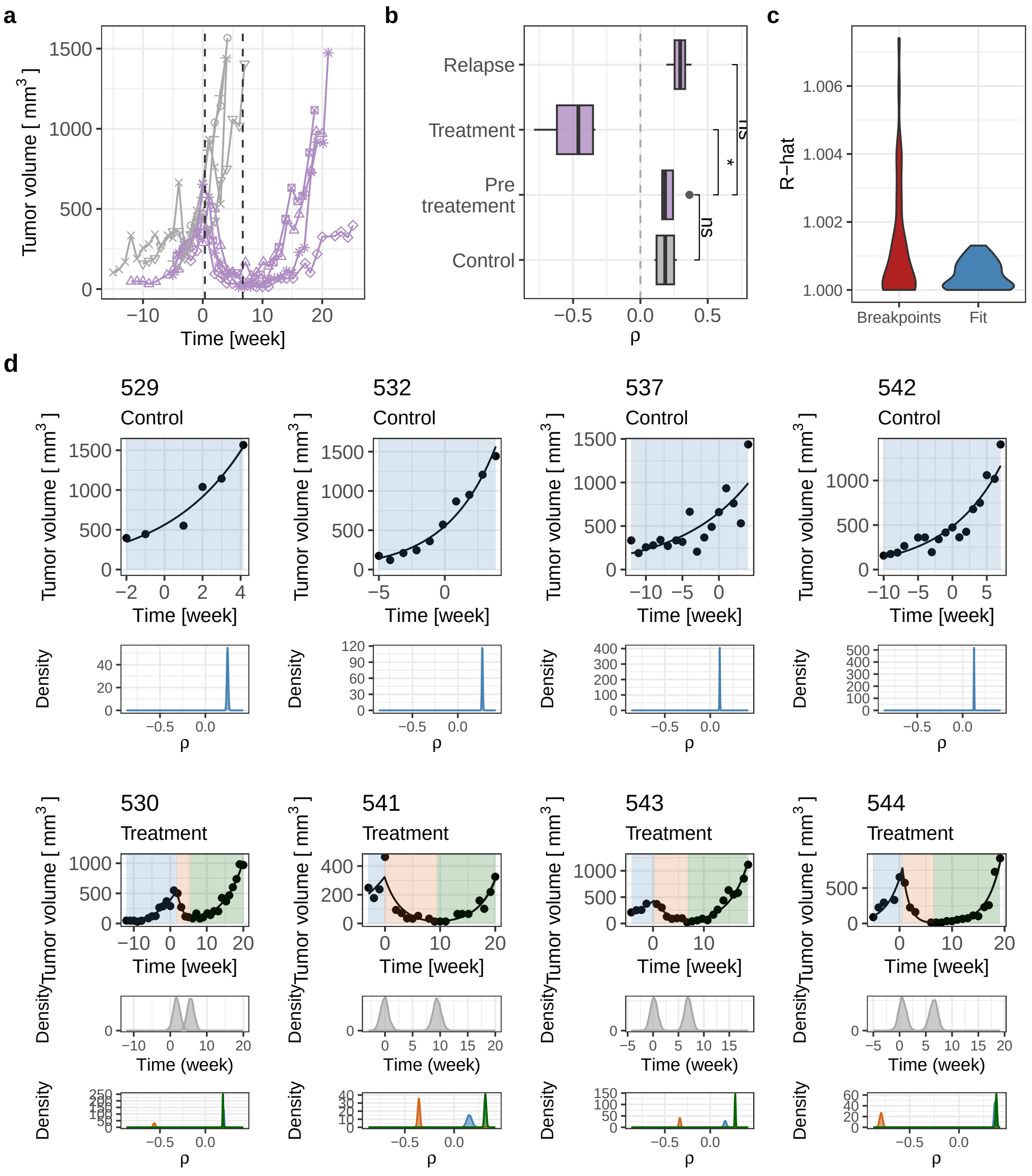

**Supplementary Figure S1.** Detailed report of patient with ID=828 from the Sauer dataset.[1] **a.** Tumour volume over time for control and treatment groups, with individual samples as points and dashed lines for median inferred change-points in the treated group. **b.** Box-plot of inferred growth rates ($\rho$) by treatment stage, coloured by group. **c.** Violin plot of R-hat obtained during the inference, divided by R-hat pertaining to the breakpoints detection and the fit of the dynamics. **d.** Analysis of each sample, showing tumour volume with model fit, posterior distributions of inferred change-points (treated samples), and growth rate density distributions.

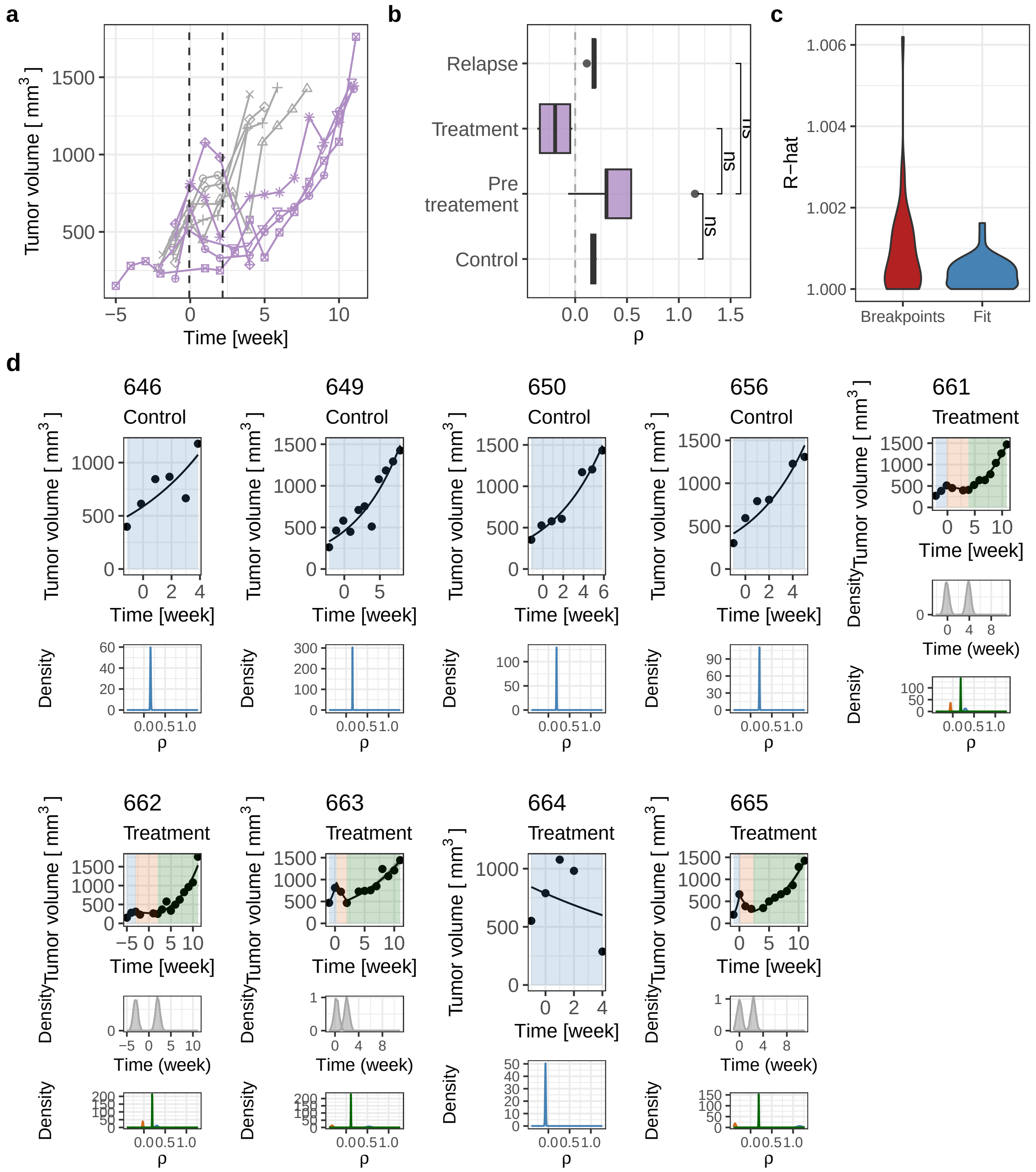

**Supplementary Figure S2.** Detailed report of patient with ID=600 from the Sauer dataset.[1] **a.** Tumour volume over time for control and treatment groups, with individual samples as points and dashed lines for median inferred change-points in the treated group. **b.** Box-plot of inferred growth rates ($\rho$) by treatment stage, coloured by group. **c.** Violin plot of R-hat obtained during the inference, divided by R-hat pertaining to the breakpoints detection and the fit of the dynamics. **d.** Analysis of each sample, showing tumour volume with model fit, posterior distributions of inferred change-points (treated samples), and growth rate density distributions.

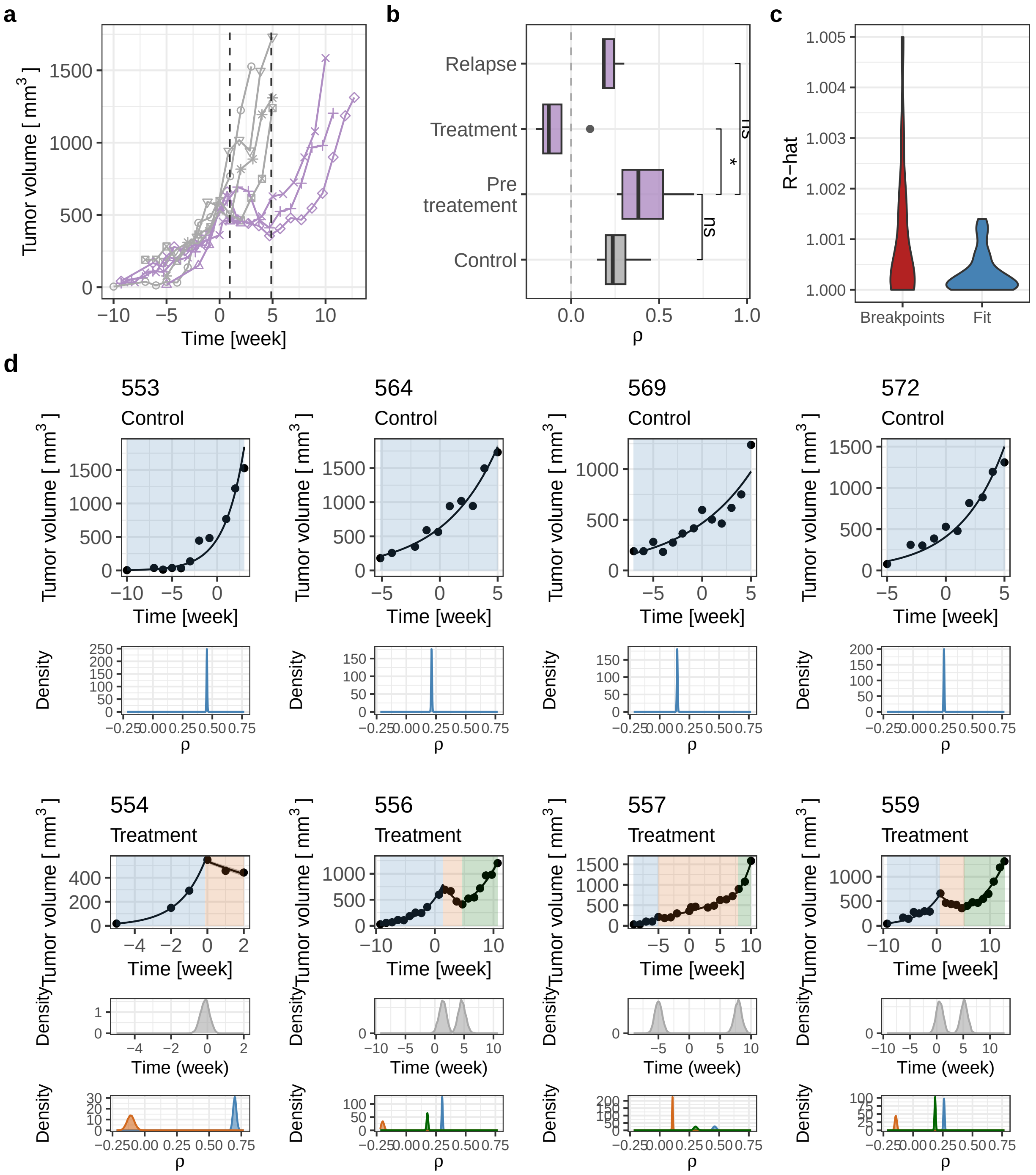

**Supplementary Figure S3.** Detailed report of patient with ID=771 from the Sauer dataset.[1] **a.** Tumour volume over time for control and treatment groups, with individual samples as points and dashed lines for median inferred change-points in the treated group. **b.** Box-plot of inferred growth rates ($\rho$) by treatment stage, coloured by group. **c.** Violin plot of R-hat obtained during the inference, divided by R-hat pertaining to the breakpoints detection and the fit of the dynamics. **d.** Analysis of each sample, showing tumour volume with model fit, posterior distributions of inferred change-points (treated samples), and growth rate density distributions.

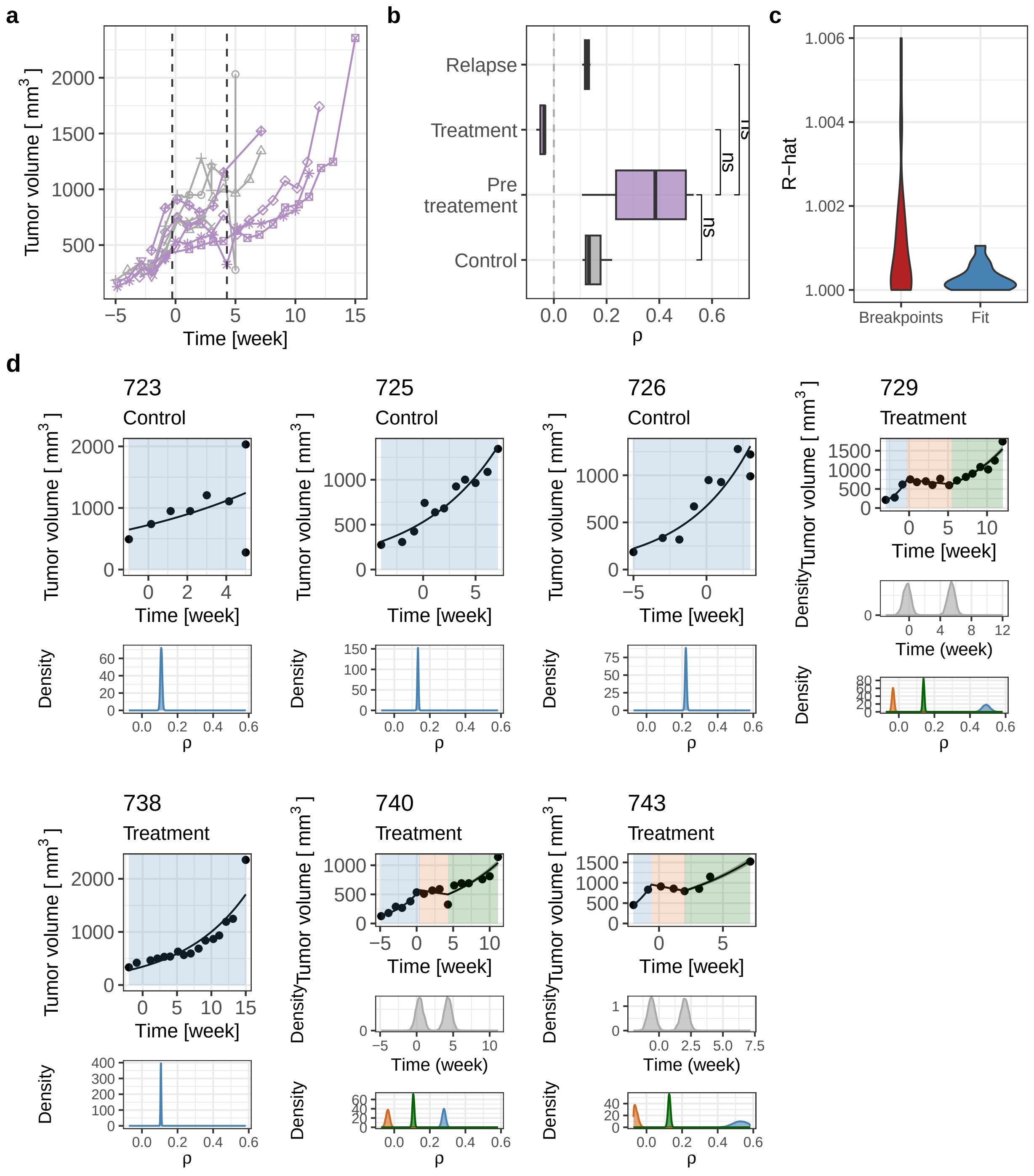

**Supplementary Figure S4.** Detailed report of patient with ID=831 from the Sauer dataset.[1] **a.** Tumour volume over time for control and treatment groups, with individual samples as points and dashed lines for median inferred change-points in the treated group. **b.** Box-plot of inferred growth rates ($\rho$) by treatment stage, coloured by group. **c.** Violin plot of R-hat obtained during the inference, divided by R-hat pertaining to the breakpoints detection and the fit of the dynamics. **d.** Analysis of each sample, showing tumour volume with model fit, posterior distributions of inferred change-points (treated samples), and growth rate density distributions.

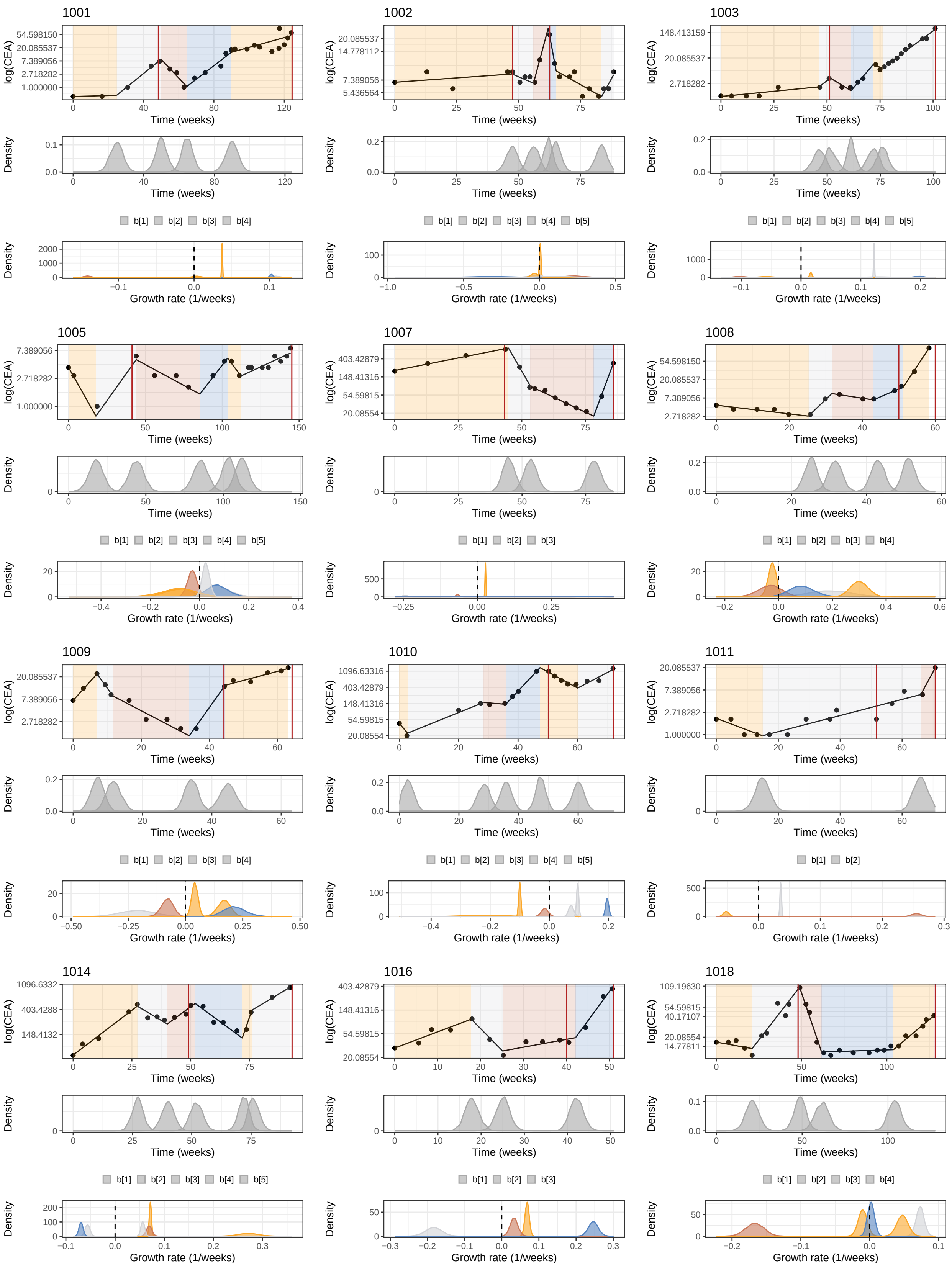
1001
1002
1003
1005
1007
1008
1009
1010
1011
1014
1016
1018
log(CEA)
Time (weeks)
Density
Growth rate (1/weeks)
b[1] b[2] b[3] b[4] b[5]

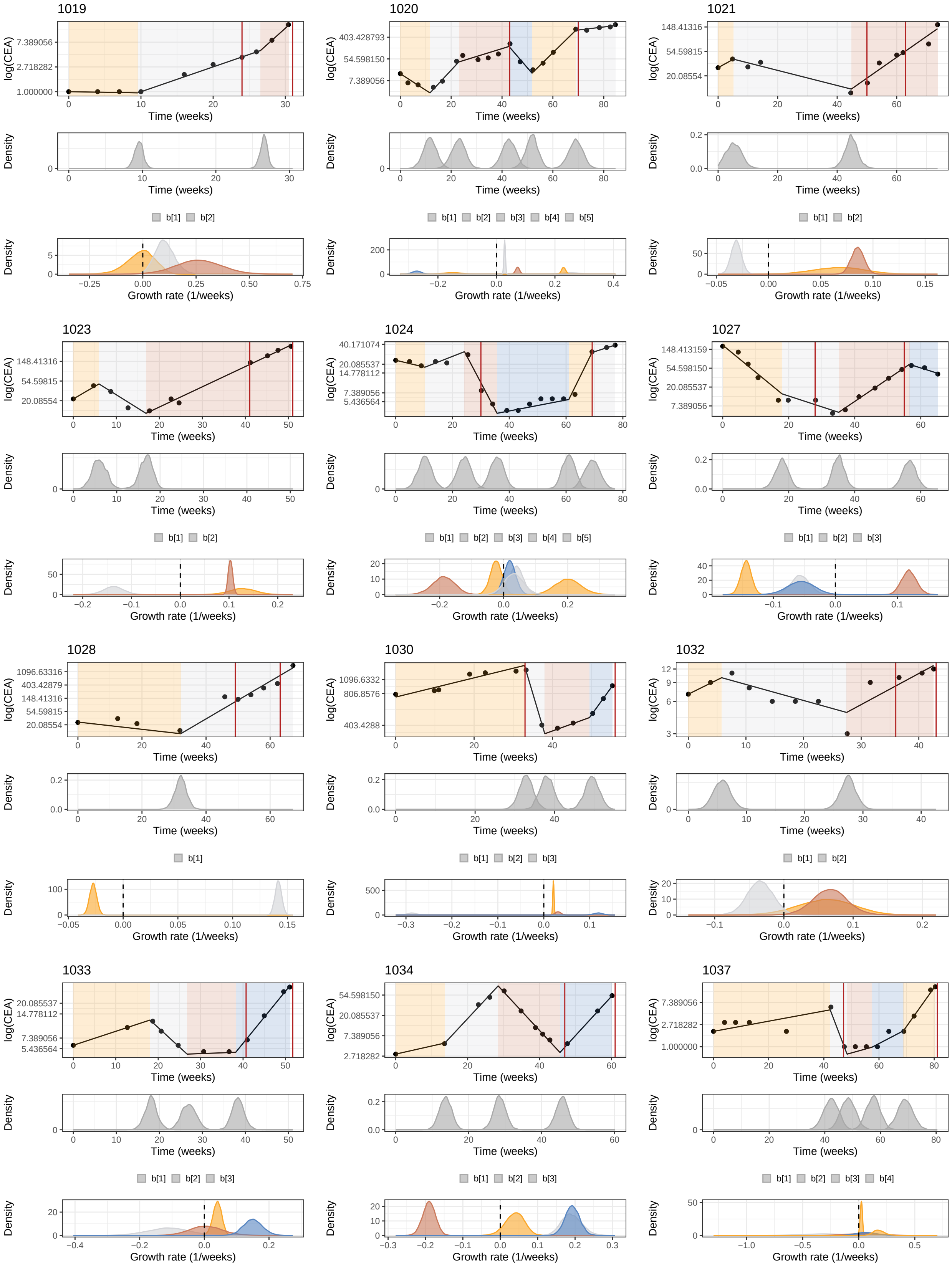
1019
1020
1021
1023
1024
1027
1028
1030
1032
1033
1034
1037
log(CEA)
Time (weeks)
Density
Growth rate (1/weeks)
b[1]
b[2]
b[3]
b[4]
b[5]

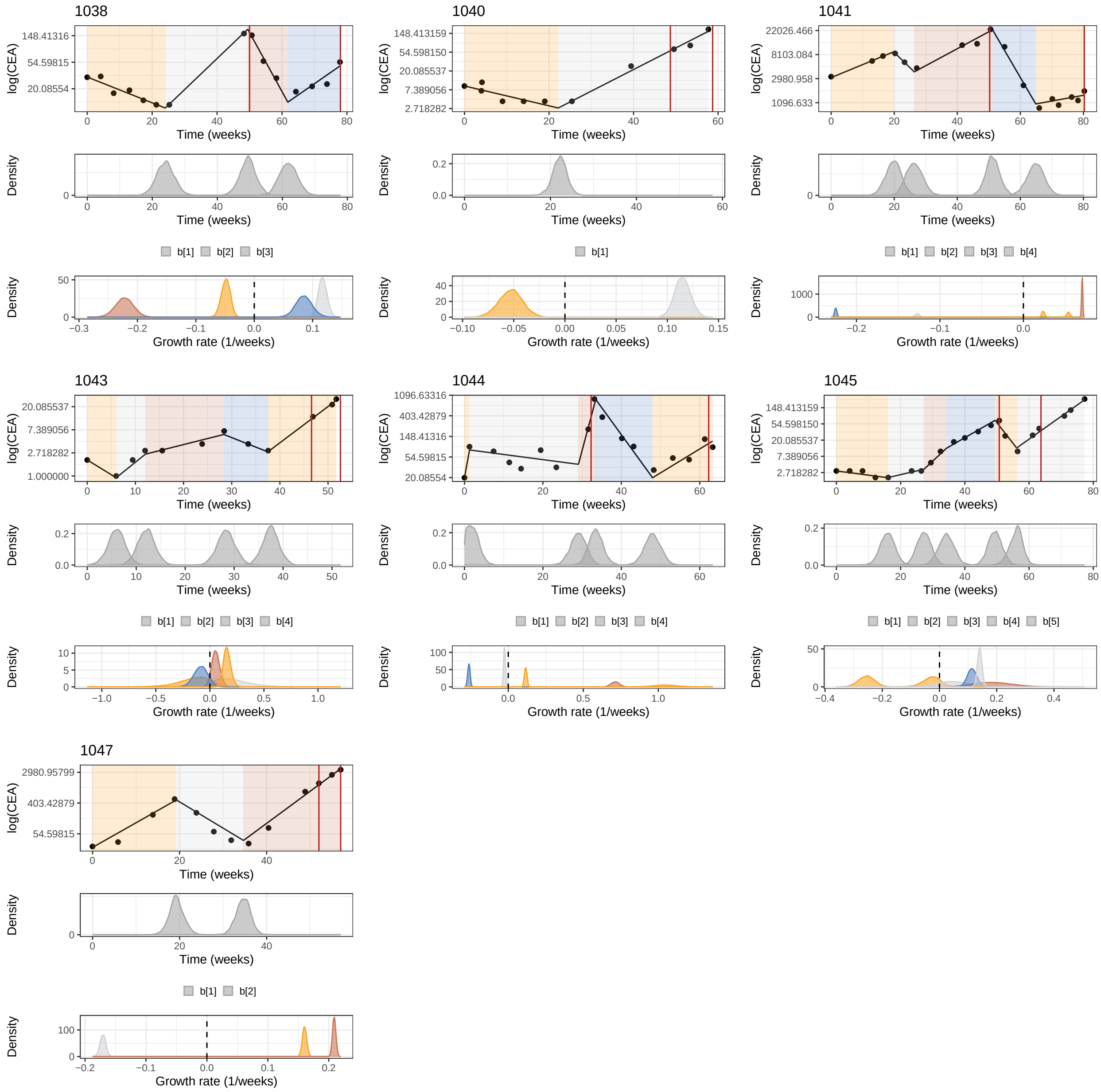

**Supplementary Figure S5.** Fit of the complete time-series of CEA (carcinoembryonic antigen) measurements for each patient in the cohort from[2], segmented into different time windows (different background colors) by biPOD. The start and end of the treatment are marked by red vertical lines. For each patient is also reported the posterior density of the inferred changepoints (in gray), and the posterior densiry of the inferred growth rates.

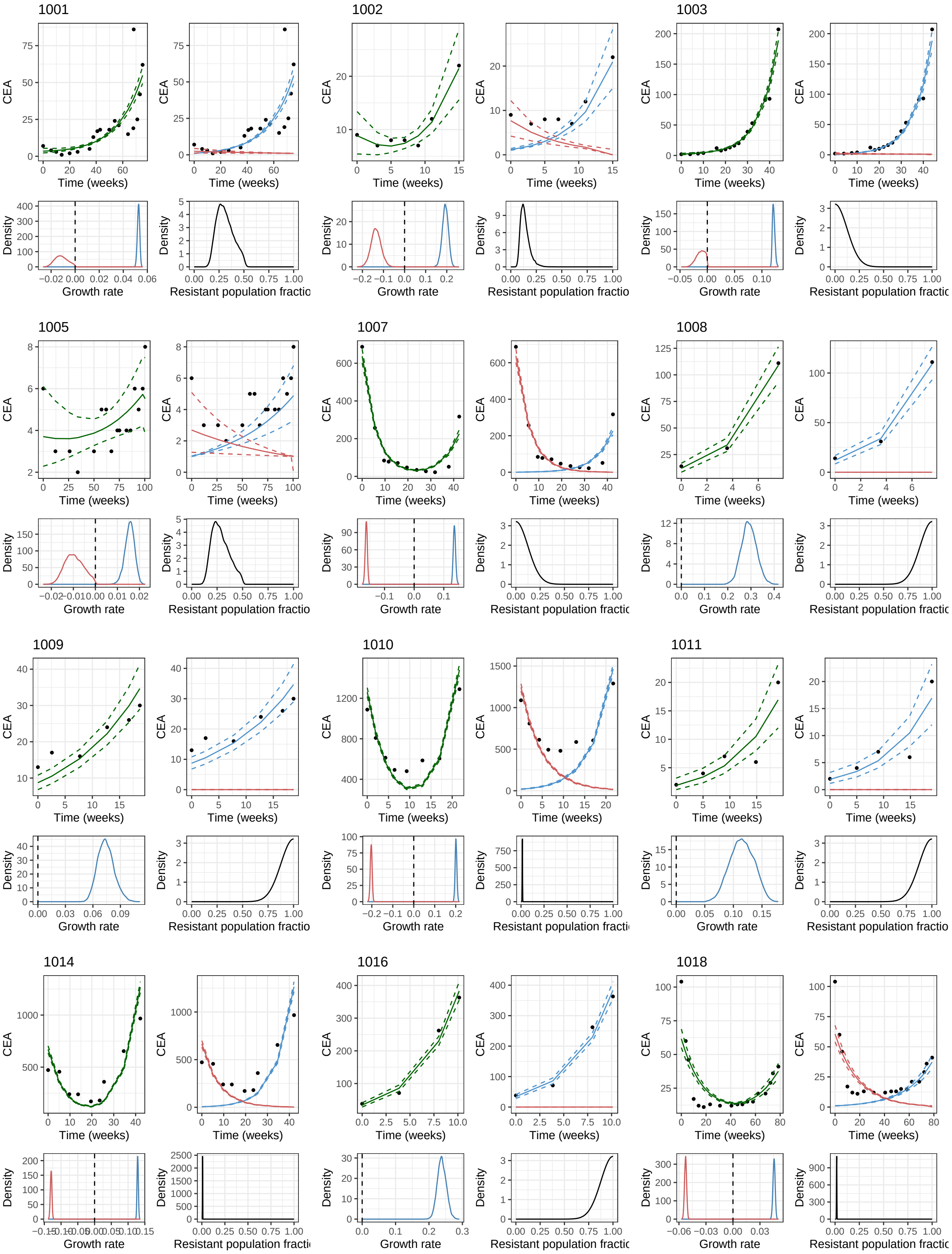
1001
1002
1003
1005
1007
1008
1009
1010
1011
1014
1016
1018
CEA
Time (weeks)
Density
Growth rate
Resistant population fractio

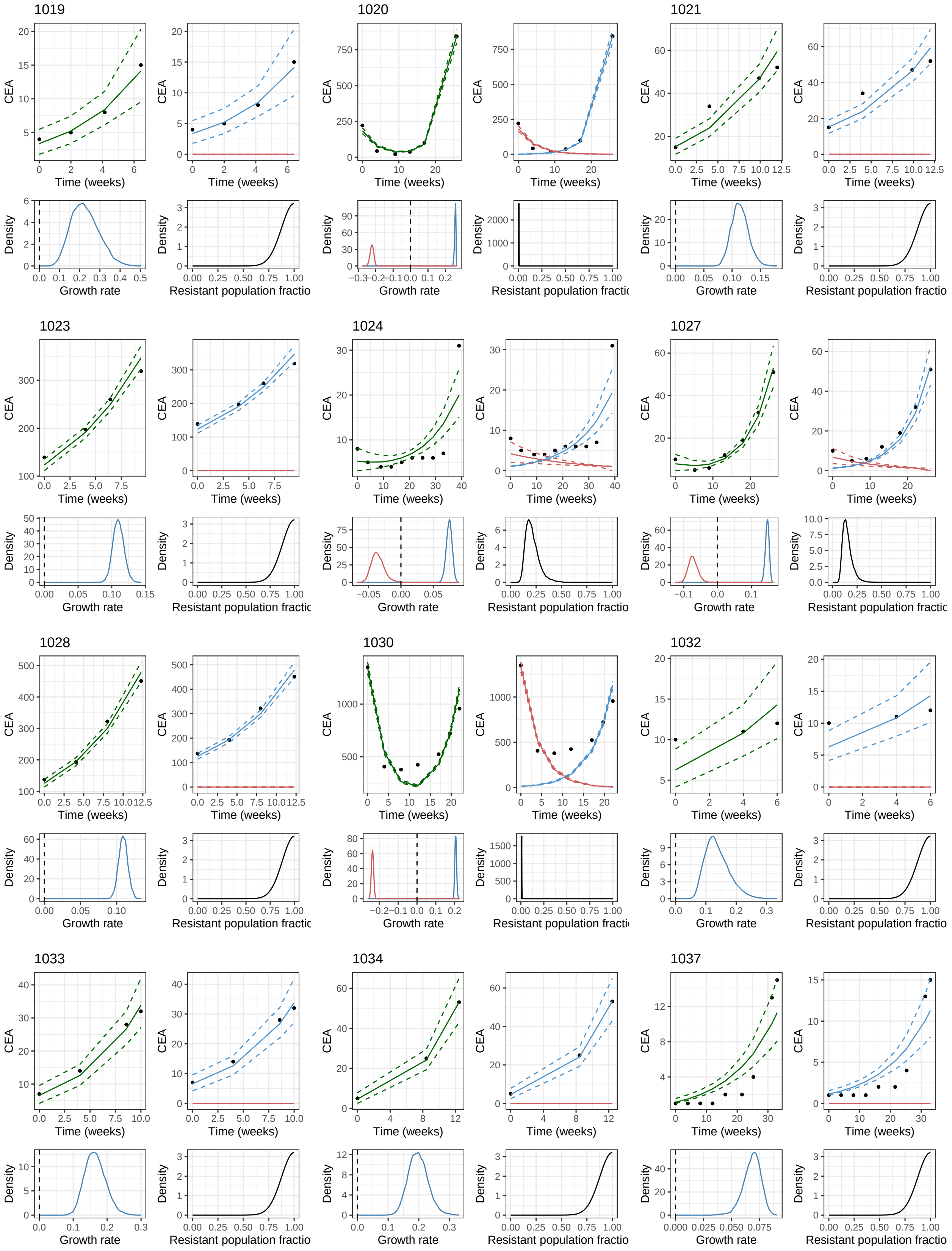
1019
1020
1021
1023
1024
1027
1028
1030
1032
1033
1034
1037
CEA
Time (weeks)
Density
Growth rate
Resistant population fractio

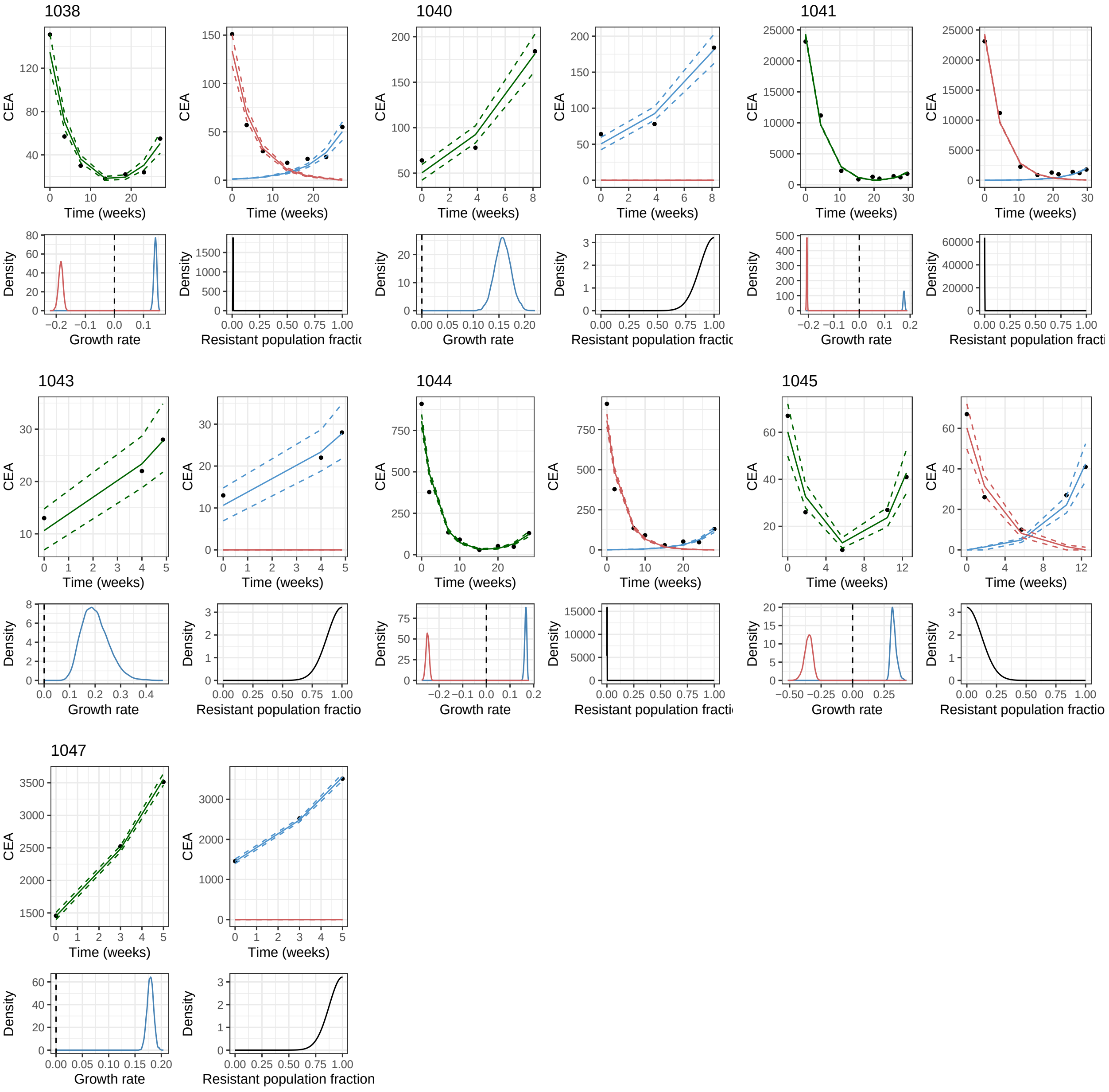

**Supplementary Figure S6.** Fit of the two-population model for each patient in the cohort from[2] during cetuximab treatment, based on CEA measurements. For each patient are reported: the dynamics fitted on the CEA measurements (in green), the dynamics fitted on the CEA measurements divided into resistant (blue) and sensitive population (red), the posterior densiry for the resistant and sensitive populations growth rates, and the posterior density for the resistant population fraction at the first observations (black).

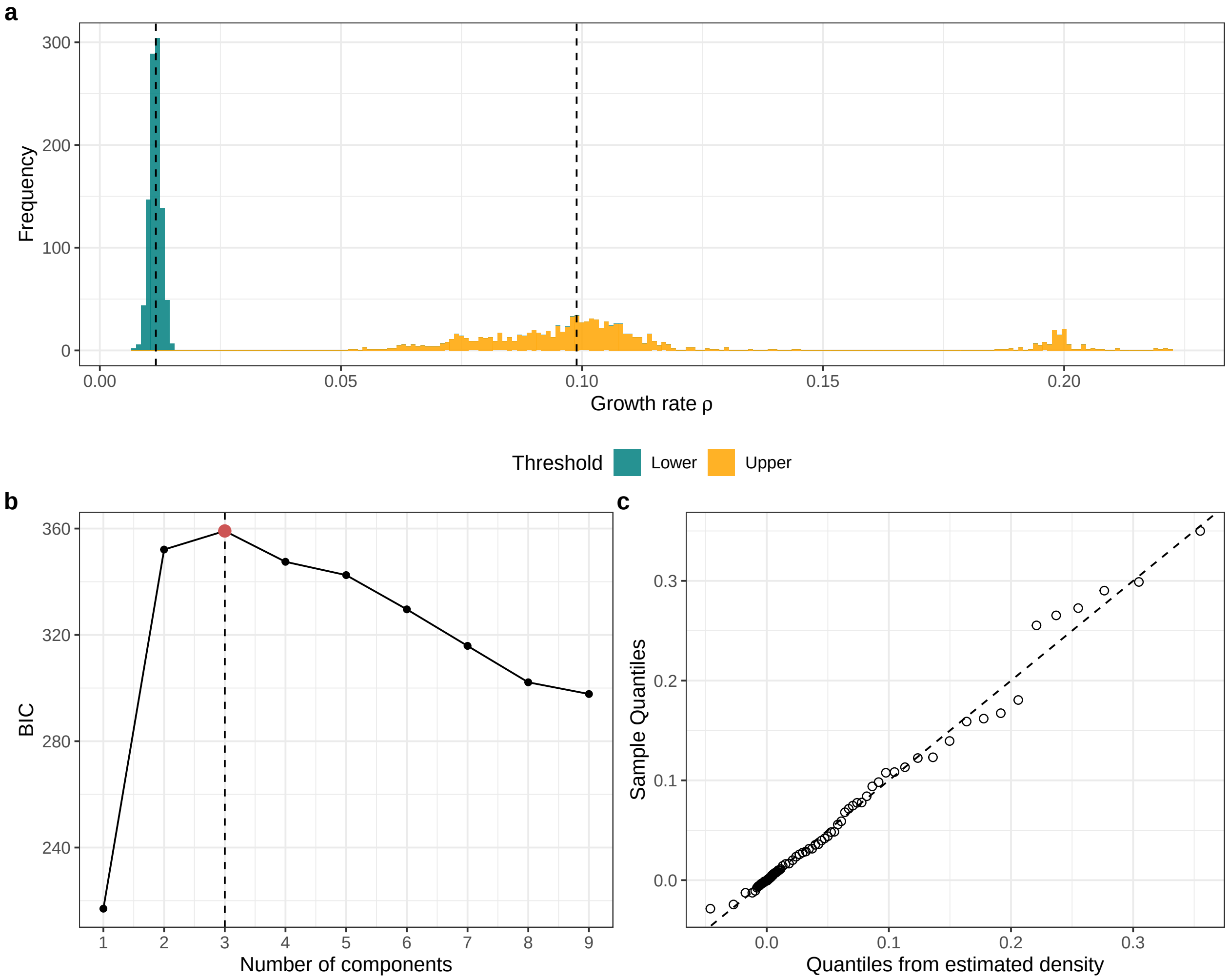

**Supplementary Figure S7. a.** Distribution for the thresholds stratifying patients into High, Medium, and Low growth rate classes in the tracerX cohort, obtained via bootstrap. **b.** Plot of BIC values used for choosing the optimal number of cluster, derived from the *mclust* package. **c.** QQ plot of sample quantiles versus the quantiles obtained from the inverse of the estimated cdf, obtained with the *mclust* package.

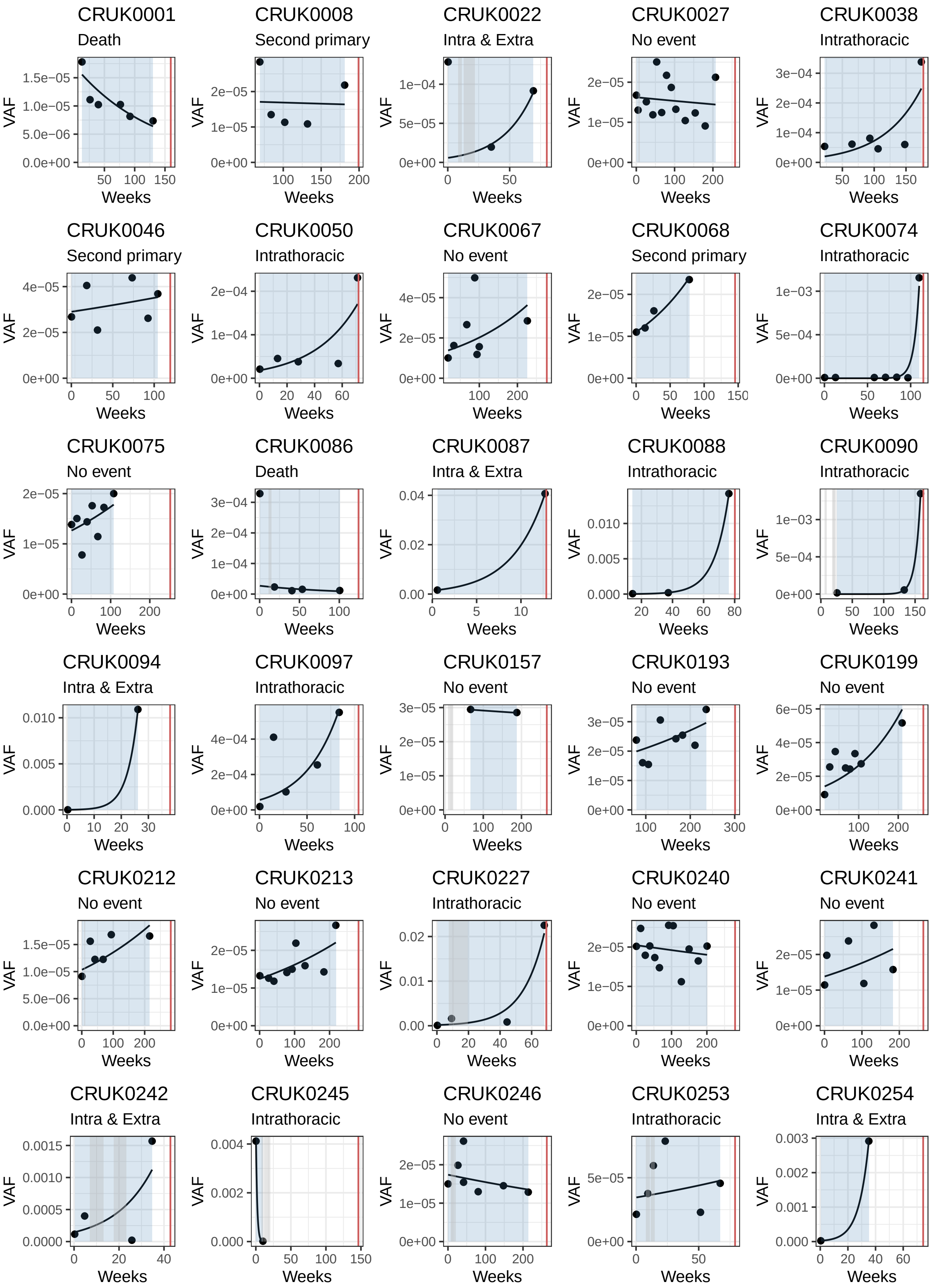
CRUK0001
Death
CRUK0008
Second primary
CRUK0022
Intra & Extra
CRUK0027
No event
CRUK0038
Intrathoracic
CRUK0046
Second primary
CRUK0050
Intrathoracic
CRUK0067
No event
CRUK0068
Second primary
CRUK0074
Intrathoracic
CRUK0075
No event
CRUK0086
Death
CRUK0087
Intra & Extra
CRUK0088
Intrathoracic
CRUK0090
Intrathoracic
CRUK0094
Intra & Extra
CRUK0097
Intrathoracic
CRUK0157
No event
CRUK0193
No event
CRUK0199
No event
CRUK0212
No event
CRUK0213
No event
CRUK0227
Intrathoracic
CRUK0240
No event
CRUK0241
No event
CRUK0242
Intra & Extra
CRUK0245
Intrathoracic
CRUK0246
No event
CRUK0253
Intrathoracic
CRUK0254
Intra & Extra
VAF
Weeks

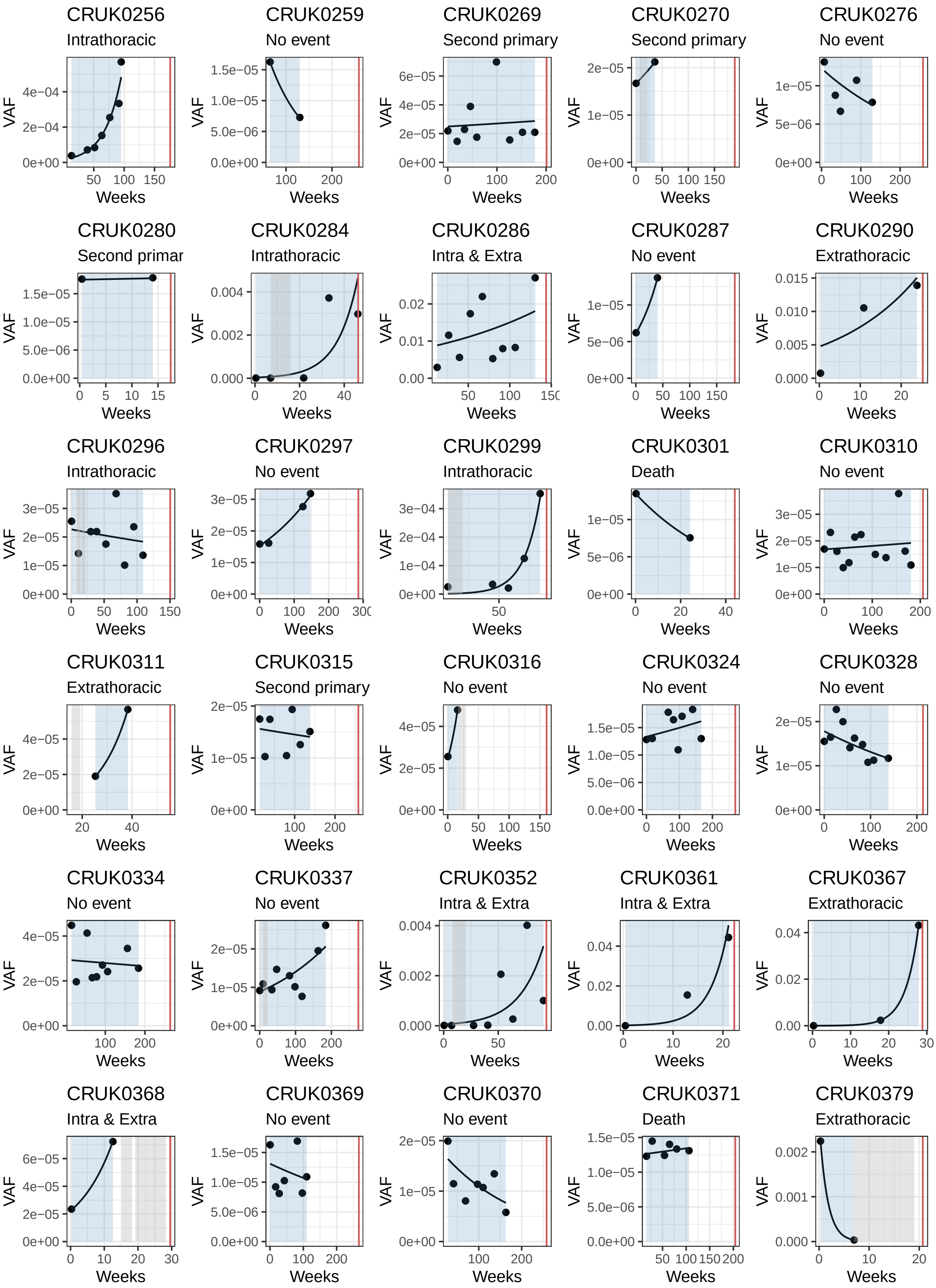
CRUK0256
Intrathoracic
CRUK0259
No event
CRUK0269
Second primary
CRUK0270
Second primary
CRUK0276
No event
CRUK0280
Second primar
CRUK0284
Intrathoracic
CRUK0286
Intra & Extra
CRUK0287
No event
CRUK0290
Extrathoracic
CRUK0296
Intrathoracic
CRUK0297
No event
CRUK0299
Intrathoracic
CRUK0301
Death
CRUK0310
No event
CRUK0311
Extrathoracic
CRUK0315
Second primary
CRUK0316
No event
CRUK0324
No event
CRUK0328
No event
CRUK0334
No event
CRUK0337
No event
CRUK0352
Intra & Extra
CRUK0361
Intra & Extra
CRUK0367
Extrathoracic
CRUK0368
Intra & Extra
CRUK0369
No event
CRUK0370
No event
CRUK0371
Death
CRUK0379
Extrathoracic
VAF
Weeks

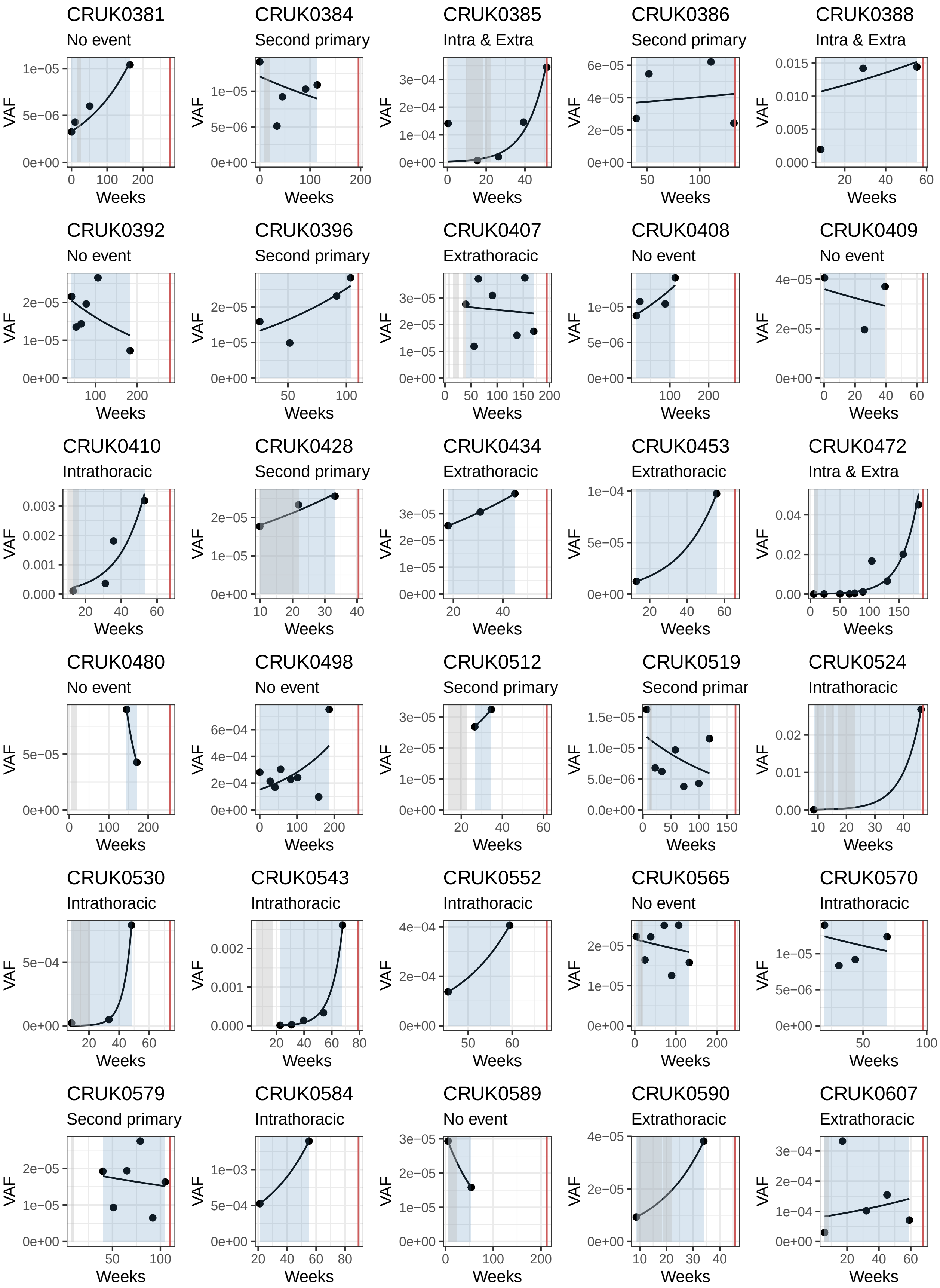
CRUK0381
No event
CRUK0384
Second primary
CRUK0385
Intra & Extra
CRUK0386
Second primary
CRUK0388
Intra & Extra
CRUK0392
No event
CRUK0396
Second primary
CRUK0407
Extrathoracic
CRUK0408
No event
CRUK0409
No event
CRUK0410
Intrathoracic
CRUK0428
Second primary
CRUK0434
Extrathoracic
CRUK0453
Extrathoracic
CRUK0472
Intra & Extra
CRUK0480
No event
CRUK0498
No event
CRUK0512
Second primary
CRUK0519
Second primar
CRUK0524
Intrathoracic
CRUK0530
Intrathoracic
CRUK0543
Intrathoracic
CRUK0552
Intrathoracic
CRUK0565
No event
CRUK0570
Intrathoracic
CRUK0579
Second primary
CRUK0584
Intrathoracic
CRUK0589
No event
CRUK0590
Extrathoracic
CRUK0607
Extrathoracic
VAF
Weeks

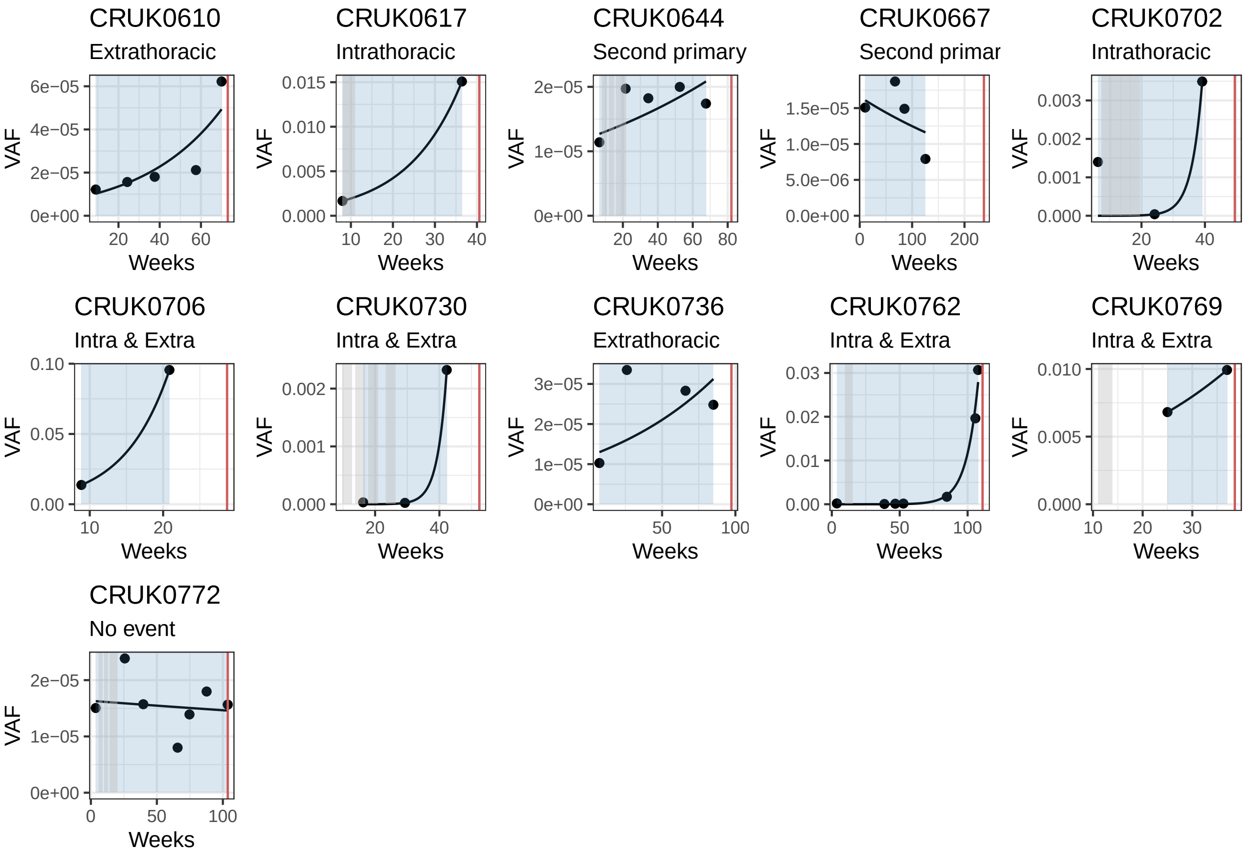

**Supplementary Figure S8.** Fit of patients in the tracerX cohort from[3]. Black dots represents positive VAF measurements after surgery, gray areas represents eventual treatment, and red line indicates date of final event (e.g relase, death, no event).

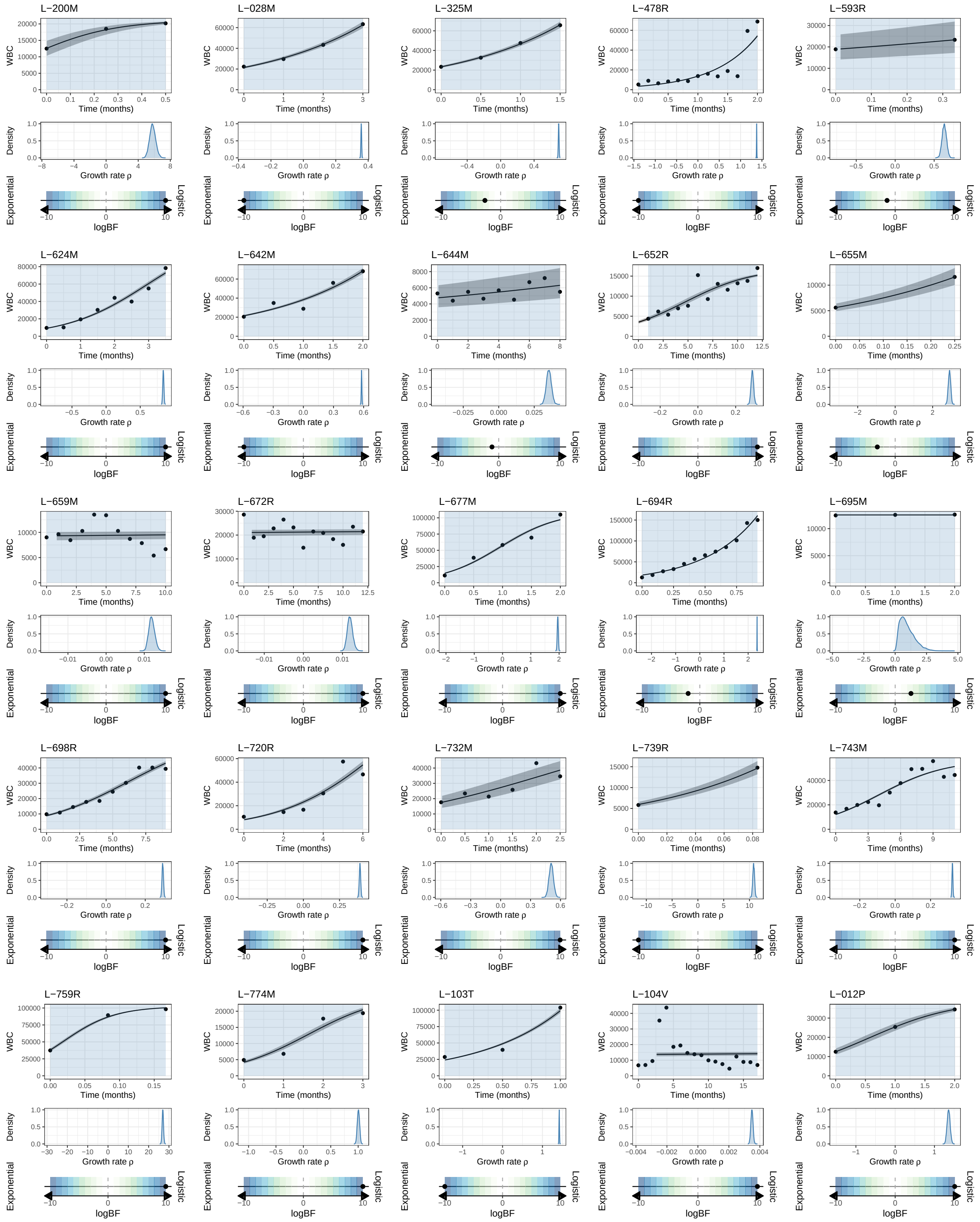
L−200M
L−028M
L−325M
L−478R
L−593R
L−624M
L−642M
L−644M
L−652R
L−655M
L−659M
L−672R
L−677M
L−694R
L−695M
L−698R
L−720R
L−732M
L−739R
L−743M
L−759R
L−774M
L−103T
L−104V
L−012P
WBC
Time (months)
Density
Growth rate ρ
Exponential
Logistic
logBF

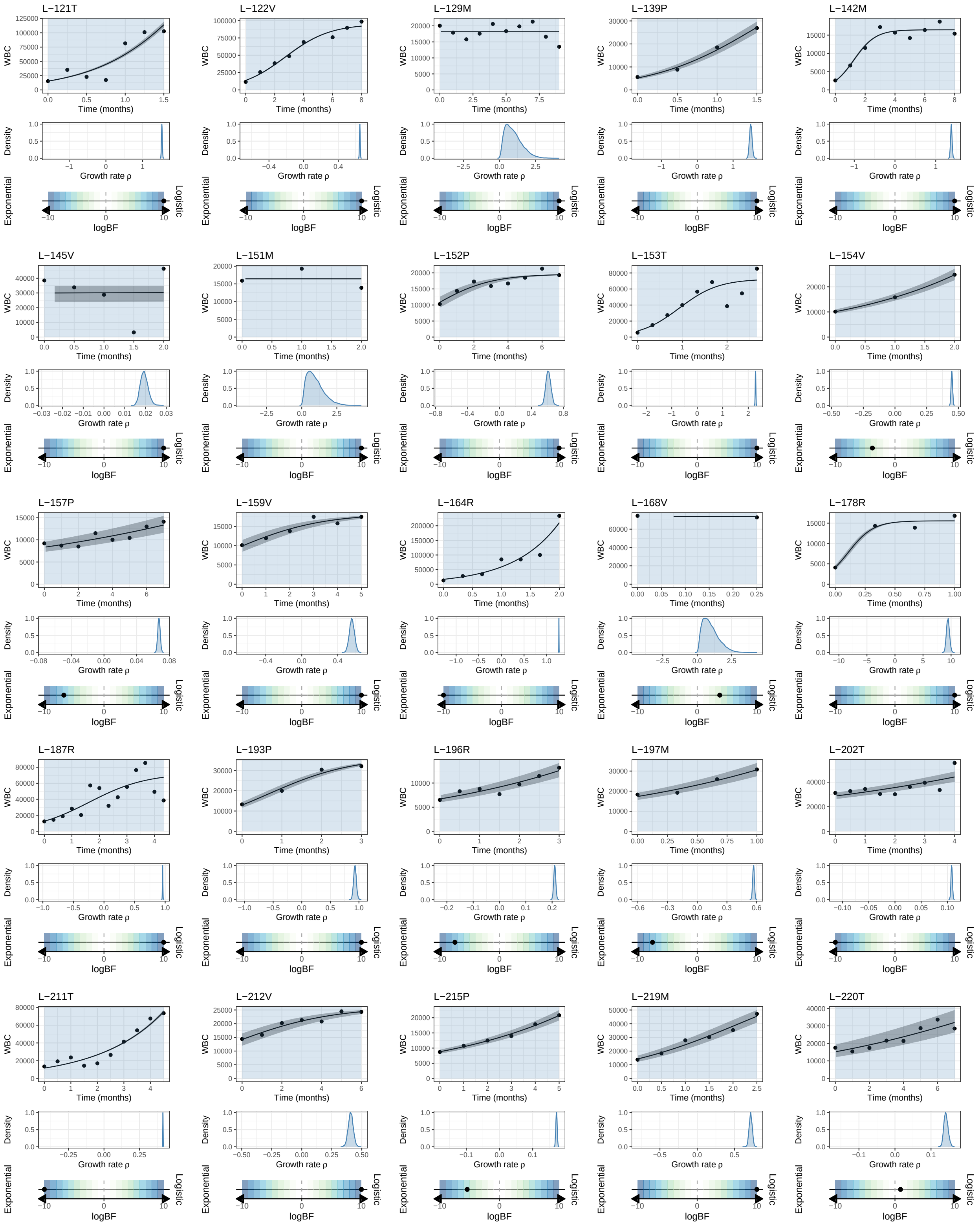
L−121T
L−122V
L−129M
L−139P
L−142M
L−145V
L−151M
L−152P
L−153T
L−154V
L−157P
L−159V
L−164R
L−168V
L−178R
L−187R
L−193P
L−196R
L−197M
L−202T
L−211T
L−212V
L−215P
L−219M
L−220T
WBC
Time (months)
Density
Growth rate ρ
Exponential
Logistic
logBF

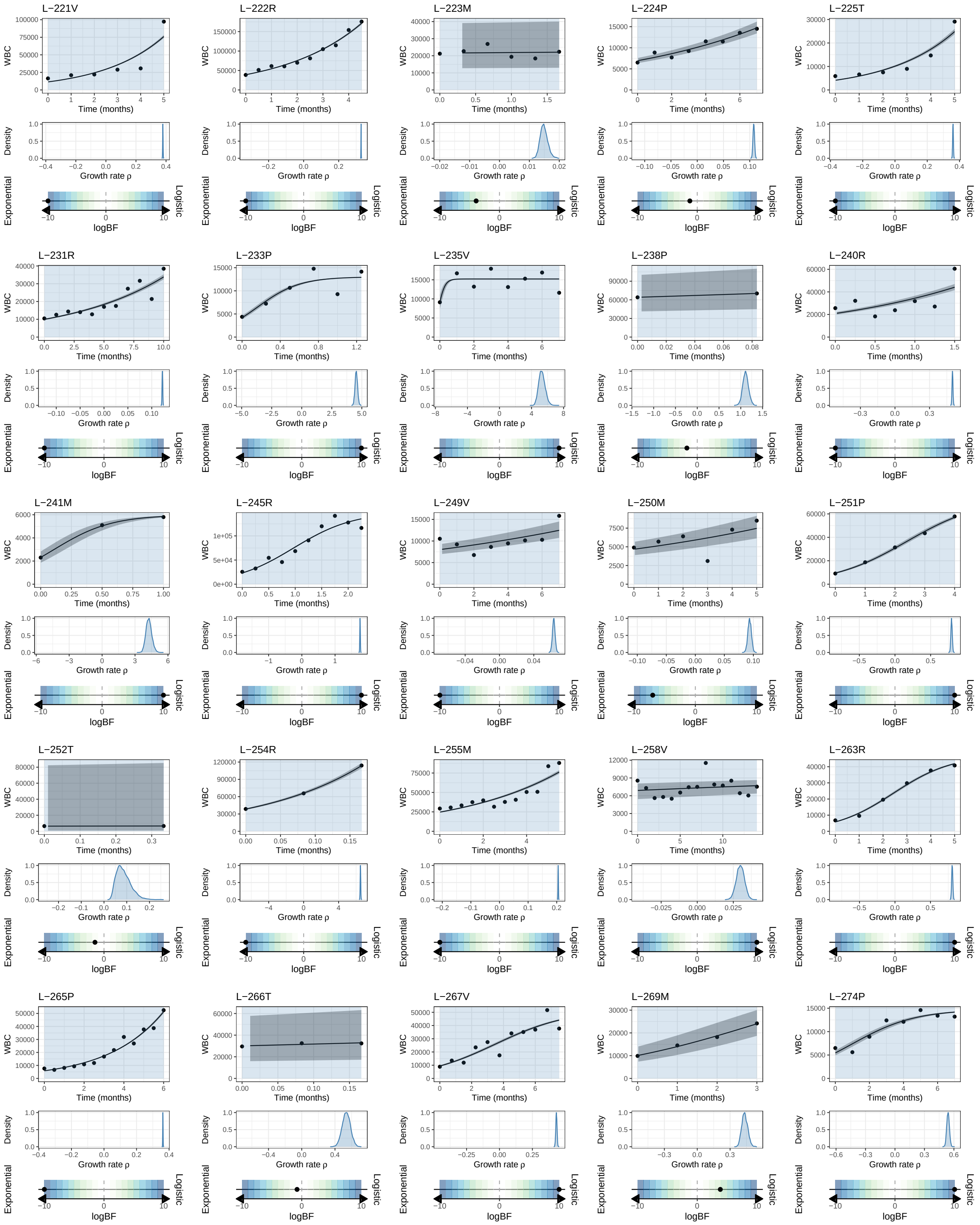
L−221V
L−222R
L−223M
L−224P
L−225T
L−231R
L−233P
L−235V
L−238P
L−240R
L−241M
L−245R
L−249V
L−250M
L−251P
L−252T
L−254R
L−255M
L−258V
L−263R
L−265P
L−266T
L−267V
L−269M
L−274P
WBC
Time (months)
Density
Growth rate ρ
Exponential
Logistic
logBF

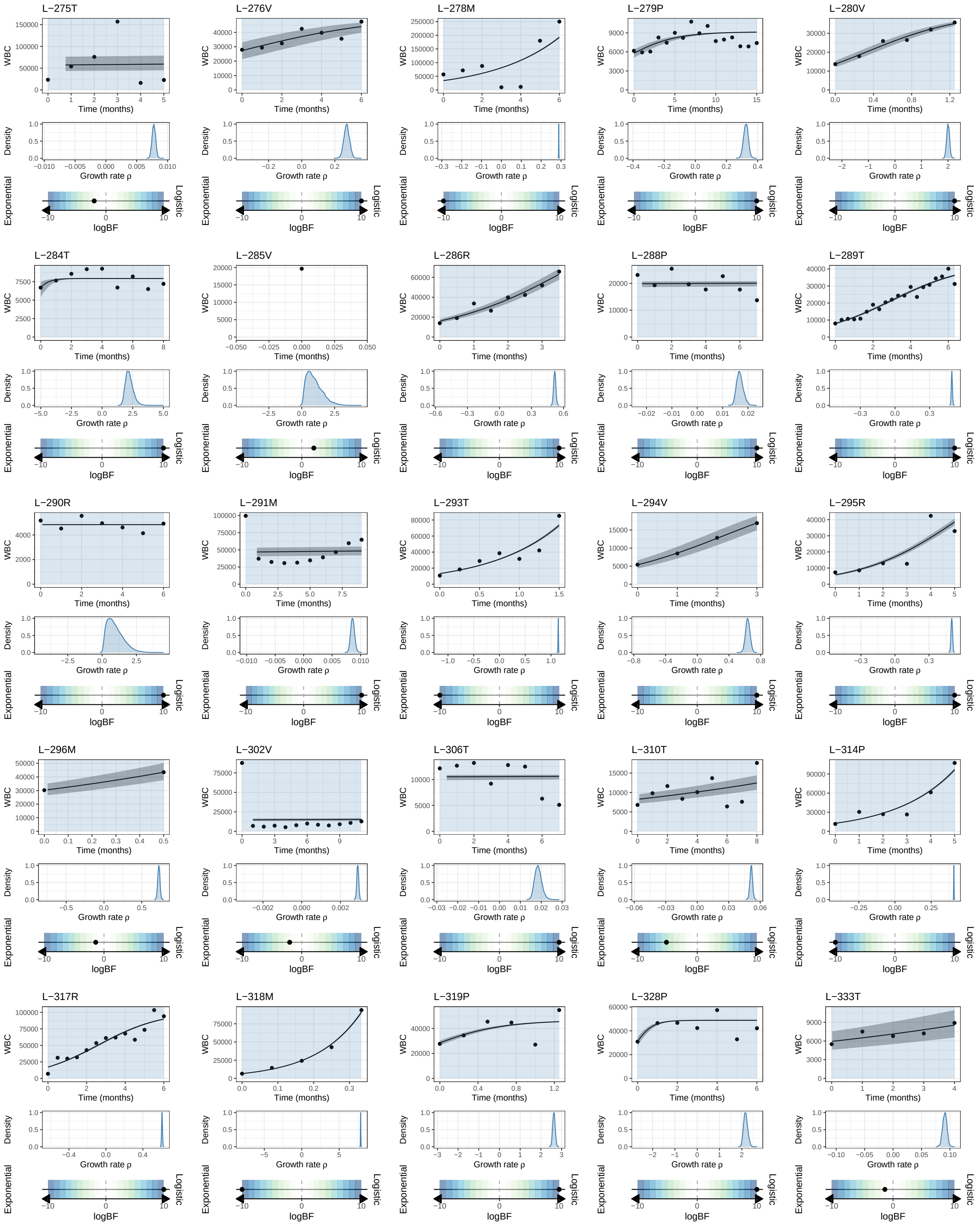
L−275T
L−276V
L−278M
L−279P
L−280V
L−284T
L−285V
L−286R
L−288P
L−289T
L−290R
L−291M
L−293T
L−294V
L−295R
L−296M
L−302V
L−306T
L−310T
L−314P
L−317R
L−318M
L−319P
L−328P
L−333T
WBC
Time (months)
Density
Growth rate ρ
Exponential
Logistic
logBF

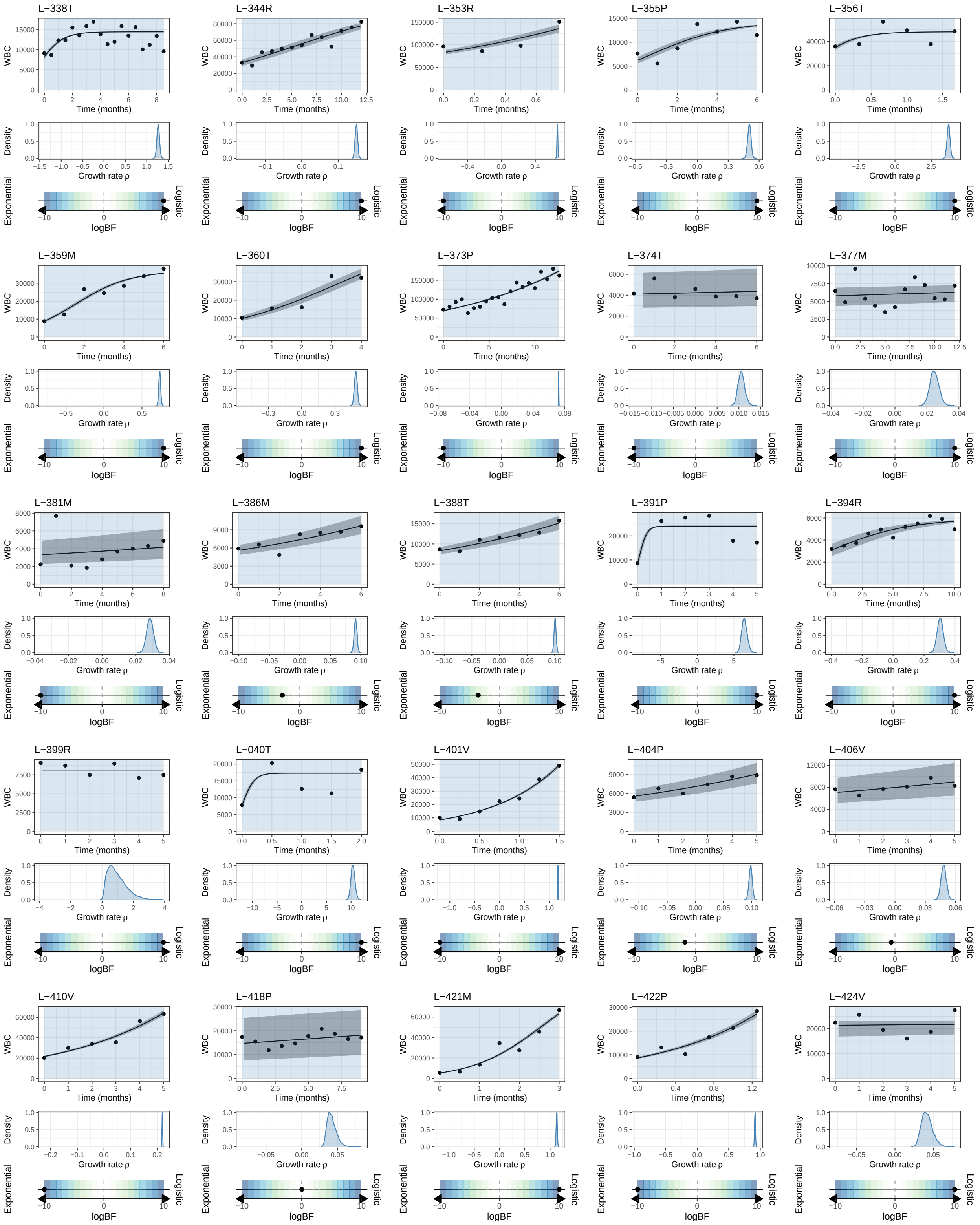
L−338T
L−344R
L−353R
L−355P
L−356T
L−359M
L−360T
L−373P
L−374T
L−377M
L−381M
L−386M
L−388T
L−391P
L−394R
L−399R
L−040T
L−401V
L−404P
L−406V
L−410V
L−418P
L−421M
L−422P
L−424V
WBC
Time (months)
Density
Growth rate ρ
Exponential
Logistic
logBF

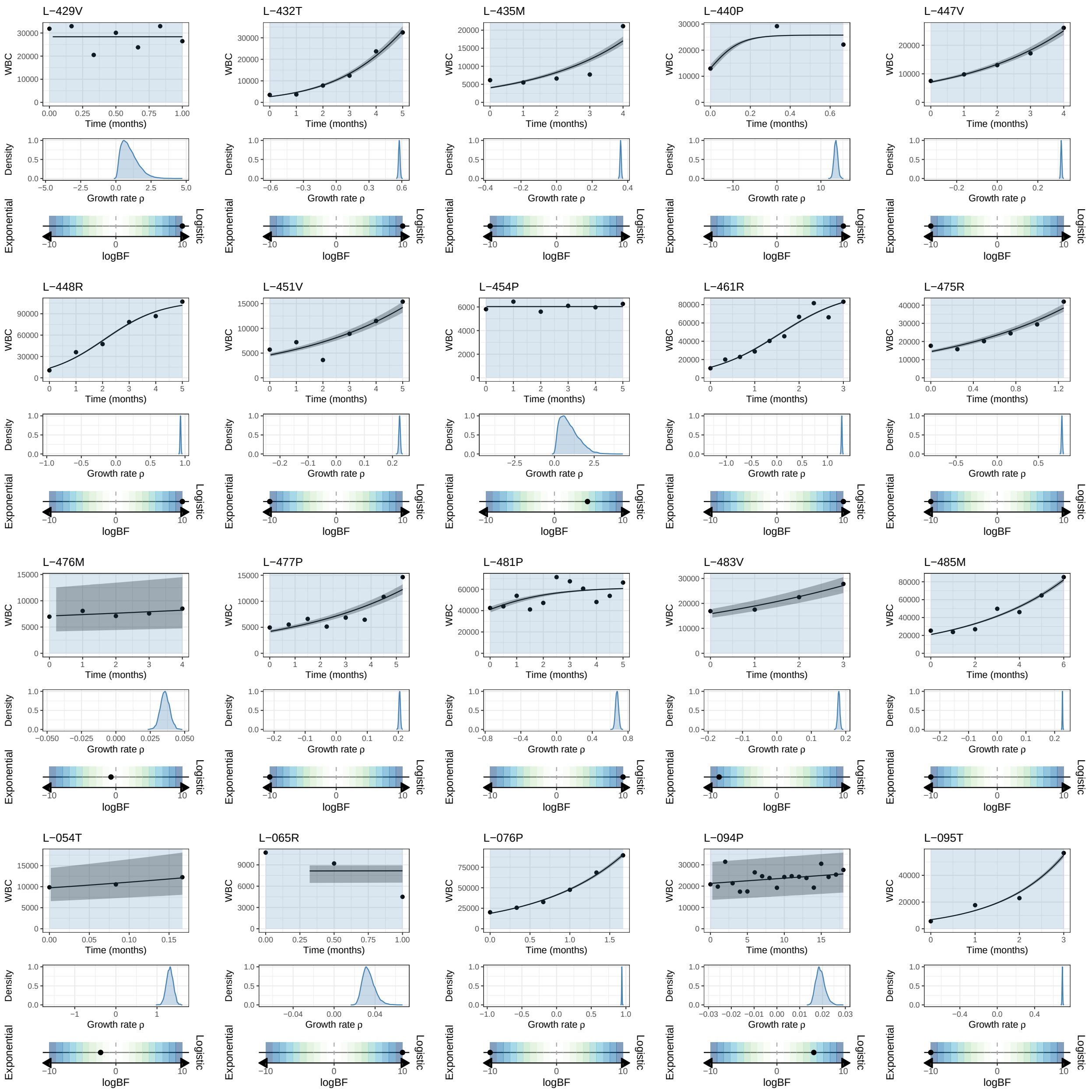

**Supplementary Figure S9.** Fit of patients in the CLL cohort. For each patient are reported the WBC measurements over time(black dots) and the biPOD fit, the posterior distribution for the inferred growth rate, and the bayes factor coming from the model selection.

### B. Supplementary Tables

| | Gene | High % | Medium % | Low % | $p_{value}$ | $p_{adj}$ |
|---|---|---|---|---|---|---|
| 1 | TP53 | 0.56 | 0.56 | 0.52 | 0.92 | 1.00 |
| 2 | KRAS | 0.38 | 0.30 | 0.29 | 0.80 | 1.00 |
| 3 | NF1 | 0.00 | 0.07 | 0.16 | 0.18 | 0.52 |
| 4 | KMT2D | 0.25 | 0.26 | 0.05 | 0.01 | 0.15 |
| 5 | CREBBP | 0.25 | 0.11 | 0.07 | 0.13 | 0.52 |
| 6 | KEAP1 | 0.12 | 0.15 | 0.10 | 0.92 | 1.00 |
| 7 | ARID1A | 0.12 | 0.00 | 0.07 | 0.19 | 0.52 |
| 8 | EGFR | 0.00 | 0.11 | 0.09 | 0.46 | 0.89 |
| 9 | PIK3CA | 0.12 | 0.07 | 0.05 | 0.66 | 1.00 |
| 10 | STK11 | 0.00 | 0.11 | 0.09 | 0.49 | 0.89 |
| 11 | ATM | 0.06 | 0.04 | 0.05 | 1.00 | 1.00 |

**Table 1.** Results of contingency table analysis for 12 genes of interest in the tracerX cohort[3]. The p-values have been corrected with the Benjamini-Hochberg procedure. Additionally, the percentages of patient in a given class (i.e. High, Medium, and Low) with a given mutation are also reported.

| | Variable | 1 | 2 | 3 | $p_{value}$ | $p_{adj}$ |
|---|---|---|---|---|---|---|
| 1 | TNM stage | I | II | III | 0.33 | 0.46 |
| 2 | Smoking | Below first quantile | Above third quantile | In between | 0.37 | 0.46 |
| 3 | Histology | Squamous cell carcinoma | Invasive adenocarcinoma | Other | 0.28 | 0.46 |
| 4 | Gender | Male | Female | | 0.80 | 0.80 |
| 5 | Age | Younger than 64 | Older than 75 | In between | 0.08 | 0.41 |

**Table 2.** Results of contingency table analysis for 5 clinical variables of interest in the tracerX cohort[3]. The p-values have been corrected with the Benjamini-Hochberg procedure. Additionally, the groups used to stratify the patienst with respect to the variables of interest are also reported.

| | Variable | % in treated | % in untreated | $p_{value}$ | $p_{adj}$ |
|---|---|---|---|---|---|
| 1 | Unmutated IGHV | 0.532 | 0.182 | 1.00e-03 | 6.00e-03 |
| 2 | Tri12 | 0.156 | 0.121 | 0.474 | 0.566 |
| 3 | Del13 | 0.608 | 0.545 | 0.518 | 0.566 |
| 4 | Del11 | 0.088 | 0.00 | 2.40e-02 | 0.072 |
| 5 | Del17 | 0.114 | 0.079 | 0.566 | 0.566 |
| 6 | Bimodal CD49d | 0.278 | 0.379 | 0.213 | 0.426 |

**Table 3.** Results of contingency table analysis for 6 clinical variables of interest in the CLL cohort. The p-values have been corrected with the Benjamini-Hochberg procedure. Additionally, the percentages of treated with and untreated patients with the considered variables are also reported.

| | Variable | 1 | 2 | 3 | $p_{value}$ | $p_{adj}$ |
|---|---|---|---|---|---|---|
| 1 | $\rho$ | First quantile | Third quantile | In between | 5.00e-04 | 1.00e-03 |
| 2 | Growth pattern | Exponential | Logistic | | 0.0975 | 0.097 |

**Table 4.** Results of contingency table analysis for the 2 main parmaters inferred by biPOD in the CLL cohort. The p-values have been corrected with the Benjamini-Hochberg procedure.

| Task | Set | $N_{pos}$ | $N_{neg}$ | Metric | Value |
|---|---|---|---|---|---|
| | | | | Sensitivity | 0.612 |
| | | | | Specificity | 0.955 |
| | | | | Pos Pred Value | 0.937 |
| | Train | 49 | 45 | Neg Pred Value | 0.693 |
| | | | | Precision | 0.937 |
| | | | | Recall | 0.612 |
| | | | | F1 | 0.741 |
| | | | | Sensitivity | 0.650 |
| | | | | Specificity | 0.904 |
| | | | | Pos Pred Value | 0.866 |
| Need of treatment | Test | 20 | 21 | Neg Pred Value | 0.731 |
| | | | | Precision | 0.866 |
| | | | | Recall | 0.650 |
| | | | | F1 | 0.743 |
| | | | | Sensitivity | 0.623 |
| | | | | Specificity | 0.939 |
| | | | | Pos Pred Value | 0.915 |
| | Whole | 69 | 66 | Neg Pred Value | 0.705 |
| | | | | Precision | 0.915 |
| | | | | Recall | 0.623 |
| | | | | F1 | 0.741 |

**Table 5.** Performance metrics in the need or treatment prediction for the CLL cohort, divided by test and train sets.

| Task | Set | $N_{pos}$ | $N_{neg}$ | Metric | Value |
|---|---|---|---|---|---|
| | | | | Sensitivity | 0.821 |
| | | | | Specificity | 0.750 |
| | | | | Pos Pred Value | 0.958 |
| | Train | 28 | 4 | Neg Pred Value | 0.375 |
| | | | | Precision | 0.958 |
| | | | | Recall | 0.821 |
| | | | | F1 | 0.885 |
| | | | | Sensitivity | 0.727 |
| | | | | Specificity | 0.25 |
| | | | | Pos Pred Value | 0.727 |
| Detect treatment within 18 months | Test | 11 | 4 | Neg Pred Value | 0.25 |
| | | | | Precision | 0.727 |
| | | | | Recall | 0.727 |
| | | | | F1 | 0.727 |
| | | | | Sensitivity | 0.795 |
| | | | | Specificity | 0.5 |
| | | | | Pos Pred Value | 0.886 |
| | Whole | 39 | 8 | Neg Pred Value | 0.333 |
| | | | | Precision | 0.886 |
| | | | | Recall | 0.795 |
| | | | | F1 | 0.838 |

**Table 6.** Performance metrics in the detection of treatment within 18 months for the CLL cohort, divided by test and train sets.

#### C. Supplementary Material

We use the following notation:

- $\boldsymbol{X} = \{(x_i, y_i) \mid i = 1, \dots, N\}$, set of tumour measurements $y_i$ (of multiple kinds) at time $x_i$, with $N$ observations;
- $\boldsymbol{\Theta}$, tumour parameters that we infer and that describe the dynamics;
- $\rho_j$, growth rate of a population in a time window $j$
- $n_0$, starting size of a population (i.e. the size of the population at time $x = 0$)
- $t_0$, instant of birth of a population
- $L$, carrying capacity (i.e. the maximum size of a given population) of a logistic growth

##### C.1. Multi-step logistic and exponential growth models

A general growth process can be seen as a continuous-time Markov process[4,5], where $y(x) \in \mathbb{N}_0$ represents the number of individuals in the population of cancerous cell at time $t$, and $t_0$ indicates the instant of time in which the first cancerous cell is born, i.e. $y(x = t_0) = 1$. The transition rates for such problem are defined as:

$$\alpha_n(x) = \lim_{h \to 0^+} \frac{1}{h} P[y(x+h) = n+1 \mid y(x) = n] \tag{1}$$

$$\beta_n(x) = \lim_{h \to 0^+} \frac{1}{h} P[y(x+h) = n-1 \mid y(x) = n] \tag{2}$$

where $P(\cdot)$ denotes the probability that a population of size $n$ will increase/decrease by one individual in the time interval $h$. Here $\alpha_n(t)$ and $\beta_n(t)$ are both positive functions and indicate the birth and death rates of the population when $y(x) = n$, respectively. Similarly, one can denote with $\alpha(t)$ and with $\beta(t)$ the individual birth and death rate at time $t$. Assuming that $y(0) = n_0$ with $n_0 \in \mathbb{N}$ and given $m \in \mathbb{N}_0$, one is interested in computing

$$P_{n_0,m}(x) = P[y(x) = m \mid y(0) = n_0], \quad t \geq 0 \tag{3}$$

which represents the probability that the population reaches a population size equal to $m$ at time $x$ conditioned on the population size at $x = 0$. Quite trivially, $P_{0,m} = 0 \; \forall m > 0$, meaning that 0 is an absorbing state for the Markov process. The probability generating function for this process is

$$G(z, x) = \sum_{m=0}^{\infty} P_{n_0,m}(x) z^m \tag{4}$$

and, as shown in[6], can be rewritten as

$$G(z, x) = [1 - (z-1)[(z-1)\phi(x) - \psi(x)]^{-1}]^{n_0}\,, \tag{5}$$

where the quantities $\phi(x)$ and $\psi(x)$ depend on the functional form of the individuals birth and death rates:

$$\psi(x) = \exp\left\{-\int_0^x (\alpha(\tau) - \beta(\tau))d\tau\right\} \tag{6}$$

$$\phi(x) = \int_0^x \alpha(\tau)\psi(\tau)d\tau\,. \tag{7}$$

One can then compute the expectation and the variance of $y(x)$ by using the probability generating function and the fact that $E[x] = G'(1, x)$ and that $\text{Var}(x) = G''(1, x) + G'(1, x) - [G'(1, x)]^2$. After some standard algebra, one obtains

$$\mathbb{E}_{n_0}(x) = \mathbb{E}[y(x) \mid n_0] = \frac{n_0}{\psi(x)} \tag{8}$$

$$\text{Var}_{n_0}(x) = n_0 \frac{\psi(x) + 2\phi(x) - 1}{\psi^2(x)}\,. \tag{9}$$

###### *Exponential growth*

The first growth pattern we consider can be expressed through a simple yet ubiquitous differential equation:

$$\frac{dy(x)}{dx} = \rho y(x) \tag{10}$$

where $\rho \geq 0$ represents the growth rate. The solution of this equation yields

$$y(x) = n_0 e^{\rho x}\,, \tag{11}$$

indicating that a population starting from $n_0$ will evolve exponentially over time with no upper bounds. In such growth pattern, the transition rates can be written as

$$\alpha_n = y\alpha(x), \quad \beta_n = y\beta(x) \tag{12}$$

It is natural to observe that the birth and death rates are linear with respect to $n$, meaning that the probability of a birth (or death) happening in an infinitesimal time-interval is proportional to the previous size of the population. Let $\rho(x) = \alpha(x) - \beta(x)$ be the individual net growth rate at time $x$ and assume that $\rho(x)$ is a piece-wise constant function, representing different net growth rates for a multi-step growth process. Without loss of generality, consider a process with a single known time-point $t_1$ in which the net growth rate changes from $\rho_1$ to $\rho_2$; in such case the form of $\psi(x)$ will be

$$\psi(x) = \exp\{-\rho_1 x\}\,, \qquad x \leq t_1 \tag{13}$$

$$\psi(x) = \exp\{-\rho_1 t_1\}\exp\{-\rho_2(x - t_1)\}\,, \qquad x > t_1 \tag{14}$$

It is trivial to see that this can be generalised to any number of time-points, i.e. to any number of different growth rates, and that the expected mean conditioned on $n_0$ can be computed as

$$\mathbb{E}_{n_0}(x) = n_0 \exp\{\rho_1 x\}\,, \qquad x \leq t_1 \tag{15}$$

$$\mathbb{E}_{n_0}(x) = n_0 \exp\{\rho_1 t_1\}\exp\{\rho_2(x - t_1)\}\,, \qquad x > t_1 \tag{16}$$

##### *Logistic growth*

The logistic model is a generalisation of the exponential one which takes into account the possibility that the maximum population size is constrained. In this case we define the carrying capacity $L$ as such upper bound, which can be caused by many environmental factors such as the availability of resources or space. The generalised differential equation becomes

$$\frac{dy(x)}{dx} = \rho\left[1 - \frac{y(x)}{L}\right]y(x)\,, \tag{17}$$

where the second term in squared parentheses modulates the effective growth rate affecting the population: a bigger population size leads to a slower growth. By letting $L \to \infty$, the logistic equation tends to the exponential one. This equation can easily be solved by separation of variables and which yields as solution

$$y(x) = \frac{n_0 L}{n_0 + (L - n_0)\exp\{-\rho x\}}\,, \tag{18}$$

and one can also merge the two terms of the differential equation and substitute them with a time-dependent growth rate:

$$\frac{dy(x)}{dx} = \nu(x)y(x)\,. \tag{19}$$

Even in this case, the solution for $\nu(t)$ is straightforward

$$\nu(x) = \frac{\rho(L - n_0)}{(L - n_0) + n_0\exp\{\rho x\}}. \tag{20}$$

Now, similarly as what has been done in the exponential case where indicated the net growth rate per capita and was defined as a piece-wise constant function, $\nu(x)$ once again represents the net growth rate per capita which depends on the initial conditions, the carrying capacity $L$ and the functional form of $\rho$. Once again, we assume $\rho$ to be piece-wise constant and we denote as to its changing points. Hence, we are modelling growth patterns in which the net growth rate changes over the time not only depending on the population size, but also on the value of the piece-wise constant function $\rho(t)$. Now, without loss of generality let's consider a single changing point $t_1$ so that one can first compute $\psi(x)$ and the the expected mean conditioned on the starting conditions (i.e. the value of $n_0$)

$$\mathbb{E}_{n_0}(x) = \frac{n_0 L}{n_0 + (L - n_0)\exp\{-\rho_1 x\}}, \qquad x \le t_1 \tag{21}$$

$$\mathbb{E}_{n_0}(x) = \frac{n_1 L}{n_1 + (L - n_1)\exp\{-\rho_2 (x - t_1)\}}, \qquad x > t_1 \tag{22}$$

where we define $n_1$ as $\mathbb{E}_{n_0}(x = t_1)$.

***Start of the process and initial size***

We discussed so far the situation in which $y(0) = n_0$ since the most common situation is that the first observation of a growth process happens much later than the actual start of the process. Nonetheless, one could also be interested in obtaining an estimate, or at least an upper bound, for

$$t_0 = \min\{x \mid y(x) = 1\}.$$

This can be done by simply shifting by $t_0$ the values of all the instant of times (i.e. with the change of variables $x' = x - t_0$). The only modification to the model is that now $y(x' = 0) = 1$. Therefore, the equation for the exponential growth becomes

$$\mathbb{E}_{t_0}(x) = \exp\{\rho_1 x'\} = \exp\{\rho_1 (x - t_0)\} \tag{23}$$

while the equation for the logistic growth becomes

$$\mathbb{E}_{t_0}(x) = \frac{L}{1 + (L-1)\exp\{-\rho_1 x'\}} = \frac{L}{1 + (L-1)\exp\{-\rho_1 (x - t_0)\}} \tag{24}$$

Depending on the parameter of interest or the nature of the process, either $n_0$ or $t_0$ can be learned (e.g. $t_0$ cannot be learned if the population observed is dying).

#### C.2. Change points detection via piece-wise linear regression

In general, the change-points $t_i$ defining the intervals of time of the piece-wise functions that we consider might not be known and, therefore, a method to infer them is needed. The main assumption we make is that the underlying growth can be described by an exponential growth and we can therefore see the task as finding the best knots for piece-wise linear regression. In fact, by assuming $K$ to be then number of breakpoints and by taking the logarithm of Eq. 15 one obtains

$$\log \mu(x) = \beta_0 + \beta_1 x + \sum_{i=1}^{K} \beta_{i+1}(x - t_i) I(x, t_i), \tag{25}$$

where $\mu(x) \equiv \mathbb{E}_{n_0}(x)$ is the mean given in Eq.15, $\beta_0$ represents $\log n_0$, and the $\beta_i$ with $i \ge 1$ represent the $\rho_i$ in Eq. 15. The equation above can also be represented in the following matrix form

$$A = \begin{bmatrix} 1 & x_1 & \max(0, x_1 - t_1) & \max(0, x_1 - t_2) & \dots & \max(0, x_1 - t_K) \\ 1 & x_2 & \max(0, x_2 - t_1) & \max(0, x_2 - t_2) & \dots & \max(0, x_2 - t_K) \\ \vdots & \vdots & \vdots & \vdots & \dots & \vdots \\ 1 & x_n & \max(0, x_n - t_1) & \max(0, x_n - t_2) & \dots & \max(0, x_n - t_K) \end{bmatrix} \tag{26}$$

Now, given a set of proposed breakpoints $\{t_1, \dots, t_K\}$, we can find the $\boldsymbol{\beta}$ vector that minimizes the sum of squared residuals between $\log y$ and $A\boldsymbol{\beta}$. We use the differential evolution algorithm from the *DEoptim*[7] package to find the optimal set of breakpoints $\{t_1, \dots, t_K\}$, i.e. the set yielding the estimated $\boldsymbol{\beta}$ vector with the lowest sum of squared residuals. This procedure is executed iteratively for each value $K$ of possible interest. Then, each set of proposed breakpoints for each $K$ of interest are used to fit the model described in Section C.1 and the best number of breakpoints $K^*$ is selected through approximate leave-one-out cross-validation[8].

#### C.3. Nested growth models for competing populations

A typical scenario is the one in which a tumor is treated, its mass shrinks and it later regrows. In this case, the observed dynamics is U-shaped. For such cases, we assume the presence of two populations $s$ and $r$; the first is sensitive to the treatment, whereas the second one is resistant to it. This means that a positive and a negative growth rates will be exhibited by the two populations, respectively. The sensitive population is assumed to be present at the start of treatment with the initial condition $y_s(x_1) = n_s$ and is governed by the differential equation

$$\frac{dy_s(x)}{dx} = -\rho_s(x). \tag{27}$$

For the second population it is assumed that the first resistant cell is born at $x = t_r$. Therefore, the differential equation governing the dynamics of the resistant population will be

$$\frac{dy_r(x)}{dx} = u(x - t_r)\rho_r y_r, \tag{28}$$

where the Heaviside function $u(x - t_r)$ is defined as

$$u(x - t_r) = \begin{cases} 0, & x \le t_r \\ 1, & x > t_r \end{cases} \tag{29}$$

Trivially, the observed counts $y(x)$ will be the sum of the two populations and can be obtained analytically by integrating the two equations above and summing them. For $x \ge 0$ this gives

$$\mathbb{E}[\underbrace{y_s(x) + y_r(x)}_{y(x)} \mid \mathbf{\Theta}] = n_s \exp\Big[-\rho_s x\Big] + u(x - t_r)\exp\Big[\rho_r(x - t_r)\Big] \tag{30}$$

or, equivalently

$$\mathbb{E}[\underbrace{y_s(x) + y_r(x)}_{y(x)} \mid \mathbf{\Theta}] = u(t_s - x)\exp\Big[-\rho_s(x - t_s)\Big] + u(x - t_r)\exp\Big[\rho_r(x - t_r)\Big] \tag{31}$$

where $t_s$ is the instant of time in which the sensitive population disappears (also called extinction time $t_e$).

##### C.4. Likelihood and inference

For all the presented models and for a given set of observations $X = \{(x_n, y_n)\}$ we consider each of them as drawn from a Poisson process with parameter equal to the expected value of the population $\mathbb{E}_y(x)$ where the analytical form of the expected value will depend on the considered task. This means that for a given task with parameters $\mathbf{\Theta}$, the likelihood will be computed as

$$\mathcal{L}(X \mid \mathbf{\Theta}) = \prod_{n=1}^{N} \text{Poisson}(y_n \mid \mathbb{E}[y(x_n) \mid \mathbf{\Theta}]) \tag{32}$$

The inference is carried out via MCMC methods[9,10] and/or variational inference[11–13], two of the most commonly used methods in probabilistic inference. The first generates sequences of samples that represents the target distibution (i.e. the posterior distirbution for the parameters of interest) and which can then be used to estimate quantities of interest such as means, variances and probabilities from the obtained empirical distribution. On the other hand, VI formulates the problem as an optimization task. Instead of sampling, VI chooses a familiy of distributions (e.g. Gaussian) and finds the best approximation to the true target posterior.

###### *Probabilistic grafical models*

The probabilistic graphical models for the tree different tasks C.1-C.3 will be presented and the prior distributions employed in each of them will be described in detail.

Given the trajectory $X = \{(x_n, y_n)\}$ of a population divided in $J$ time windows, the parameters of interest will be the $J$ growth rates $\rho_j$, either the instant of time in which the population was born $t_0$ or the initial population size $n_0$ and, if the population is undergoing a logistic growth, the carrying capacity $L$. The priors for those values are

$$\begin{aligned} \rho_j &\sim \text{Normal}(0, 1) \\ t_0 &\sim \text{TruncatedNormal}(x_1, 100, -\infty, x_1) \\ n_0 &\sim \text{TruncatedNormal}(y_1, y1/20, 0, \infty) \\ L &\sim \text{TruncatedNormal}(\max(x_n), \max(x_n), 0.85 \cdot \max(x_n), \infty) \end{aligned}$$

were a random variable $X$ follows a truncated normal distibution with parameters $(\mu, \sigma, a, b)$ if it follows a normal distribution with mean $\mu$ and variance $\sigma^2$ with the additional constraint that $a \le X \le b$. We choose a standard normal for the growth rates, a broad normal for $t_0$ with the constraint that $t_0$ must be smaller or equal to the time of the first observation, a normal biased by the first observed population size $y_1$ for the initial size $n_0$, and a broad normal for the carrying capacity with the constraint that $L$ must be greater or equal to the largest observation. The joint therefore is

$$
\begin{aligned}
p(\boldsymbol{X}, \boldsymbol{\Theta}) = & \left( \prod_{n=1}^{N} \text{Poisson}(y_n \mid \mathbb{E}[y(x_n) \mid \boldsymbol{\Theta}]) \right) \cdot \left( \prod_{j=1}^{J} \mathcal{N}(\rho_i \mid 0, 1) \right) \\
& \underbrace{\mathcal{N}_{\text{trunc}}(t_0 \mid x_1, 100, -\infty, x_1)}_{\text{if } t_0} \cdot \underbrace{\mathcal{N}_{\text{trunc}}(n_0 \mid y_1, y_1/20, 0, \infty)}_{\text{if } n_0} \\
& \underbrace{\mathcal{N}_{\text{trunc}}(L \mid \max(x_n), \max(x_n), 0.85 \cdot \max(x_n), \infty)}_{\text{if Logistic}}
\end{aligned}
\tag{33}
$$

The parameters of interested of the second task are the same as the one discussed for the previous task, with the only exception of the breakpoints $t_j$, that were known in the previous task while they are proposed using a piece-wise linear regression strategy in the latter. Therefore, the prior for the parameters are the same as the one discussed above. Given the trajectory $X = \{(x_n, y_n)\}$ where a resistant and a sensitive population are assumed to co-exist, the parameters of interest will the two growth rates $\rho_s$ and $\rho_r$, the size of the sensitive population $n_s$ at the instant of time of the first observation and the instant of time in which the resistant population is born $t_r$. The priors for those values are

$$
\begin{aligned}
\rho_s &\sim \text{HalfNormal}(0, 1) \\
\rho_r &\sim \text{HalfNormal}(0, 1) \\
t_r &\sim \text{TruncatedNormal}(x_1, 1, -\infty, x_N) \\
t_s &\sim \text{TruncatedNormal}(x_N, 1, x_1, \infty)
\end{aligned}
$$

Essentially, we choose a standard normal distribution for the two growth rates, a Normal distribution bounded from above by $x_N$ for $t_r$, and a Normal distibution bounded from below by $x_1$ for $t_s$. The full joint is therefore

$$
\begin{aligned}
p(\boldsymbol{X}, \boldsymbol{\Theta}) = & \left( \prod_{n=1}^{N} \text{Poisson}(y_n \mid \mathbb{E}[y(x_n) \mid \boldsymbol{\Theta}]) \right) \\
& \mathcal{N}(\rho_s \mid 0, 1) \cdot \mathcal{N}(\rho_r \mid 0, 1) \\
& \mathcal{N}_{trunc}(t_s \mid x_1, 1, -\infty, x_N) \cdot \mathcal{N}_{trunc}(t_s \mid x_N, 1, x_1, \infty)
\end{aligned}
\tag{34}
$$

##### C.5. Model validation with simulations

We first tested how observational noise affects the inference of growth rates and $t_0$ by fixing the initial population size $y(0) = 1000$ and simulating exponential growth processes with 1-3 time windows. Observed data points ranged from 10 to 20, with noise $\epsilon_n$ added to each true observation $y_n$. This noise was drawn from a Normal distribution with mean 0 and a standard deviation proportional to $f_{noise} y_n$. The noise factors $f_{noise}$ ranged between 0 and 0.2 (Fig.1b). We also explored the impact of noisy change-points by defining $t_{i,noisy} = k t_i$, with $k$ as a noise factor between 0.9 and 1.1. The number of time windows ranged from 2 to 4, while observational noise was fixed at 0, and 20 data points were used (Extended Data Fig.2a). Next, we examined how population size affects inference accuracy by simulating growth processes for different initial populations $y(0)$, ranging from 10 to $10^6$. As expected, inference error decreased as the population size increased (Extended Data Fig.2b). We then tested biPOD's performance in distinguishing between logistic and exponential growth. For this, we simulated growth processes with 10 data points and tracked the carrying capacity ratio (the ratio between the largest observation $y_i$ and the true carrying capacity) in logistic growth. Since logistic growth is often indistinguishable from exponential growth in its early stages, we divided our performance analysis between high and low ratio (Extended Data Fig.2c). We compared the two inference algorithms in biPOD, MCMC and Variational Inference, in terms of both accuracy and speed. Simulations were performed with increasing numbers of data points (from 10 to 150) and varying time windows (from 1 to 3). As expected, the inference time increased linearly with the number of data points and time windows for both algorithms (Extended Data Fig.2d). Variational Inference was up to 100x faster than MCMC (Extended Data Fig.2e), though MCMC had slightly better accuracy, particularly at low observation densities (Extended Data Fig.2f). We further tested biPOD's ability to infer change-points in the presence of noise by simulating exponential processes with high observational noise levels ($f_{noise} \geq 0.15$). Errors in change-point inference surpassed 10% when the number of time windows exceeded 2 (Fig.1c). Finally, to test biPOD's ability to model a convoluted exponential process involving two populations (one growing, one shrinking), we simulated a scenario with a resistant population potentially existing before the first observation. As shown in Fig.1d, biPOD could reliably detect the resistant population as long as observations were made over a sufficiently long period. Longer observation times also reduced error in the inference of parameters for the resistant population ($t_r$ and $\rho_r$), whereas shorter observations still yielded accurate results for the sensitive population (Extended Data Fig.2g). Additionally, goodness of fits can also be assessed using standard Bayesian diagnostics like trace plots and posterior predictive checks[14].

#### C.6. Model validation with xenograft data

The input data consists of weekly measured tumour volumes ($mm^3$) of patient derived xenografts (PDX) mice. The cohort consists of $n = 4$ patients and of $n = 35$ mice. Of those, 18 mice underwent carboplatin treatment while 17 were used as control. The x-axis, originally measured in days and with $x = 0$ indicating the start of treatment, was converted in weeks. Additionally, the data has been pre-processed by removing a couple of outliers from the original data. More specifically, the maximum tumor volume for mouse 530, and the second-highest tumor volume for mouse 541 have been removed. biPOD was used to fit the dynamics of a given PDX mouse only if the time-series had more than 4 observations for $x \geq 0$ (with less observations detecting eventual changepoints is not feasible). Therefore, the used cohort consisted of 32 mice ($n = 15$ control, and $n = 17$ treated mice). The data used are available from Sauer et al.[1].

#### C.7. Case studies

##### *RAS-resistance in colorectal cancers*

##### *Data*

The data consists of carcinoembryonic antigen (CEA) measurements over time, where CEA can be considered a surrogate for tumour burden. The cohort consists of $n = 45$ patients which underwent cetuximab treatment. The response of each patient to treatment is also classified following RECIST[15] criteria between progression, stable disease, and responder. The time series of each patient includes not only the treatment period but also measurements taken before and/or after the treatment period. The median number of observations per patient is $n = 22$ (min $=$ 12,max $=$ 54), while the median length of the time series was of 25 months (min $=$ 7 months, max $=$ 107 months). The time scale on the x-axis is measured in weeks, while CEA is measured in $\mu$mol/l. For the breakpoints detection part, we considered only a relevant timeframe including one year before and after each patient's treatment window. The data are available from Khan et al[2].

##### *Timing of resistant and sensitive populations*

Given two population, one sensitive and one resistant, it is possible to compute the time $t_k$ in which the resistant clone reaches a size which is $k$ times the size of the sensitive population. Assuming that the size of the two populations can be described by an exponential growth model, i.e.

$$N_r(t) = N_r(0)\exp(\rho_r t) \tag{35}$$
$$N_s(t) = N_s(0)\exp(\rho_s t) \tag{36}$$

then it become straightforward to solve for the equation $N_r(t) = kN_s(t)$ and find the time $t_k$ that makes the equation true. This time is given by

$$t_k = \frac{\ln \frac{N_r(0)}{kN_s(0)}}{\rho_s - \rho_r} \tag{37}$$

The equation can therefore be used to estimate, for example, when the resistant populations reaches the same size as the sensitive population ($k = 1$) or when it doubles the size of the sensitive population ($k = 2$). Additionally, given the fact that the growth rates are estimated with a certain uncertainty, it is possible to incorporate the standard deviation of the growth rate estimates to achieve a probabilistic prediction. This means that the denominator in 37 becomes a random variable $\Delta\rho = \rho_s - \rho_r$ that is distirbuted as the difference of the two growth rates posterior.

Therefore a Monte Carlo schema can be used to account for the uncertainty in the growth rate estimates and provide a more robust understanding of the time dynamics for the resistant clone's expansion.

##### Patients stratification and MRD testing in lung cancers

##### Data

The input data consists of longitudinal observations of variant allele frequency (VAF) for 200 patient-specific mutations linked to the tumour. Given that the tumour size is expected to correlate with the amount of dead tumour cells shed into the bloodstream, the VAF is used as proxy for the tumour burden. Each observation comes also with a p-value for an MRD test based on the sequencing error rates, indicating whether the observed mutated reads are true or false positives. For fitting the VAF dynamics, we considered only patients with at least 2 observations atfer surgery with positive VAF. After pre-preprocessing, the cohort comprised 101 patients. Of those, 4 died, 49 relapsed, 16 developed a second primary tumour, and 32 underwent remission. For relapsing patients, we considered only observations before relapse, gathering from 2 to 13 observations per patient (median of 5), over an observation period from 49 to 1,651 days (median of 640 days). To classifify patients into MRD negative, early positive and late posiitve we used a p-value of 0.01 as significance level for the reported MRD test. The data is available from Abbosh et al[3].

##### Growth rates clustering

The inferred growth rates $\rho$ for each one of the 101 patients have been clustered in three different groups using the R package mclust[16]. The clustering procedure is based on a finite Gaussian mixture models in which an expectation maximization algorithm fits the parameter for different numbers of mixture components $G$ and then the optimal number of clusters is selected according to Bayesian Infromation Criterion (BIC)[17]. Then, given the inferred parameters of the mixture of gaussians, we retrieved the separation boundaries, i.e. the values where the posterior probabilities of two (or more) clusters are equal, in order to separate patients bewteen clusters. In our case, we found 3 to be the optimal number of cluster (Supplementary Fig.S7b) and the thresholds that separated the three groups were $\rho_{lower} = 0.011$ and $\rho_{upper} = 0.099$. We validated the obtained number of clusters and thresholds using bootstrap. We repeated the clustering for 1000 iterations using at each iteration a subsample of 89 values (i.e. 90% of the original dataset). In 981 cases out of 1000 (i.e. 98.1% of the time), the optimal number of clusters was 3. On the other hand, by inspecting the distibution of the inferred thresholds (Supplementary Fig.S7a) it is possible to observe that the lower threshold has a unimodel distribution centered around $\rho_{lower}$ while the upper threshold has a bimodal distribution, one centered around $\rho_{upper}$ and one centered around $\rho = 0.2$. Since the higher value is obtained only 106 times out of 981 (i.e. ˜10% of the time), we consider $\rho_{upper}$ to be a more robust choice.

##### Contingency analysis

We investigated whether the three clusters induced a stratification associated with any variable of interest. To do so, for each variable of interest we created a contingency table containing the multivariate frequency distirbution of the given variable and the clusters assignment. Then, we tested the hypothesis of independence of the observed frequencies using a chi-squared test and the obtained p-values were corrected using the Benjamini–Hochberg procedure[18]. First, we selected 12 genes (i.e. TP53, KRAS, NF1, KMT2D, CREBBP, KEAP1, ARID1A, EGFR, PIK3CA, STK11, ATM) including crucial NSCLC genes and the most frequently mutated genes in the cohort. Interestingly, no mutations were associated with the three groups since all the adjusted p-values were above the significance threshold $\alpha = 0.05$. All the results are reported in Supplementary Table 1, along with the proportion of patients in each cluster with the mutation. Second, we analysed 5 variables of interest, namely TNM stage, smoking, histology, gender, and age. For continuous variables (i.e. age and smoking) we stratified the patients in three groups based on the first and third quantile. Also in this case, no significant associations were found. All the results are reported in Supplementary Table 2.

##### Simulations

We tested how fast biPOD can correclty detect patients with a high growth rate $\rho$. To do so we simulated patient trajectories and classified them based on relapse dynamics. A single patient is simulated as follows. First, a growth rate $\rho$ is sampled from the three component gaussian mixture clusters already discussed in detail previously. Second, given the sampled $\rho$, an exponentially growing population is simulated. Third, the integer counts of the simulated population are converted into probability of observing mutating read assuming that each cell increases the probability by $3 \times 10^{-6}$ (which corresponds to an expected VAF of $3 \times 10^{-6}$ for a population of a single cell). Additionally, noise sampled from a beta distribution is added to this probability. Finally, the VAF is simulated using a binomial sampling with an assumed coverage of $2 \times 10^5$. To simulate how a real-world clinical setting might work when monitoring a patient's disease progression, we classifify each patient starting with the first two observations and slowly incorporating new data points. As more data comes in, the model refines the inference of $\rho$ and the probability that a patient is "Fast relapse" is computed as $P(\rho \geq \rho_{upper})$ using the posterior distribution of the growth rate. The simulations were conducted for different time intervals bewteen observations, in order to mimic the profiling of patients evry $n = 1, 2, 4$ weeks.

##### MRD test

We developed an alternative MRD test using biPOD, based on the assumption that the number of observed variant reads (NV) originates either from technical noise alone or from a combination of technical noise and the presence of the tumor. To achieve this, we first modeled the technical noise. Using the MRD results reported in the original data and a p-value threshold of 0.01, we selected $n = 54$ patients who never had a positive MRD test (i.e., no p-value $\leq 0.01$). From this group, we further subsampled to obtain a final set of $n = 27$ MRD-negative patients. For each of these patients, we used biPOD to regress their variant allele frequency (VAF) at the time of surgery and fit a beta distribution to model the noise, resulting in the noise distribution Beta($\alpha_{\text{noise}}, \beta_{\text{noise}}$). Under the null hypothesis $H_0$, which assumes only technical noise is present, the number of variant reads (NV), given the total number of reads (DP), follows a Beta-Binomial($DP, \alpha_{\text{noise}}, \beta_{\text{noise}}$) distribution. Hence

$$H_0 : NV \sim \text{Beta-Binomial}(DP, \alpha_{\text{noise}}, \beta_{\text{noise}}) \tag{38}$$

We then calculate a bayesian p-value to quantify how much the observed reads deviate from the null distribution

$$p = P(NV \geq NV_{obs} \mid DP, \alpha_{\text{noise}}, \beta_{\text{noise}}) = \sum_{x=NV_{obs}}^{DP} \text{Beta-Binomial}(x \mid DP, \alpha_{\text{noise}}, \beta_{\text{noise}}) \tag{39}$$

If the p-value is small enough, we reject $H_0$ in favor of the alternative hypothesis, considering the test positive.

##### Growth patterns and adaptive monitoring in leukemia

###### Data

The input data consists of lonigitudinal measurements of white blood counts (WBC) for patients with chronic lymphocytic leukemia (CLL). The cohort consists of $n = 147$, of whom 79 underwent treatment and 68 experienced natural evolution. The WBC were measured at irregular intervals (every $k$ months per patient). The median number of observations per patient is $n = 7$ (min $= 3$,max $= 20$), while the median length of the time series was of 60 months (min = 2 months, max = 216 months). The dataset contains also clinical variables commonly used in CLL stratifications, such as trisomy of chromosome 12 (Tri12), deletions in chromosomes 11, 13, and 17 (Del11, Del13, Del17), TP53 mutation status, IGHV mutation status, and CD49d expression. The dynamics of each patient has been fit using one year as unit of time for the x-axis, considering only observations prior to an eventual treatments, and, if a patient had 3 or more observations, using a model selection algorithm to classify the growth patter between exponential and logistic.

##### Contingency analysis

We investigated whether the treatment status (treated vs. untreated) was associated with any variable of interest. For each variable, we created a contingency table and tested the hypothesis of independence between the observed frequencies using a chi-squared test. The resulting p-values were corrected using the Benjamini–Hochberg procedure[18]. First, we analyzed six clinical variables of interest: IGHV mutational status, CD49d expression, and the presence of four copy number alterations (Tri12, Del13, Del11, Del17). Interestingly, only IGHV mutational status showed a significant p-value, with 54% of treated patients having unmutated IGHV (compared to only 18% of untreated patients). This result is expected, as patients with unmutated IGHV generally have a worse prognosis[19]. The full results are presented in Supplementary Table 3. Second, we examined two main parameters derived from biPOD: growth rate $\rho$ and growth pattern. The growth rate $\rho$ exhibited a significant p-value. We stratified patients based on the first and third quartiles of the growth rate distribution, referred to as $\rho_{\text{q1}}$ and $\rho_{\text{q3}}$, respectively. We found that 37.2% of treated patients had a growth rate $\geq \rho_{\text{q3}}$, while this was true for only 11.9% of untreated patients. Conversely, 16.6% of treated patients had a growth rate $\leq \rho_{\text{q1}}$, compared to 35.8% of untreated patients. The results are presented in Supplementary Table 4.

##### *Model training*

We used a gradient boosting algorithm (xgboost) to build a first model that predicts the probability that a patient will undergo treatment and a second model that predicts whether a patient will undergo treatment in in the following 18 months. Along with the clinical variables mentioned in the contingency analysis previously described were used along with

- Pattern: represents the tumor growth pattern, categorized as Exponential or Logistic.
- IGHV: indicates the mutation status of the gene IGHV, which is either mutated or unmutated.
- WBC: represents the white blood cell count at the most recent time point. High WBC counts can indicate disease progression.
- $WBC_0$: represents the white blood cell count at the first time point, serving as a baseline measure.
- $\hat{\rho}$: represents a smoothed growth rate of the disease over time, summarizing the growth trend up to a given observation point

The last variable represents the average growth rate across a subset of observations. Therefore, given a patient with $n$ observations, $\hat{\rho}$ would be computed as $\sum_i (\rho_i \times i)/\sum_i i$ with $i$ ranging from 1 up to $n$. The dataset was then split into two parts: 70% of the patients were used tothe model, while the remaining 30% were used to test it. In the first model, predictions were considered positive (indicating that the patient is likely to undergo treatment) if the predicted probability was above 0.6, while a probability of 0.5 was sufficient to deem an observation as positive in the second step. The model's performance was evaluated using training and testing accuracy, comparing predicted outcomes to actual treatment statuses and are reported in Supplementary Tables 5-6.

##### *Software and code*

Analysis was performed in the R statistical environment (v.4.4.1)[20]. For general data preparation and manipulation we used the R packages tidyverse (v.2.0.0)[21], and readxl (v.1.4.3)[22]. For performance of binary classification, accuracy was computed as the number of correctly classified results over the total number of results; sensitivity (i.e. TPR) was computed as the number of true-positive results divided by all the positive results; false positive rate (i.e. FPR) was computed as the number of false-positive results divided by all the negative results. For additional statistical analysis we used the R packages MASS (v.7.3.61)[23], mclust (v.6.1.1)[16], and survival (v.3.7.0)[24]. All the plots were produces using the R packages ggplot2 (v.3.5.1)[25], ggpubr (v.0.6.0)[26], patchwork (v.1.3.0)[27], ggsurvfit (v.1.1.0)[28], and ComplexHeatmap (v.2.20.0)[29].
